# Membrane contact probability: an essential and predictive character for the structural and functional studies of membrane proteins

**DOI:** 10.1101/2021.01.17.426988

**Authors:** Lei Wang, Jiangguo Zhang, Dali Wang, Chen Song

**Affiliations:** Center for Quantitative Biology, Academy for Advanced Interdisciplinary studies, Peking University, Beijing 100871, China; School of Life Sciences, Peking University, Beijing 100871, China; Peking-Tsinghua Center for Life Sciences, Academy for Advanced Interdisciplinary Studies, Peking University, Beijing 100871, China

## Abstract

One of the unique traits of membrane proteins is that a significant fraction of their hydrophobic amino acids is exposed to the hydrophobic core of lipid bilayers rather than being embedded in the protein interior, which is often not explicitly considered in the protein structure and function predictions. Here, we propose a characteristic and predictive quantity, the membrane contact probability (MCP), to describe the likelihood of the amino acids of a given sequence being in direct contact with the acyl chains of lipid molecules. We show that MCP is complementary to solvent accessibility in characterizing the outer surface of membrane proteins, and it can be predicted for any given sequence with a machine learning-based method by utilizing a training dataset extracted from MemProtMD, a database generated from molecular dynamics simulations for the membrane proteins with a known structure. As the first of many potential applications, we demonstrate that MCP can be used to systematically improve the prediction precision of the protein contact maps and structures.

## Introduction

Proteins with various amino acid sequences are folded into specific structures with unique functions [1]. The relationship between the sequence, structure, and function of proteins has been extensively studied for decades. To characterize the structural and functional features of various proteins, researchers defined some essential and predictive properties, such as the secondary structure (SS) and solvent accessibility (SA) [2–5], which are widely used in the analysis and prediction of the structure and function of proteins [6–8].

Membrane proteins represent a large subgroup of proteins, which are responsible for the substance transport and signal transduction across cell membranes. Currently, over 50% of the known drugs target membrane proteins [9]. Therefore, studying the structures of membrane proteins is of high interest. The recent progress of Cryo-EM has facilitated the determination of many membrane protein structures [10], but these are still only a small fraction of the known sequences that encode into membrane proteins [11, 12], rendering the structural prediction of membrane proteins highly desirable.

Recent developments in deep learning have improved the protein structure prediction accuracy to a large extent [7, 8, 13–15]. However, researchers have typically focused mostly on soluble proteins, which are easier for structural determination; and therefore, a large structural dataset is available. For those soluble proteins, SS and SA can be routinely predicted to characterize their local structural features [2, 3, 5], which are then extensively used as inputs for contact map (CM) and structure predictions [6–8, 16, 17]. Although membrane proteins share some common features with soluble proteins, and both SS and SA are essential and applicable to membrane proteins too, membrane proteins are distinct from soluble proteins in the sense that a significant part of their amino acids on the outer surface are in direct contact with the hydrophobic acyl chains of lipid molecules. This means that a large fraction of the outer surface of membrane proteins is covered by hydrophobic residues, which would be considered not ‘solvent exposed’ and therefore embedded in the ‘interior,’ if one predicts in the same way as for soluble proteins. Strikingly, this remarkable difference between membrane proteins and soluble proteins has been rarely considered in the structural predictions to date, probably due to the absence of a quantitative prediction method for this lipid-exposing property from sequences. We believe that the membrane-contacting feature of a membrane protein should be explicitly considered and utilized with the same weight as SA in the structure and function studies of membrane proteins.

The SA of a given protein can be predicted with deep learning-based models to high accuracy [18]. The dataset of SA for the training was generated by analyzing the outer surface of proteins with a known structure via rolling a water-size sphere over the surface [19, 20]. In principle, one can use similar protocols to generate lipid accessibility datasets of membrane proteins with a known structure. In fact, there have been such attempts [21–24] to calculate the relative or absolutely accessible surface area for lipids. For example, mp_lipid_acc [25], part of the Rosetta software suite, can identity lipid-accessible surface area or lipid accessible residues based on known 𝛼-helix and 𝛽-sheet membrane protein structures. However, in most of the above cases, researchers used a highly simplified and shape-fixed membrane slab to define the lipid environment for the calculation of the lipid accessibility of residues, which was merely a spacial property. Based on these calculations, researchers also tried to predict lipid accessibility from membrane protein sequences [26–29], but the dataset of only about 100 proteins seemed too small to get a satisfactory prediction. Therefore, there has not been a well-defined and widely accepted quantity to describe the membrane-contacting properties of amino acids of a sequence, although there are multiple methods to predict the transmembrane topology of a membrane protein [30–34] (Table S1).

To account for the hydrophobic-surface feature of membrane proteins, we propose a new characteristic quantity in this work, the membrane contact probability (MCP), to describe the likelihood of direct contact between the protein amino acids and the hydrophobic acyl chains of lipid molecules. We show that we can use a deep learning-based method, DCRNN [35], to predict the MCP to a good accuracy for a given protein sequence, based on the highly informative data from the MemProtMD database [36, 37]. We integrated MCP into the recently developed ResNet-based contact map predictor [6, 17, 38], and the results showed a consistent and significant improvement of the contact map and structure prediction. Therefore, we propose that the MCP is an essential property of membrane proteins, which can be predicted and used for broad applications such as the contact map and structure prediction of (membrane) proteins.

## Results

### MCP can be predicted with good accuracy for membrane proteins

We consider an amino acid to be in direct contact with the hydrophobic core of membranes if its 𝛼 carbon atom is within 6 Å of the lipid acyl chain carbon atoms, and define the MCP to be the fraction-of-time probability of a certain amino acid in direct contact with the hydrophobic acyl chains of a lipid bilayer. The MCP is difficult to obtain directly from experimental measurements, but it can be easily calculated from molecular dynamics (MD) simulations of membrane proteins. Stansfeld et al. performed systematic coarse-grained (CG) MD simulations for all of the membrane proteins with a known structure, and the simulation and analysis results were deposited into a database named MemProtMD [36, 37]. Based on this pioneering work, we extracted the MCP information of all of the simulated membrane proteins as the training dataset (termed ‘MCP-Large’) for our DCRNN model (please refer to the “Methods” section and Fig. 1 for details). With this model, we were able to predict the MCP for a given sequence to good accuracy. Please note that, while the training dataset was obtained from MD simulations for the membrane proteins with a known structure, our prediction model does not require any structural information. A protein sequence is all that is needed for the MCP prediction.

**Figure 1.**
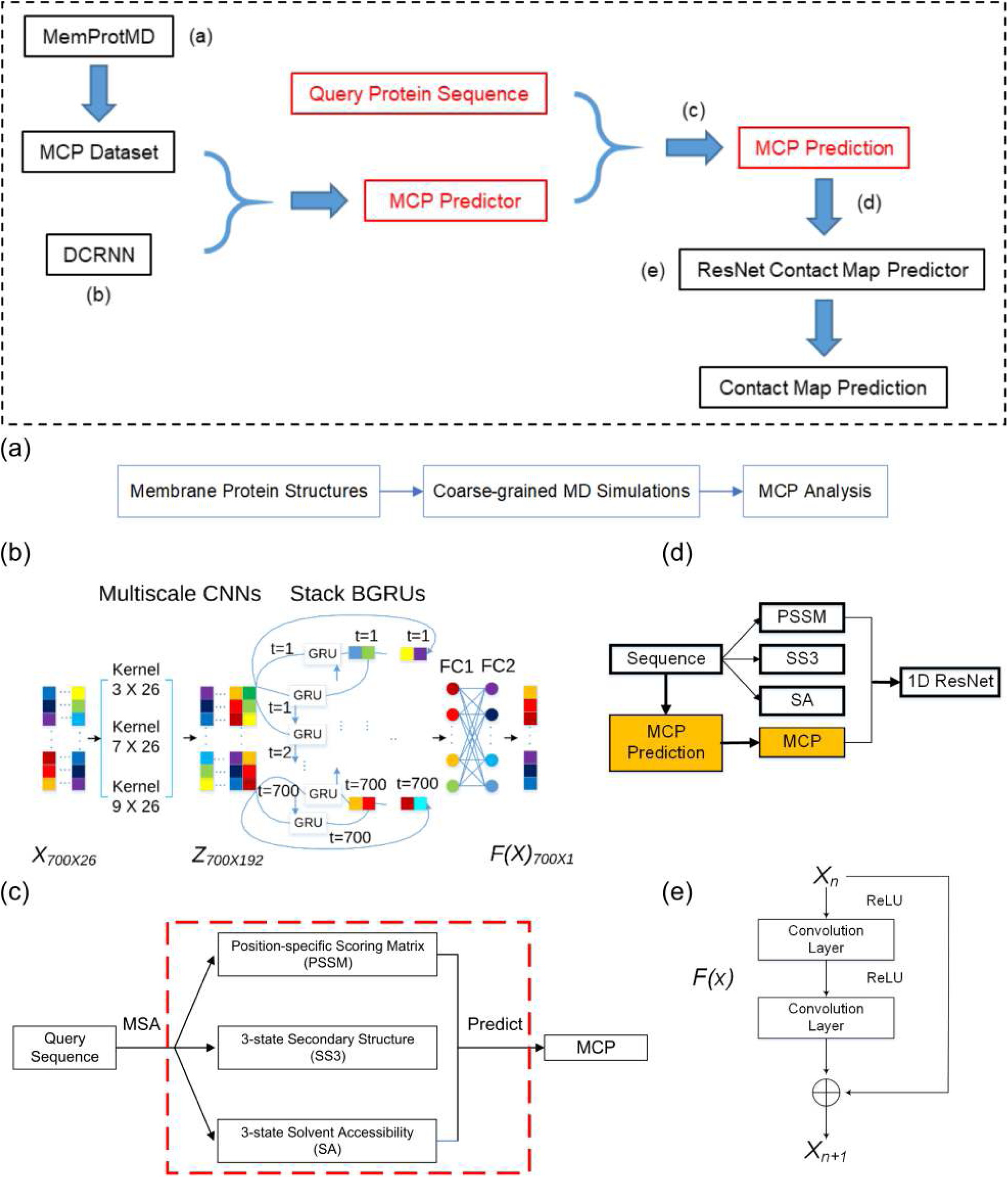
The method schemes. The overall scheme is presented in the black dashed rectangular in the top panel, and the key steps are described in: (a) Extraction of MCP from the MemProtMD database. (b) The DCRNN model for the MCP prediction. (c) The process of MCP prediction from a query sequence. (d) The method of MCP incorporated into the ResNet model. The MCP was used as a 1D input in the same way as SA. (e) The unit of the ResNet model. Each unit of the ResNet model contains two convolution layers and two activation layers.

As shown in Table 1, the overall Pearson Correlation Coefficient (PCC) between our MCP prediction and the MD observation (obtained from MemProtMD, which can be viewed as the ground truth here) of the studied membrane proteins reached 0.77 for the training set, 0.76 for the validation set, and 0.77 for the test set, respectively. Ideally, it would be better to use datasets of lower sequence similarity (< 25%). However, due to the limited membrane protein data available, using a low-sequence-similarity dataset would lead to the under-training problem. As a test, we trained an MCP predictor with a small dataset containing less-redundant sequences, in which the sequence identity between the training set and test set was less than 40% (termed ‘MCP-Small’). The prediction accuracy was still satisfactory with an overall PCC of about 0.65 (Table S2), but significantly dropped compared to with the larger dataset. Therefore, in this work, we utilized the MCP predictor trained by the MCP-Large dataset, which made the method less *De Novo* but generating more accurate predictions. As a comparison, the highest prediction accuracy of SA is around 80% at present [18], after many years of development with a much larger training dataset.

**Table 1:**
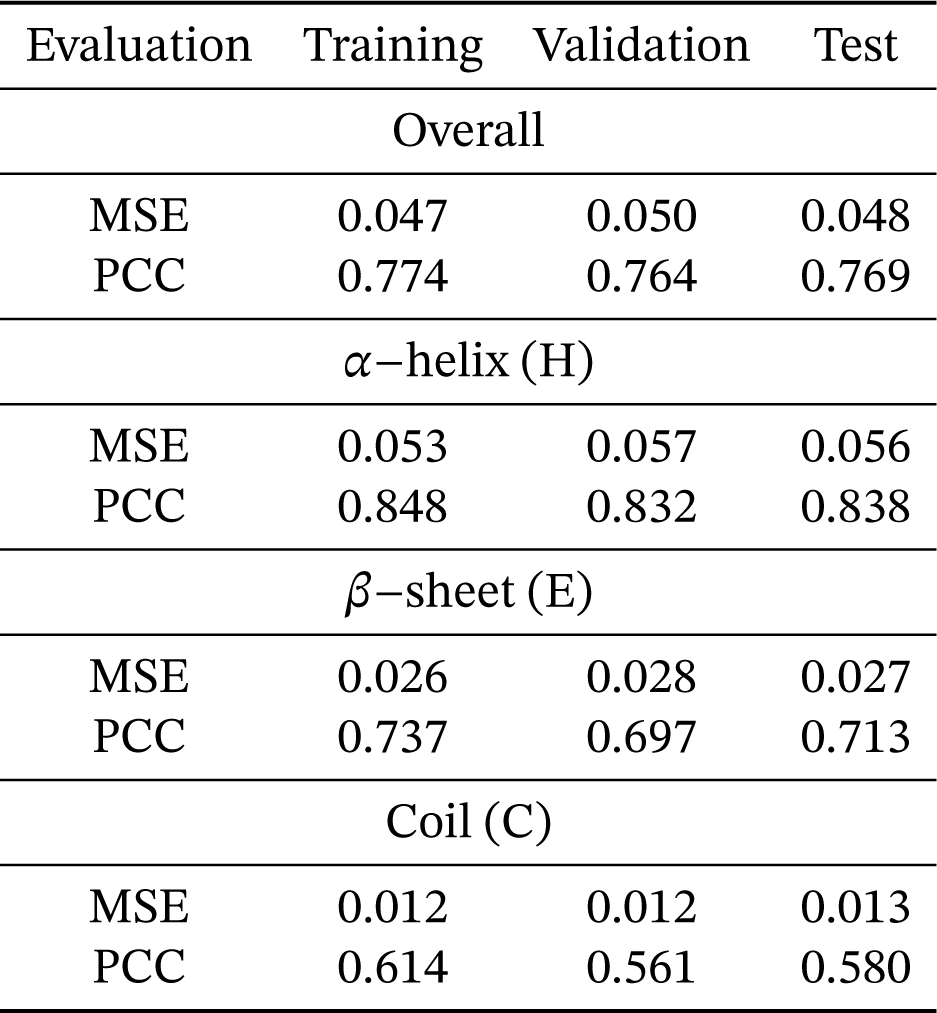
The performance of our MCP predictor.

| Evaluation | Training | Validation | Test |
| --- | --- | --- | --- |
| Overall |  |  |  |
| MSE | 0.047 | 0.050 | 0.048 |
| PCC | 0.774 | 0.764 | 0.769 |
| α-helix (H) | | | |
| MSE | 0.053 | 0.057 | 0.056 |
| PCC | 0.848 | 0.832 | 0.838 |
| β-sheet (E) | | | |
| MSE | 0.026 | 0.028 | 0.027 |
| PCC | 0.737 | 0.697 | 0.713 |
| Coil (C) |  |  |  |
| MSE | 0.012 | 0.012 | 0.013 |
| PCC | 0.614 | 0.561 | 0.580 |

We analyzed the prediction performance for the 𝛼-helix and 𝛽-sheet structures, two of the most common secondary structures of transmembrane proteins. The 𝛼/𝛽 PCC was calculated according to the MCP values of each residue within the corresponding secondary structure types. As shown in Table 1, the prediction accuracy is better for the 𝛼-helix structures (PCC=0.84 for the test) than for the 𝛽-sheet structures (PCC=0.71), probably because we had a larger dataset of 𝛼-helix structures in the training set (54%), compared to that of the 𝛽-sheet structures (23%). The coil structures were the most difficult ones (PCC=0.58), as these are the most flexible and less abundant (23%) structures in the datasets. Further analysis showed that the predictor performs more reliably for multi-pass 𝛼-helix proteins than single-pass ones (Table S3), which may also be related to the amount of proteins of different classes in the datasets (Table S4).

We picked an 𝛼-helix and a 𝛽-sheet membrane proteins from the test set as representative cases to investigate the prediction details (Fig. 2). As can be seen, most of the hydrophobic lipid-contacting amino acids were successfully predicted (Fig. 2a&b), with the overall prediction PCCs of 0.76 and 0.71 for the 𝛼-helix and 𝛽-sheet membrane proteins, respectively. We mapped the MCP onto the surface of the protein structures, and it is evident that the distribution of high MCP values is indeed in the transmembrane region, consistent with the MD observations (Fig. 2c-h). For the 𝛼-helix protein (5aym), most of the residues contacting the hydrophobic core of the lipid bilayer were identified (Fig. 2a&g). For the 𝛽-sheet protein (4e1t), there were some lipid-contacting residues missing in our prediction, as shown with the dashed circles in Fig. 2h. However, the predicted high-MCP residues are mostly in the transmembrane region and on the outer surface (Fig. S1), indicating that the MCP prediction can reach a satisfactory accuracy in characterizing the membrane hydrophobic core-contacting residues. The results obtained from the MCP predictor trained by the MCP-Small dataset were similar (Fig. S2).

**Figure 2.**
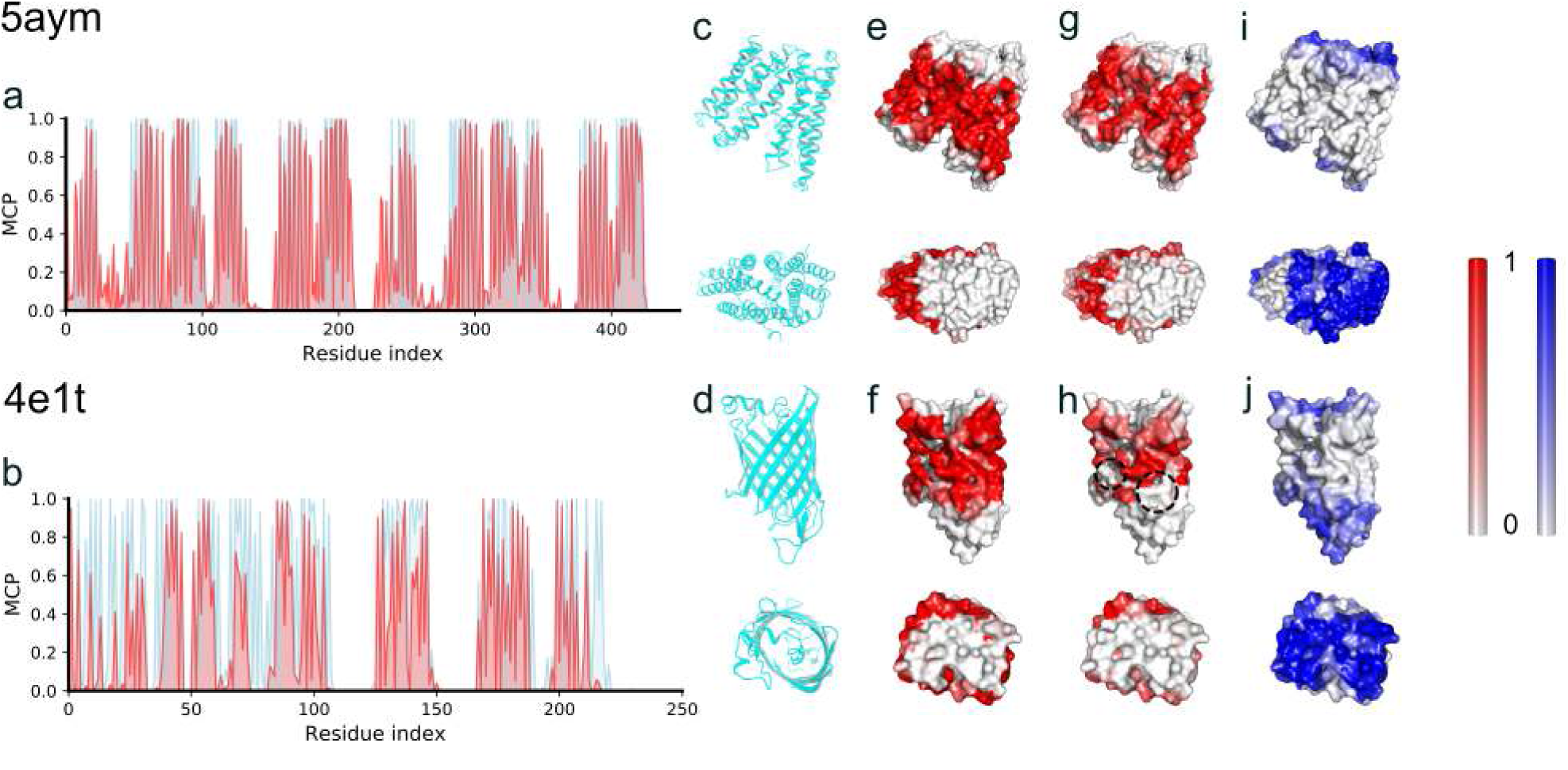
Membrane contact probability (MCP) of two representative membrane proteins. (a-b) Comparison between the MD observation (cyan) and the prediction (red) of the MCPs. (c-d), Side and top views of the two representative proteins. (e-f), The outer surface of the representative membrane proteins, colored according to the MCP values obtained from MD simulations. (g-h), Similar to (e-f), but colored according to the predicted MCP values. (i-j), Similar to (e-f), but colored according to the predicted SA values by RaptorX. The PDB ID of the 𝛼-helix membrane protein was 5aym [88], representing a crystal structure of a bacterial homologue of iron transporter ferroportin in the outward-facing state with soaked iron with more than 10 transmembrane helices and 440 residues. The PDB ID of the 𝛽- sheet membrane protein was 4e1t [89], an X-ray crystal structure of the transmembrane beta-domain from invasion from *Yersinia pseudotuberculosis* with 245 residues.

### MCP is complementary to SA and provides important structural information for membrane proteins

For the two membrane proteins discussed above, we also predicted the SA from their sequences [4] and colored the protein according to the SA values of each amino acid (Fig. 2i-j). As can be seen, the SA prediction was reasonable, but the high-SA amino acids do not cover the full surface of the two membrane proteins, highlighting the deficiency of only using SA to describe the outer surface of a membrane protein. In fact, the high-SA region (blue-colored surface) and high-MCP region (red-colored surface) are complementary to each other on the outer surfaces of the two proteins, and together they give a complete description of the outer surface of the membrane protein structures (Fig. 2e-j).

To give a more comprehensive analysis, we plotted the predicted MCP and SA of two datasets in Fig. 3. Fig. 3a was generated from the test dataset pdb25-test-500 [6], which contained 25 membrane proteins and 302 soluble proteins after removing the sequence redundancy with respect to our training set (327 test proteins in total). Therefore, some outer-surface amino acids in the dataset should be exposed to water molecules, and some to lipid molecules. If one does not consider MCP and uses only the SA to predict and characterize the amino acids of the dataset, the residues would be considered to be either exposed to water or embedded in the interior, which is usually the logic of most of the SA predictors. However, from Fig. 3a, we can tell that a significant number of amino acids, especially those in region II, are neither exposed to water molecules nor embedded in the interior of proteins. They are exposed to lipid molecules. Therefore, this plot highlighted the necessity of taking the MCP into account while considering the outward-facing amino acids of a membrane protein.

**Figure 3.**
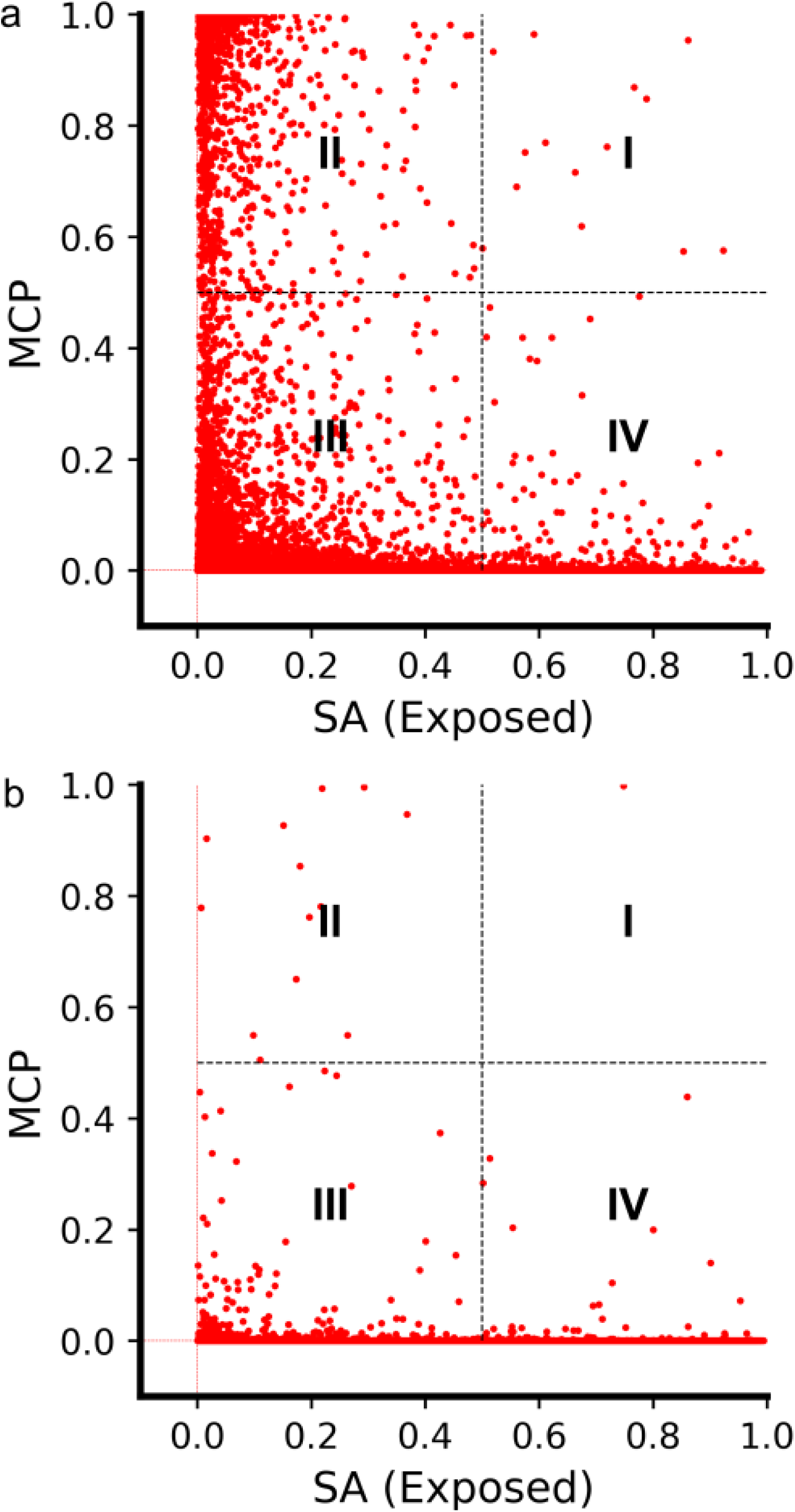
The 2D plot of the complementary MCP and SA predicting the likely location of amino acids in a protein, for the 327-protein dataset (a) and 102 Pfam protein dataset (b), respectively. The figures are roughly divided into four regions: (I) Both the MCP and SA values are larger than 0.5, indicating the amino acids in this region are likely to be exposed to both water and lipid molecules; (II) MCP > 0.5 and SA < 0.5, meaning the amino acids in this region are more likely to be exposed to hydrophobic lipid molecules than to water molecules; (III) MCP < 0.5 and SA < 0.5, meaning the amino acids are not likely to be exposed to either lipid or water molecules, so they are probably embedded in the interior of proteins; and (IV) MCP < 0.5 and SA > 0.5, meaning the amino acids are more likely to be exposed to water molecules than to the lipid bilayer.

Therefore, the amino acids located in regions II, III, and IV are likely to be those exposed to the hydrophobic core of the lipid bilayers, those embedded in the protein interior, and those exposed to water molecules, respectively. At a first glance, region I seems puzzling: are there residues that have a high probability to be exposed to both water and lipid molecules? Further analysis using the OPM server [39] revealed that most of the amino acids located in region I actually represent these residues of the membrane protein sitting at the water-membrane interface (Fig. S3). It should be noted that only a very small fraction of the amino acids was found to be located in this region under the cutoff of 0.5 for this dataset. To be specific, the amino acids in region I only account for 0.018% (14/79796) of the total amount of the amino acids in the dataset.

### The MCP predictor performed well for non-transmembrane proteins too

We also examined how the predictor performs for non-transmembrane proteins. We picked another two representative proteins from the test dataset: one is a half-membrane-embedded protein, and the other is a soluble protein. As can be seen in Fig. 4, most of the membrane-contacting residues of the half-membrane-embedded protein were correctly predicted, and none of the soluble protein residues were predicted to be membrane-contacting. These results indicate that our predictor may have a good performance for non-transmembrane proteins too.

**Figure 4.**
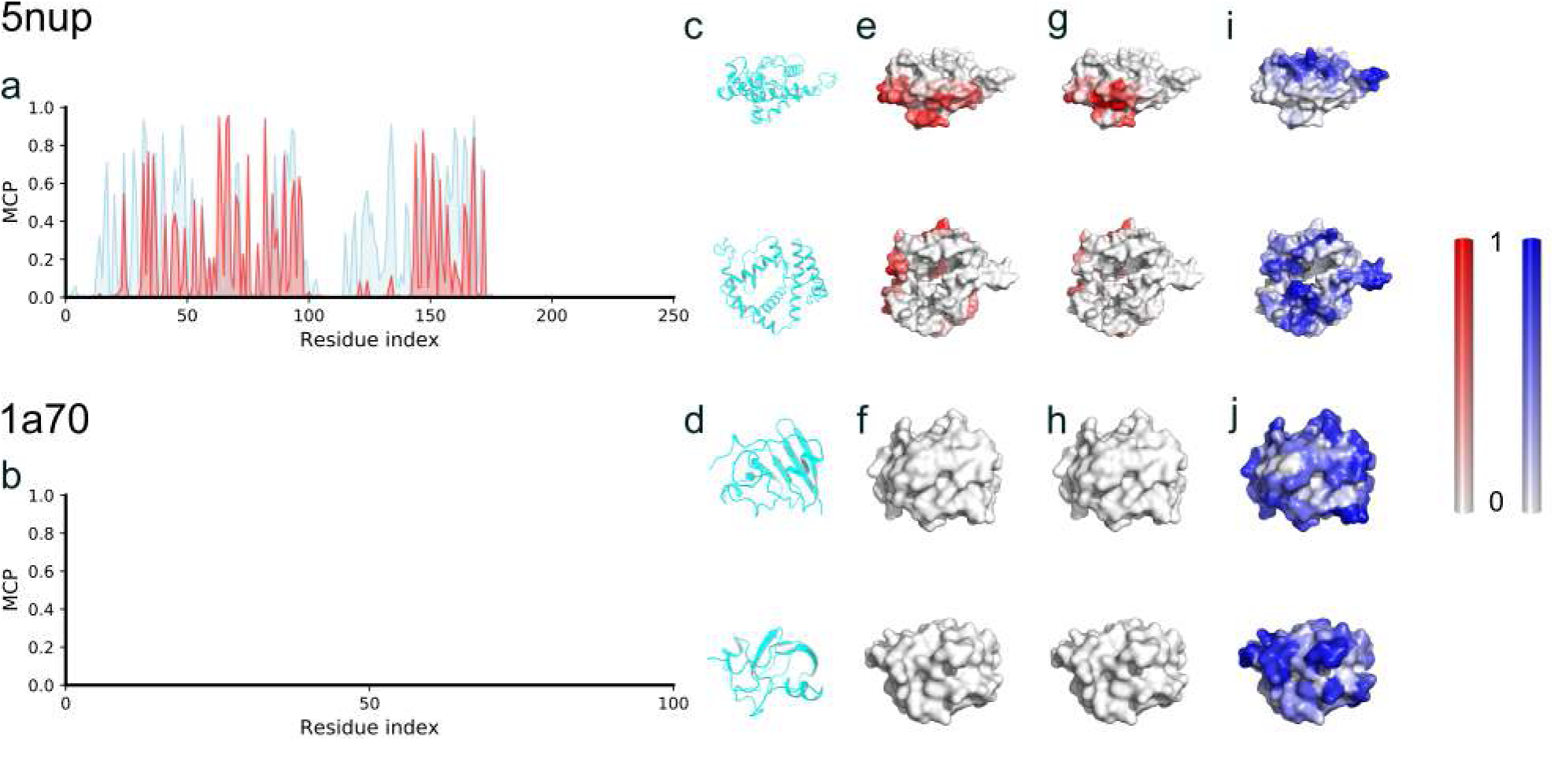
Membrane contact probability (MCP) of two representative non-transmembrane proteins. (a-b) Comparison between the MD observation (cyan) and the prediction (red) of the MCPs. (c-d), Side and top views of the two representative proteins. (e-f), The outer surface of the representative proteins, colored according to the MCP values obtained from MD simulations. (g-h), Similar to (e-f), but colored according to the predicted MCP values. (i-j), Similar to (e-f), but colored according to the predicted SA values by RaptorX. The PDB ID of the half-membrane-embedded protein was 5nup, which is an X-ray crystal structure of the Gram-negative bacterial 𝛼-helical outer membrane (OM) protein composed of 236 residues. The PDB ID of the soluble protein was 1a70, which is an X-ray crystal structure of ferredoxin I (Fd I) from Spinacia oleracea with 97 residues.

As a further test, we conducted the MCP prediction for a soluble protein dataset, composed of 102 Pfam proteins after removing the sequence redundancy with respect to our training set, which was previously used as a test set for contact map and protein structure prediction [6]. The results are shown in Fig. 3b. As can be seen, most of the amino acids were predicted to be not lipid-exposed (MCP < 0.5), which was expected since the dataset was supposed to contain soluble proteins only. However, there were still about 14 amino acids predicted to be likely having direct contact with the hydrophobic acyl chains of lipid molecules, which was unexpected.

These 14 amino acids were distributed in seven proteins (Table S5). Considering our average prediction performance was around 70% (PCC), we ruled out the cases in which there were less than three high-MCP predictions in ten successive amino acids of a protein, considering them outliers, and then there was only one protein left, whose sequence corresponds to a crystal structure of the Sar1-GDP complex [40]. Our prediction indicated that six of its residues should be in direct contact with the hydrophobic lipid acyl chains, and these residues are in proximity in the sequence (Table S5), located on the N-terminal helix of the protein Sar1. Two of the six residues were resolved in the X-ray structure, both of which were on the outer surface. We suspected that these residues may interact with the hydrophobic core of a membrane, although they belong to a soluble protein. Therefore, we ran multiscale MD simulations to check whether this soluble protein would have direct contact with the hydrophobic core of a lipid bilayer. Indeed, our self-assembly CG MD and atomistic MD simulation results demonstrated that this protein would be stably anchored onto a membrane’s surface (Fig. 5a-c), and both the coarse-grained and atomistic simulations confirmed a stable binding interface (Fig. 5d-g). In fact, previous experimental studies also showed that Sar1 is responsible for membrane trafficking, and its N-terminal helix probably serves as a wedge that inserts into the outer leaflet of the endoplasmic reticulum (ER) membrane and regulates the membrane curvature and fission [41]. Therefore, the ‘abnormal’ MCP prediction turned out to be a functionally relevant one: although some soluble proteins are not embedded in a lipid bilayer, they may bind and dip into the membrane deeply enough so that some amino acids can reach the hydrophobic core region.

**Figure 5.**
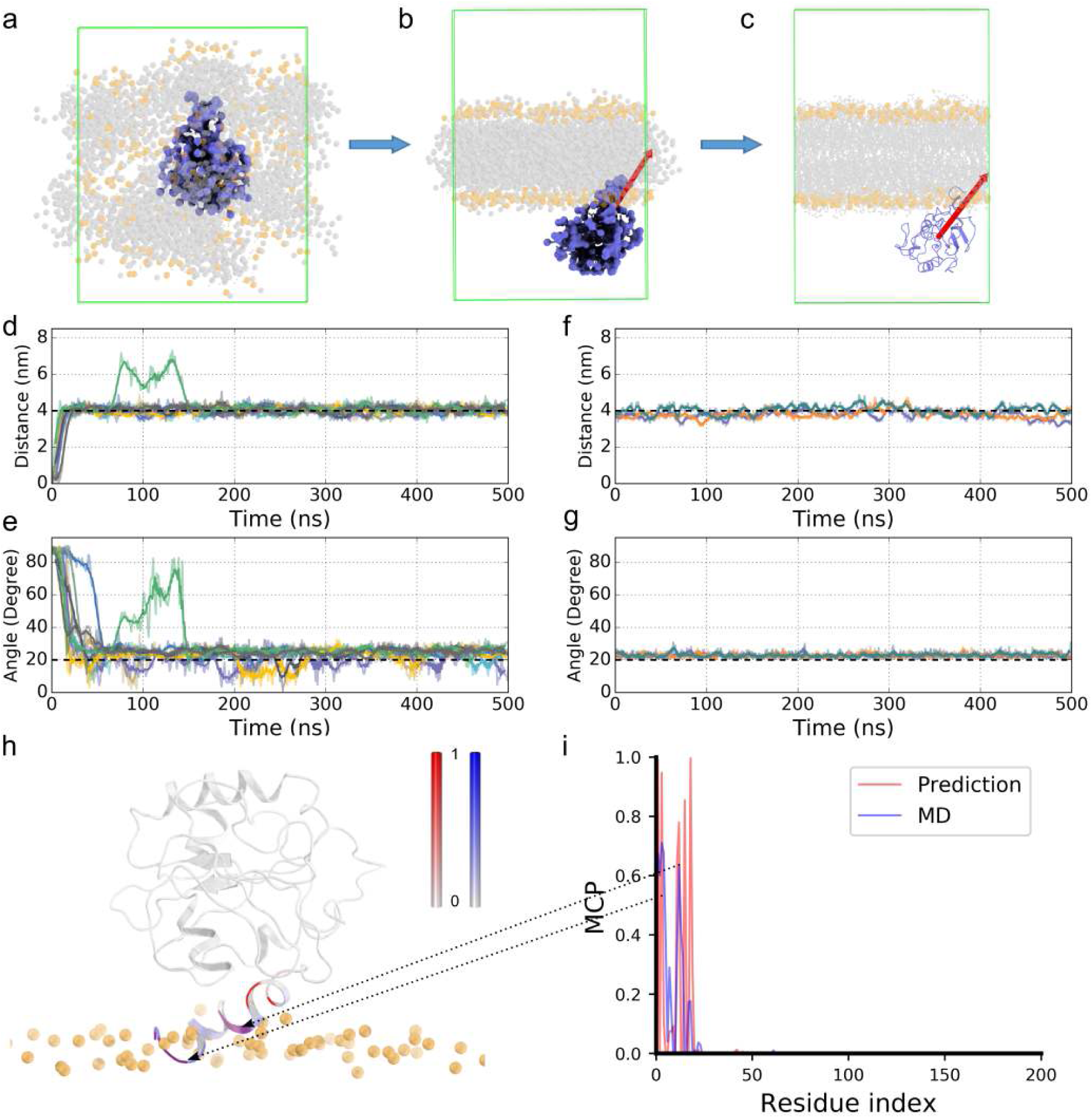
The Sar1 structure (1f6b) anchored on a membrane. (a) The initial and (b) final structures of the 1f6b system in the coarse-grained self-assembly MD simulations. (c) The atomistic structure of 1f6b transformed from the CG MD simulation outcome (b). The orange and grey spheres represent the lipid head groups and hydrophobic tails, respectively. The red arrow represents the first principal axis of the protein. Water molecules were filled in the whole simulation box, but are not shown here for clarity. (d) The distance between the COM of the protein and the COM of the bilayer in the nine CG simulation trajectories. (e) The orientation of 1f6b in the trajectories, represented by the angle between the first principal axis of the protein structure and the Z-axis of the simulation system. (f) Similar to (d), (g) similar to (e), but obtained from three atomistic MD simulations. (h) The orientation of 1f6b bound to the lipid bilayer obtained from MD simulations, with the membrane-interacting residues colored according to panel (i). (i) The overlaid MCP generated by our MCP predictor (red) and MD simulations (blue).

Following Stansfeld’s protein-lipid contacting definition [42], we determined the lipid-interacting residues as those within 6 Å of the lipid tails and calculated the lipid interacting probability of each residue in the simulations. The MCP values obtained from our MD simulations and DCRNN prediction were not completely identical (Fig. 5h&i), but the comparison demonstrated that they overlapped at residues M1, F3, G11, F12 and F18 (the purple regions in Fig. 5h), showing a converged membrane-interacting interface. Therefore, it appeared that the MCP predictor was good for the soluble proteins too, and perhaps could be used to identify the membrane-interacting residues of soluble or membrane-anchored proteins.

As a comparison, we used other software of relevant functions to conduct predictions for this protein (Fig. S4). From the result of BCL::Jufo9D [43], a server for the prediction of transmembrane span, the N-terminal helix of the protein Sar1 was predicted to be more likely a transition region (TR in Fig. S4), which is somewhat consistent with our results, but less quantitative and less specific when compared to the MCP prediction and the MD observations. The N-terminal helix was not predicted to be membrane-spanning by the transmembrane topology predictors OCTOPUS [32] or TMHMM [30] (Fig. S4).

### MCP can improve the prediction precision of protein contact maps and structures

SA has long been used as a fundamental input for protein structure prediction. As demonstrated, MCP is an essential character of amino acids and complementary to SA in describing the outer surface of membrane proteins, so it is natural to think that the MCP would be helpful for protein structure prediction, too. Notably, the prediction precision of protein contact map was hugely improved in the last couple of years, especially since the introduction of the ResNet into the field by the XU group [6, 44]. In the state-of-the-art ResNet method, as well as most if not all of the prediction methods, SA is an essential 1D input for the model. To validate the usefulness of the MCP, we incorporated the MCP into the ResNet the same way as SA, considering the MCP as a 1D input in parallel with SA (please refer to the method and Fig. 1c-d for details), and then we checked whether the contact map prediction can be improved.

We took the above dataset containing 327 test proteins as our prediction target, and we compared the prediction results of our MCP-incorporated ResNet contact map prediction with the original ResNet model. The top L/k (k=10, 5, 2, 1) results are shown in Table 2 and demonstrate that the inclusion of MCP in the ResNet predictor systematically improved the prediction precision. On average, the prediction precision can be improved by about 3%, if only the top L/k predictions are considered. The results showed that the MCP is particularly useful for the improvement of long-range contact prediction, with an improvement of up to 7% in the first L predictions of the long-range tests. In order to evaluate the prediction precision for the whole contact maps, we calculated the PCC between the predictions and the native contact maps calculated from known protein structures of the whole dataset. The PCC is about 0.29 with the original ResNet predictor and about 0.37 with our MCP-incorporated ResNet predictor (Fig. S5). Thus, the relative improvement of the prediction precision is about 28% with the incorporation of the MCP into the ResNet model, when the whole contact map is considered.

**Table 2:** The overall contact prediction precision of the 327 test proteins.

| Methods | Short |  |  |  | Medium |  |  |  | Long |  |  |  |
| --- | --- | --- | --- | --- | --- | --- | --- | --- | --- | --- | --- | --- |
|  | L/10 | L/5 | L/2 | L | L/10 | L/5 | L/2 | L | L/10 | L/5 | L/2 | L |
| CCMpred | 0.46 | 0.33 | 0.19 | 0.13 | 0.53 | 0.40 | 0.24 | 0.16 | 0.65 | 0.58 | 0.42 | 0.29 |
| ResNet | 0.75 | 0.62 | 0.39 | 0.23 | 0.80 | 0.70 | 0.49 | 0.31 | 0.89 | 0.85 | 0.74 | 0.59 |
| ResNet+MCP | 0.78 | 0.65 | 0.41 | 0.24 | 0.85 | 0.75 | 0.53 | 0.33 | **0.91** | **0.89** | **0.80** | **0.66** |

As further validation, we calculated the prediction precision for a dataset composed of 495 test proteins, which was a subset of PDB25 [45] created in May 2020 (25% non-redundant sequences with a resolution higher than 2.5 Å and R-factors less than 1.0). Any test proteins sharing > 25% sequence identity with any training proteins were excluded. Again, the ResNet predictor with MCP incorporated showed a consistently higher precision (Table S6). We further separated the dataset into two subsets, the soluble proteins and membrane proteins, and we calculated the prediction precision for each subset. As can be seen in Table S6, the prediction precision was improved by about 2% for the top predictions of all of the proteins in the dataset, which was not very significant. However, when we looked at the whole prediction rather than the top predictions of the contact maps, we can see that the MCP is highly useful to improve the prediction of membrane proteins, with the PCC values of 0.132 for the original ResNet predictor and 0.222 for our MCP-incorporated ResNet predictor when compared to the native contact maps of all the membrane protein in the dataset. Thus, the relative PCC improvement for membrane proteins was around 68%. The prediction for soluble proteins was improved too, with the values of 0.147 and 0.177 for the two models, respectively. Thus, the relative PCC improvement for soluble proteins was around 20%. Although the overall PCCs are still low for the entire contact maps, these results present a clear improvement in the contact map prediction when the MCP is incorporated into the ResNet model, and the improvement is more significant for membrane proteins than soluble proteins.

Using the aforementioned 5aym and 4e1t as representative cases, we analyzed the differences of the CM predictions before and after the incorporation of MCP in the ResNet model and compared to the results from other CM prediction tools [46–49] (Table S7). It is obvious that the MCP-incorporated ResNet predictor performed the best. In Fig. 6, we show the native contact maps, and the ResNet predictions with and without using the MCP information for the two membrane proteins, respectively. The prediction with the MCP is better correlated with the native CMs with a PCC of 0.53 and 0.69, while the prediction without the MCP showed a PCC of about 0.35 and 0.62 compared to the native CMs, respectively. Looking at the six panels of Fig. 6, one can immediately recognize that the incorporation of MCP removed many false positives compared to the model without MCP. As a result, the top L/5 long-range contact prediction precision was improved by 10% and 6% for the two cases, respectively (Table S7).

**Figure 6.**
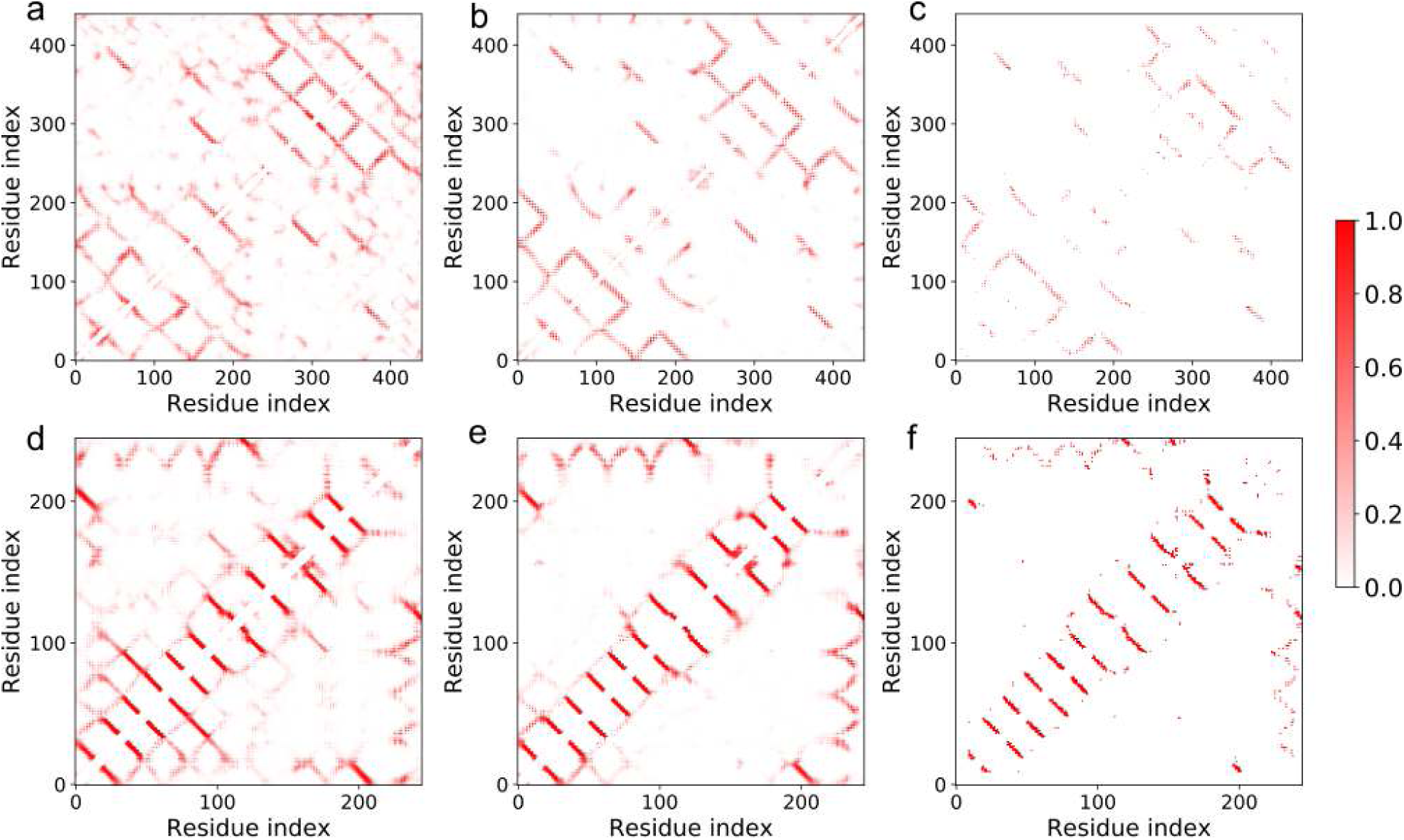
Incorporation of the predicted MCP improved the CM prediction for the two representative cases. The cutoff for the CM prediction was 8 Å between 𝛽 carbon atoms. (a), (b) The predicted CMs for 5aym without and with MCP incorporated, respectively. (c) The native CM calculated from the known structure 5aym. (d-f) Similar to (a-c), but for the protein 4e1t.

Based on the predicted contact maps, we further predicted the structures of the above two representative proteins using CONFOLD2 [50]. From the top-five models, it is clear that the prediction with the MCP-incorporated contact maps yield consistently better results than with the contact maps without MCP incorporated (Table S8). The best-predicted structures are shown in Fig. 7, where the predicted structures using the MCP-incorporated contact maps are closer to the crystal structures with RMSDs of 3.25 and 3.41 Å, while the predictions without MCP have RMSDs of 4.55 and 4.12 Å compared to the crystal structures, respectively.

**Figure 7.**
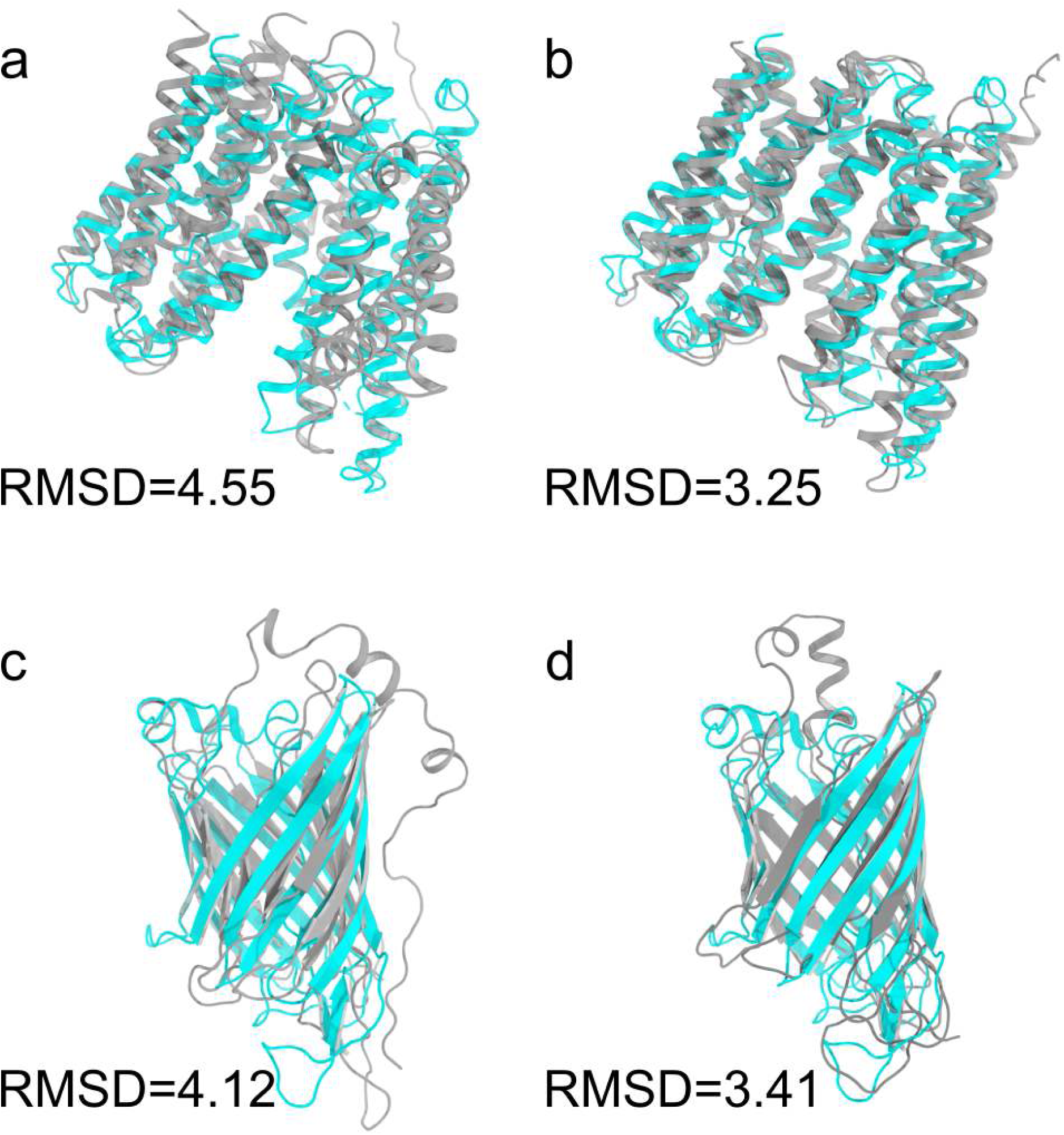
The MCP-incorporated CM predictions improved the protein structure prediction accuracy for the two representative cases. (a), (b) The predicted protein structures for 5aym using the predicted CMs without and with MCP incorporated, respectively. The crystal structure was shown in cyan and the predicted structures were shown in grey. (c-d) Similar to (a-b), but for the protein structure 4e1t.

## Discussion

Protein folding is driven by both enthalpy and entropy. The distribution of the amino acids on the outer surface of a protein is mainly determined by entropy. For a soluble protein, the majority of the outer-surface amino acids are hydrophilic to form a better match with the surrounding water molecules and maximize the entropy. This phenomenon has been widely recognized and utilized in accessing the structural and functional properties of soluble proteins for many years [51]. The widely-accepted quantity describing which amino acids are more likely to be exposed to water is solvent accessibility (SA), which is a crucial quantity for the analysis and prediction of soluble protein structures [4, 5, 8]. Amino acids with high SA values are likely located on the outer surface, while those with low SA values are probably embedded in the interior of the protein to reduce the hydrophobic mismatch, thus providing a strong conformational constraint on the protein structure for a given sequence. However, for membrane proteins, SA is not the whole story about their outer surfaces. A significant part of the outer-surface amino acids of membrane proteins are hydrophobic, and they have direct contact with the hydrophobic acyl chains, also driven by entropy. Therefore, high-SA amino acids cannot cover the full surface of a membrane protein, and low SA values may even apply a false constraint on the distribution of the outer-surface hydrophobic amino acids by wrongly embedding them into the interior of the membrane protein. Therefore, we believe a complementary quantity to describe the membrane contact probability of amino acids is highly valuable for membrane protein studies.

Lipid molecules are much larger and more complex than water molecules, introducing more uncertainties to the MCP calculation if the same roll-a-sphere protocol is adopted as in the SA calculations. For example, the interface of some oligomeric transmembrane proteins can be filled with lipid molecules, which is hard to determine from the structural viewpoint alone [52]. Therefore, here, we propose a new method to extract the MCP information for the membrane proteins with a known structure by utilizing the outcome of MD simulations (MemProtMD). The advantage of this method is that the MCP values were directly obtained from statistical analysis of the MD simulations, which contain the full details of the acyl chains moving around the membrane proteins, both spatially and temporally, and therefore provide much more accurate, dynamic, and complete information about the MCP.

Hydrophobic scale is widely used to characterize the hydrophobic property of amino acids, and thus can also be used to characterize/color the outer surface of a protein. We compared the MCP prediction to the Wimley-White hydrophobicity scale [53] to check whether MCP has any advantages over simple hydrophobicity scales. As shown in Fig. S6, it is obvious that our MCP prediction gives a much more clear representation of the membrane-interacting residues than the simple hydrophobicity scale for both membrane proteins and soluble proteins. The MCP prediction also agrees much better with the MD observations than the simple hydrophobicity scale, indicating that MCP is indeed a more appropriate quantity in characterizing the outer surface of a (membrane) protein.

As shown in Fig. 3, the MCP and SA are complementary to each other in defining the outer surface of a membrane protein. One can easily imagine that the inclusion of the MCP is beneficial to the structural analysis and prediction of proteins for two reasons. First, the incorporation of the MCP can tell which hydrophobic residues should face outward rather than be embedded in the interior (as learned from soluble proteins), and thus can reduce the false constraints forcing the hydrophobic residues to become embedded. Second, the opposite is probably true as well: the incorporation of the MCP can increase the true positives for the hydrophobic residues embedded inside and the hydrophilic residues exposed to the outside. Therefore, when we trained the MCP-incorporated ResNet model for the contact map prediction, the precision for both the soluble and membrane proteins was improved. As a natural result, we show that MCP is helpful for 3D protein structure prediction as well (Fig. 7 and Table S8). In addition to improving contact maps, which is helpful for 3D structure prediction, we believe MCP can also be directly used for 3D structure prediction and evaluation. For example, one can apply extra constraints to push the high-MCP residues onto the protein surface during the 3D structure modeling of membrane proteins, or take the MCP into account when scoring the modeled protein structures.

As only the single-chain information was used to construct the dataset, one might wonder whether the model works for oligomeric complexes too. Our analysis showed that: 1) The model performs better for single-chain proteins than for complexes (Table S9). 2) For a single chain within a complex, the protein-membrane and protein-protein interfaces can be distinguished from the predicted MCP (Fig. S7), where the protein-protein interface residues showed lower MCP predictions than those of the protein-membrane interface. However, one cannot tell where the protein-protein interface is from the MCP alone, as the low-MCP region may be embedded in the protein interior as well. 3) Similar to above, for a single-chain protein and a monomeric chain of a complex with similar fold but low sequence similarity, the predicted MCP can distinguish them (Fig. S8). Therefore, it appears that the MCP predictor works reasonably well for oligomeric transmembrane complexes too. Another potential application of MCP is to determine the correct orientation of membrane proteins of known structure when embedded into membranes, similar to OPM and MemEmbed [39, 54, 55], for example. The predicted MCP can also help to position a membrane-anchored protein in close proximity to a membrane with the interface residues facing the membrane surface, which would be useful for setting up structural models for further studies such as molecular dynamics simulations. We reserve to explore these potential applications in future studies.

There are several known limitations of the method presented in this work: 1) To achieve a better performance, the datasets contain redundant sequences, which may add bias to the model and makes it not really *De Novo*. Unfortunately, with the limited amount of membrane protein structures, it is difficult to use a low-sequence-similarity dataset to achieve the best prediction performance for now. 2) The prediction for the random coiled structures is not very reliable, which is natural considering these are the most flexible and least abundant structures in membrane proteins. 3) In the datasets extracted from MemProtMD, there was only one type of lipid molecules, 1,2-dihexadecanoyl-rac-glycero-3-phosphocholine (DPPC), making it impossible to identify lipid-type-specific interactions. However, the analysis showed that a membrane protein has very similar membrane contacts in spite of the lipid molecule types in the membrane (Fig. S9), so as far as the hydrophobic core-contacting information is concerned, the prediction should be satisfactory. 4) The highly probable lipid-interacting sites can be predicted, but one can not distinguish whether it is a specific lipid binding or a non-specific binding site from the prediction alone, as the training dataset does not contain such information, so the method cannot be used to identify specific lipid-binding sites. To overcome these problems, much larger datasets with more rich information, such as the distribution of diverse lipids and the lipid residence time around membrane proteins, would be required.

In summary, we propose that the MCP is an essential, characteristic, and predictive quantity of proteins that should be explicitly considered in the study of membrane proteins. The usage of MD outcomes as the training dataset for deep learning may generate a more accurate prediction for the MCP and other similar properties of proteins. We believe that the MCP would be able to find a wide range of applications in various aspects of the structure and function studies of membrane-interacting proteins.

## Materials and Methods

### MCP prediction

#### Extraction of the MCP information from the MemProtMD database

Our aim was to predict the MCP efficiently and accurately from any given protein sequence in the absence of structural information, which could then be used for structural and functional analysis of the protein. To do this, we needed a large dataset of the MCP of membrane proteins for the training using machine learning-based methods. It is difficult to obtain the MCP information directly from experiments, but molecular dynamics (MD) simulations have been proven to be a reliable method for studying the interactions between the membrane proteins with a known structure and lipid bilayers [56–58]. With MD simulations, we can get dynamic and quantitative information regarding the protein and membrane contact. In fact, Stansfeld et al. performed extensive MD simulations for all of the membrane proteins with a known structure and deposited the relevant data into the MemProtMD database [36, 37, 42], which paved the way for the current study.

The MemProtMD database contains the information about the stable orientation of the membrane proteins with a known structure in an explicit lipid bilayer environment. The information was obtained by running sophisticated multi-scale MD simulations [56–58]. The database also contains statistical information of the contact between the membrane proteins and lipid bilayer. As it is more difficult to discriminate the headgroup contact from the solvent contact, we chose to only consider the hydrophobic acyl tail contact of each protein residue at this stage. We considered that a certain protein residue was in direct contact with the hydrophobic acyl tail if the distance between them was less than 6 Å. Such a cutoff value was shown to be appropriate for discriminating the transmembrane region from water-exposed regions of membrane proteins [56, 59] (Fig. S10). The contact probability was defined as the average occupancy of the selected groups in direct contact calculated over the final 800 ns of the coarse-grained (CG) MD trajectory, after each membrane protein simulation system reached equilibrium. There are more than 3,500 MD simulation results in the web database and the number continues to count (http://memprotmd.bioch.ox.ac.uk/) [37]. We downloaded the PDB files of the atomistic structure with the acyl tail contact probability from the MemProtMD database. In the PDB files, the temperature factor value (also called the B-factor) of each atom was replaced by the membrane contact probability obtained from MD analysis. Therefore, we were able to extract the membrane contact probability value of each C_𝛼_ as our MCP observation for each residue. This procedure is shown in Fig. 1a.

We extracted 3604 simulation results from the web database in April 2019. There were about 90% 𝛼-helical membrane proteins and 10% 𝛽-barrel membrane proteins in the dataset. We separated each file by the number of chains. We excluded the sequences that were longer than 700 residues or shorter than 26 residues.

During the dataset generation, the multi-chain proteins were split into single-chain sequences, and only one chain of the same sequence was adopted into the dataset. We obtained 12691 result files, removed duplicate sequences, and extracted 5900 of them at random as the membrane protein dataset in the end.

#### Generation of the dataset for the MCP model training

The values of MCP lie between 0 and 1, with 0 meaning no contact at all and 1 meaning persistent contact with the hydrophobic core of membranes throughout the simulation time. A fractional number tells in what percentage of the simulation time a direct contact between an amino acid and hydrophobic acyl chains of lipid molecules was observed. The above information extracted from MemProtMD provided the original dataset for the sequences of membrane proteins with a known structure.

To do the MCP prediction, we also included soluble proteins in the dataset. The training dataset was composed of 5000 membrane protein sequences from the MemProtMD database and 5000 soluble protein sequences. The soluble protein sequences were chosen randomly from the soluble proteins of the PDB25 dataset with less than a 25% sequence identity [45]. Then, we set the MCP of each soluble protein residue to be zero. In addition, we divided the remaining 900 membrane protein sequences into two subsets: 500 membrane protein sequences were used as the test set, and the other 400 membrane protein sequences were used as the validation set. The above dataset was termed ‘MCP-Large’ in this work.

We also constructed a dataset with less-redundancy sequences, in which we only used the sequences with less than a 40% identity between any two sequences in the original dataset of membrane proteins, resulting in 898 sequences. This smaller dataset was termed ‘MCP-Small’. To ensure that the training set is relatively sufficient [24, 60], the 898 sequences were divided into three subsets: 718 randomly chosen membrane protein sequences formed the training set, 90 membrane protein sequences were used as the test set, and the other 90 membrane protein sequences were used as the validation set. Such a ∼8:1:1 dataset construction ensures a relatively larger training set with reasonable validation and test sets for better convergence, which was adopted and recommended by previous work [61, 62].

#### The DCRNN model for the MCP prediction

We considered the MCP prediction as a regression problem and used a combination of deep convolutional and recurrent neural network (DCRNN) to do the prediction, which is one of the state-of-the-art models used for protein secondary structure prediction [35]. Due to the long-range dependencies in the protein sequence-based model, we referred to the bidirectional gate recurrent units (BGRUs) for the global context extraction, which contains a forward gate recurrent unit (GRU) [63] and a backward GRU. In the model, we also combined multiscale CNN layers for the local context extraction.

As illustrated in Fig. 1b, we can see an overview of the model for the MCP prediction. We defined the loss function as the residual sum of the squares between the values from the prediction and the MD observation plus L_2_ norms of the model parameters.

In our model, we utilized the protein features generated by RaptorX-Property [4], including the predicted three-state secondary structure (SS3), three-state solvent accessibility (SA), and PSSM, which were concatenated to be a 1×26 array for each residue. For a protein with a sequence length of L, the resulting matrix had a dimension of L×26, which was padded with zeros to be a 700×26 matrix for the sequences shorter than 700. Then the matrix was operated by a sliding window with the kernel size of k×26 (k=3, 7, 9) and a channel size of 64, as shown in Fig. 1b. For each protein sequence, we ran HHblits 3.0.3 [64] (with E-value 0.001 and 3 iterations) to search the uniclust30 database dated October 2017 to find its sequence homologous and then built its multiple sequence alignment (MSA). We only consider single chains for the prediction for now.

Our code was implemented with Tensorflow (https://www.tensorflow.org) of Python (https://www.python.org/). The weights in our neural networks were initialized with the default parameters in Tensorflow. We trained all of the layers in the deep network using the Adam optimizer [65]. We set the batch size to be 1. The training was conducted on a workstation with a six-core (12-thread) Intel Xeon E5-1650 CPU and a GTX 1080 Nvidia GPU. It took around 24 hours to train one model with 200 epochs.

The model reached convergence after 20 epochs of training according to the MSE curves, and we stopped the training at 100 epochs when no sign of over-training was observed (Fig. S11). The 10-fold validation results are shown in Table S10 and Table S11.

### Protein contact map prediction

#### The dataset for the protein contact map prediction

Our training data were a subset of PDB25 [45] created in April 2018, which only included proteins with less than a 25% sequence identity. We excluded a protein from the training set if it met one of the following conditions: (1) sequence length shorter than 26 or larger than 700, (2) resolution worse than 2.5 Å, (3) has domains made up of multiple protein chains, or (4) has unusual amino acids other than the 20 standard ones. In the end, there were 10054 sequences in our training set, which contained around 150 membrane proteins. We did not manually balance the soluble and membrane proteins during training, as previous studies showed that ResNet works for membrane proteins even if the training set only contained a small fraction of membrane proteins [44], meaning the learning is quite transferable. The basic statistics of the dataset are shown in Table S12 according to the database SCOPe [66, 67].

#### The ResNet model for the protein contact map prediction

The deep residual net (ResNet) has been widely used for image recognition and won first place on the ILSVRC 2015 classification task [68]. Xu et al. developed and utilized the ResNet for the contact map prediction of proteins and won first place on CASP 12 and CASP13 (RaptorX). To test if the MCP is useful for the structural prediction of membrane proteins, we integrated the MCP into the ResNet model created by Xu et al. [6] and checked whether the prediction performance could be improved. The revised model contained two residual neural networks. Fig. 1e shows each residual block of ResNet.

For the design of the convolution kernel, we used 17 in 1-D convolution and 5× 5 in 2-D convolution like the original implementation in the RaptorX-Contact [6]. We constructed the model with 60 2D convolutional layers and two 1D convolutional layers when we combined the model with the MCP.

We used similar input features as the original ResNet model [6], plus the additional MCP predicted from the protein sequence in parallel to SA as a 1D input (Fig. 1d). In addition to the MCP predicted by our model, the input features included the PSSM, SS3, and SA generated by RaptorX-Property [4], as well as the evolutionary coupling (EC) information generated by CCMpred [46], pairwise potential, and mutual information generated by alnstats [69]. The pairwise potential is computed by averaging contact potential terms [70, 71] across the two alignment columns, derived from the MSA. The mutual information is calculated between positions of two different protein families in a joint alignment of sequences from the same set of organisms [72]. For each protein sequence, we ran HHblits 3.0.3 (with an E-value of 0.001 and 3 iterations) to search the uniclust30 database dated October 2017 to find its sequence homologous and then built its MSA. Then we generated sequence profiles from the MSA and predicted all of the needed features above.

The prediction of the protein CM was transformed into a binary classification problem. For each amino acid pair, we restricted the prediction result within [0, 1] through the sigmoid function, which represents the possibility of the two residues (𝛽 carbon) within a distance of 8 Å, a cutoff value widely accepted in the field. Therefore, the output of the contact map prediction was a matrix showing the probability of two residues within 8 Å.

We then evaluated the prediction precision of the top L/k (k = 10,5,2,1) predicted contacts, where L is the protein sequence length, by comparing the predictions with the native contacts calculated from known protein structures. The prediction precision was defined as the percentage of native contacts among the top L/k predicted contacts. We also divided the contacts into three groups according to the sequence distance of two residues. A contact is short-, medium-, and long-range when its sequence distance falls into [6, 11], [12, 23], and ≥ 24, respectively.

For the MCP-incorporated ResNet CM predictor, we compared the CM prediction results using the MCP predictors trained by the MCP-Large dataset and the MCP-Small dataset. In the end, we used the MCP-Large predictor for the CM prediction in this work.

Our code was implemented with Tensorflow in Python. Weights in our neural networks were initialized with the default parameters in Tensorflow. We used the Adam optimizer to do the training with a batch size of 1. The training was conducted with a Tesla V100 Nvidia GPU with 32 GB of GPU memory, on which it takes around 40 hours to train a model with 20 epochs.

The model reached convergence after 20 epochs of training according to the precision curves (Fig. S12). The 10-fold validation results are shown in Table S13. The normalized confusion matrix, the prediction accuracy, and the area under the curve (AUC) are shown in Table S14 and S15, respectively. A cutoff of 0.5 was used for the confusion matrix and prediction accuracy calculation. Although these are not commonly used to evaluate the model performance in the CM predictions as the CM matrices are highly sparse, they show that the MCP-incorporated CM predictor consistently outperforms the ResNet model without MCP.

### Contact-driven protein structure prediction

With the predicted contact maps and the predicted three-state secondary structures [73], we can build protein structure models of a query sequence using CONFOLD2 [50]. With the scripts in the CONFOLD2 package, we converted the predicted secondary structures to distance, dihedral, and hydrogen bond restraints. We used the top-xL contacts as contact distance restraints, where x = 0.1, 0.2, 0.3, …, 4.0 and L is the length of the protein. For each predicted contact, the distance of the two corresponding residues was set in the range from 3.5 Å to 8 Å. Then we fed the processed data to the Crystallography & NMR System (CNS) [74] for model construction. For each x value, 20 structure models were constructed, and then the top-five models in each subset were selected according to the contact energy score [50], resulting in 200 models in total. Then, we ranked these 200 models using the satisfaction score according to their top L/5 long-range contacts [50]. We selected the top-50 models and clustered them into five subsets with the pairwise TM-score [75]. Finally, we selected the model closest to the centroid of each cluster and obtained five top models. Then we calculated the RMSD value of each model with regard to the crystal structure (Table S8). In the above processes, the contact maps (with or without MCP incorporated) were the only difference for the model construction.

### Molecular dynamics simulations

We used MD simulations to validate whether the predicted soluble protein with multiple high-MCP amino acids was indeed interacting with and anchored into the membrane. The atomistic protein structure was downloaded from Protein Data Bank (PDB ID: 1f6b). There were missing residues (residue 1-12 in 1f6b) in the N-terminus. To avoid the uncertainty induced by the incomplete protein structure, we filled these missing residues with MODELLER [76]. According to the secondary structure prediction results generated by RaptorX [4], we constrained the residue 3-9 of 1f6b to form a helix. 15 possible structures were generated by MODELLER and the best-scored structure was picked as the initial structure for the following MD simulations.

First, coarse-grained (CG) MD simulations were performed with the MARTINI 3.0 force field [77]. The CG structure and topology files were generated with the script *martinize.py* [78, 79]. Following Stansfeld’s protocol when generating the MemProtMD database [36], we built the lipid-around-protein system with the self-assembly protocol, in which the protein was put into the simulation box with a random pose and 1,2-dihexadecanoyl-rac-glycero-3-phosphocholine (DPPC) molecules were placed randomly around the protein (Fig. 5a). After a tens-of-nanoseconds simulation, the lipid molecules formed a bilayer spontaneously, and the protein found its most stable orientation (Fig. 5b). The elastic network (ELN) was used to maintain the global conformation of the proteins during the simulations. To reproduce the flexibility of the loop, we removed the ELN between all of the loop regions and their neighboring residues. Before performing production MD simulations, we equilibrated the system to eliminate inappropriate contacts and reach the target conditions. After the 5000-step energy minimization procedure, 0.5-ns NVT (canonical ensemble) equilibration was performed with a time step of 20 fs. Then, we ran nine 500-ns independent simulations with a time step of 20 fs under the NPT (isothermal-isobaric) ensemble for each system. The V-rescale algorithm and the Berendsen algorithm were used to maintain the system temperature (310 K) and pressure (1.0 Bar) [80, 81], respectively. The electrostatic interactions were calculated with the reaction-field method. The Coulomb interaction and van der Waals interaction were both cut off at 1.1 nm.

Then the script *backward.py* [82] was used to transform the equilibrated coarse-grained system (Fig. 5b) to all-atom system, which was utilized as the initial system (Fig. 5c) for the following all-atom simulations. After the 5000-step energy minimization, the system was equilibrated for 0.5 ns in the NVT ensemble and 1.0 ns in the NPT ensemble. Position restraints with a force constant of 1000 kJ/mol/nm^2^ were applied on all heavy atoms of the protein to maintain the conformation during the equilibration process. The Berendsen algorithm [81] was used to keep the system temperature and pressure at 310 K and 1.0 Bar, respectively. The van der Waals interaction were cut off at a distance of 1.0 nm. The long-range electrostatic interactions were calculated with the Particle-Mesh Ewald (PME) method [83]. After the system was equilibrated to the desired condition, we removed the position restraints and performed three 500-ns all-atom MD simulations to evaluate the stability of the protein anchoring on the membrane surface. The temperature and pressure coupling algorithms were set to V-rescale and Parrinello-Rahman [80, 84] to maintain the system temperature at 310 K and pressure at 1.0 Bar, respectively. Both coupling constants were set to 1.0 ps. The all-atom MD simulation was performed with the *Amber99sb-ildn* force field [85] in combination with the *Slipids* force field [86, 87].

## Data Availability

The source code of the MCP predictor and MCP-incorporated CM predictor can be found on GitHub (https://github.com/computbiophys), and a public computation server can be found online (http://www.songlab.cn).

## Author Contributions

C.S. conceived the idea and supervised the project. L.W. designed the MCP predictor and performed the calculation and analysis. J.Z. reproduced the ResNet model. D.W. performed molecular dynamics simulations and related analyses. C.S., L.W., and D.W. wrote the manuscript.

## Supporting information

Dataset_Used_in_this_Work

## Acknowledgements

We thank Xiao Luo and Prof. Minghua Deng at the School of Mathematical Sciences at Peking University, who discussed with us and provided instructive suggestions. This work was supported by grants from the Ministry of Science and Technology of China (2021YFE0108100 to C.S.) and the National Natural Science Foundation of China (21873006 and 32071251 to C.S.). Part of the MD simulations was performed on the Computing Platform of the Center for Life Sciences at Peking University.

## Legends for Supporting Information

**File S1** Datasets used in this work. This file contains the datasets for the MCP model and contact map predictor in this work.

## Supplementary Information for

### Supplementary Tables

**Table S1:** The related prediction methods for membrane protein features.

| Method | Year | Model | Prediction | URL |
| --- | --- | --- | --- | --- |
| α-helical transmembrane protein | | | | |
| TMHMM2.0 [1] | 2001 | HMM | Topology | http://www.cbs.dtu.dk/services/TMHMM/ |
| MEMSAT3 [2] | 2007 | NN |  | http://bioinf.cs.ucl.ac.uk/?id=756 |
| OCTOPUS [3] | 2008 | HMM, ANN |  | http://topcons.net/ |
| SCAMPI2 [4] | 2016 | HMM |  | http://scampi.bioinfo.se/ |
| MemBrain3.0 [5] | 2020 | ResNet |  | http://www.csbio.sjtu.edu.cn/bioinf/MemBrain/ |
| PolyPhobius [6] | 2005 | HMM | Topology<br>Signal peptide | http://phobius.sbc.su.se/poly.html |
| Philius [7] | 2008 | DBN |  | http://www.yeastrc.org/philius/pages/philius/runPhilius.jsp |
| SPOCTOPUS [8] | 2008 | HMM, ANN |  | http://topcons.net/ |
| TOPCONS2 [9] | 2009 | HMM |  | http://topcons.net/ |
| MEMSAT-SVM [10] | 2009 | SVM |  | http://bioinf.cs.ucl.ac.uk/psipred/ |
| CCTOP [11] | 2015 | HMM |  | http://cctop.enzim.ttk.mta.hu/ |
| MPRAP [12] | 2010 | SVM | Surface accessibility | https://mprap.cbr.su.se/ |
| β-barrel transmembrane protein | | | | |
| PROFtmb [13] | 2006 | HMM | Topology | https://www.predictprotein.org/ |
| BetAware [14] | 2013 | ELM, GRHCRF |  | http://betaware.biocomp.unibo.it/BetAware |
| BOCTOPUS2 [15] | 2016 | SVM, HMM |  | http://boctopus.bioinfo.se/ |
| PRED-TMBB2 [16] | 2016 | HMM |  | http://www.compgen.org/tools/PRED-TMBB2/ |
| α-helical transmembrane protein or β-barrel transmembrane protein | | | | |
| BCL::Jufo9D [17] | 2013 | ANN | Transmembrane spans | http://meilerlab.org/index.php/servers/show?s_id=5 |
| mp_lipid_acc [18] | 2017 | Concave hull algorithm | Lipid accessibility | https://rosie.graylab.jhu.edu/mp_lipid_acc/submit |

HMM: Hidden Markov model; NN: Neural network; DBN: Dynamic Bayesian network; ANN: Artificial neural network; SVM: Support vector machine; ELM: Extreme learning machine; GRHCRF: Grammatical-restrained hidden conditional random field.

**Table S2:** The performance of the MCP predictor using the MCP-Small dataset.

| Evaluation | Training | Validation | Test |
| --- | --- | --- | --- |
| Overall |  |  |  |
| MSE | 0.057 | 0.074 | 0.080 |
| PCC | 0.705 | 0.656 | 0.635 |
| α-helix (H) | | | |
| MSE | 0.063 | 0.080 | 0.090 |
| PCC | 0.819 | 0.766 | 0.728 |
| β-sheet (E) | | | |
| MSE | 0.032 | 0.045 | 0.041 |
| PCC | 0.706 | 0.598 | 0.575 |
| Coil (C) |  |  |  |
| MSE | 0.014 | 0.021 | 0.018 |
| PCC | 0.618 | 0.446 | 0.429 |

**Table S3:** The performance of our MCP predictor for different transmembrane protein classes.

| Class | Single-pass α-helical | Multi-pass α-helical | β-barrel |
| --- | --- | --- | --- |
| MSE (train) | 0.063 | 0.051 | 0.042 |
| PCC (train) | 0.821 | 0.830 | 0.745 |
| MSE (validation) | 0.061 | 0.054 | 0.043 |
| PCC (validation) | 0.816 | 0.818 | 0.700 |
| MSE (test) | 0.060 | 0.055 | 0.043 |
| PCC (test) | 0.800 | 0.818 | 0.728 |

**Table S4:** The amount of proteins of different classes in the datasets.

| Class | Single-pass α-helical | Multi-pass α-helical | β-barrel |
| --- | --- | --- | --- |
| MCP-Large |  |  |  |
| train | 1062 | 2832 | 1106 |
| validation | 81 | 231 | 88 |
| test | 103 | 284 | 113 |
| MCP-Small |  |  |  |
| train | 134 | 367 | 217 |
| validation | 16 | 44 | 30 |
| test | 19 | 46 | 25 |

**Table S5:**
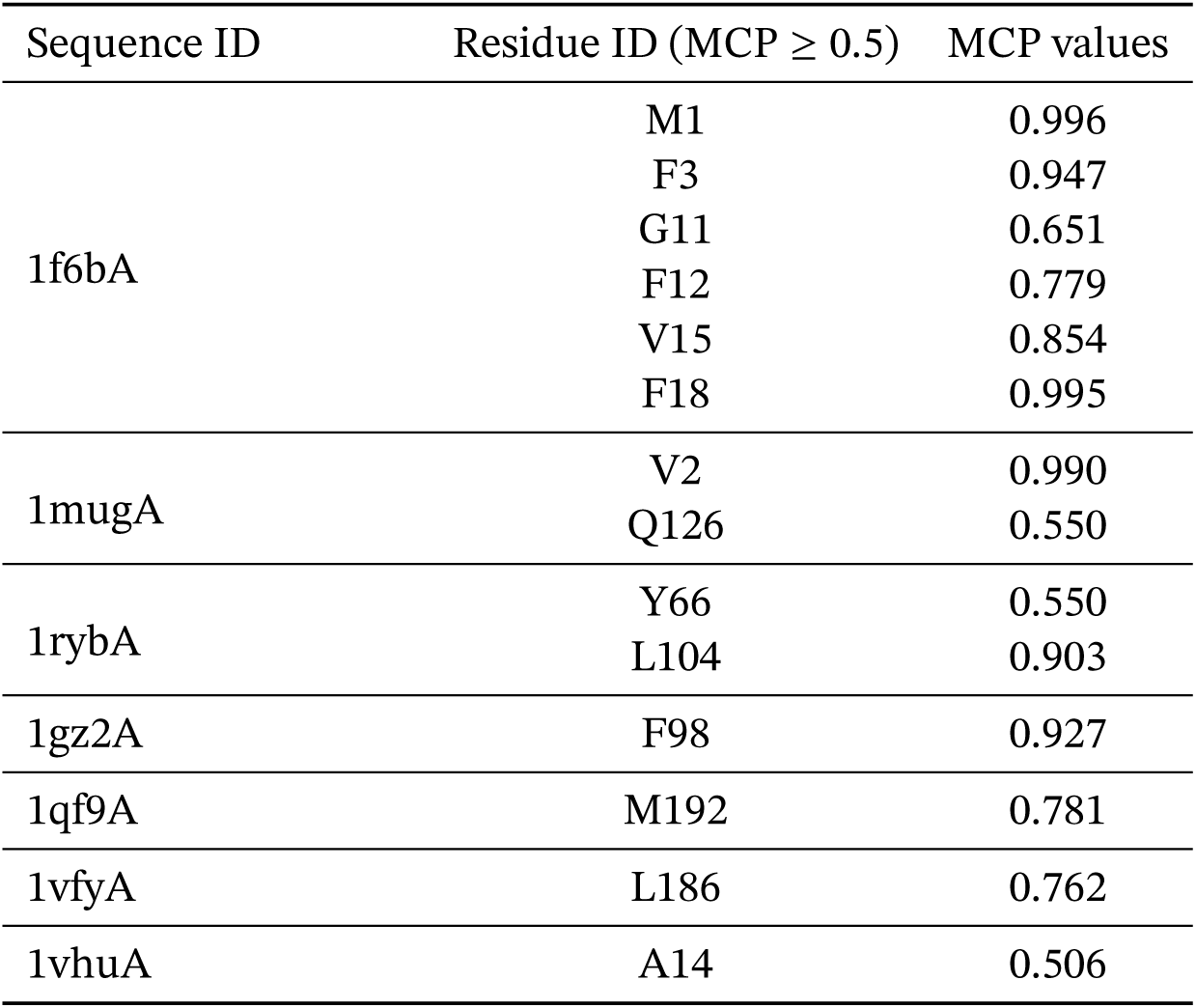
The amino acids predicted with high MCP values in the 102 Pfam soluble protein dataset.

| Sequence ID | Residue ID (MCP ≥ 0.5) | MCP values |
| --- | --- | --- |
| 1f6bA | M1 | 0.996 |
|  | F3 | 0.947 |
|  | G11 | 0.651 |
|  | F12 | 0.779 |
|  | V15 | 0.854 |
|  | F18 | 0.995 |
| 1mugA | V2 | 0.990 |
|  | Q126 | 0.550 |
| 1rybA | Y66 | 0.550 |
|  | L104 | 0.903 |
| 1gz2A | F98 | 0.927 |
| 1qf9A | M192 | 0.781 |
| 1vfyA | L186 | 0.762 |
| 1vhuA | A14 | 0.506 |

**Table S6:** The contact map prediction precision of the additional 495-protein dataset: overall, soluble, and membrane proteins.

| 495 test proteins in total |  |  |  |  |  |  |  |  |  |  |  |  |
| --- | --- | --- | --- | --- | --- | --- | --- | --- | --- | --- | --- | --- |
| Methods | Short |  |  |  | Medium |  |  |  | Long |  |  |  |
|  | L/10 | L/5 | L/2 | L | L/10 | L/5 | L/2 | L | L/10 | L/5 | L/2 | L |
| ResNet | 0.75 | 0.62 | 0.38 | 0.22 | 0.78 | 0.67 | 0.44 | 0.27 | 0.86 | 0.82 | 0.71 | 0.55 |
| ResNet + MCP | 0.76 | 0.64 | 0.40 | 0.23 | 0.79 | 0.69 | 0.46 | 0.28 | 0.86 | 0.83 | 0.73 | 0.57 |
| 480 soluble proteins |  |  |  |  |  |  |  |  |  |  |  |  |
| Methods | Short |  |  |  | Medium |  |  |  | Long |  |  |  |
|  | L/10 | L/5 | L/2 | L | L/10 | L/5 | L/2 | L | L/10 | L/5 | L/2 | L |
| ResNet | 0.75 | 0.62 | 0.38 | 0.22 | 0.78 | 0.68 | 0.45 | 0.27 | 0.86 | 0.82 | 0.71 | 0.56 |
| ResNet + MCP | 0.76 | 0.64 | 0.40 | 0.23 | 0.80 | 0.69 | 0.46 | 0.28 | 0.86 | 0.83 | 0.73 | 0.57 |
| 15 membrane proteins |  |  |  |  |  |  |  |  |  |  |  |  |
| Methods | Short |  |  |  | Medium |  |  |  | Long |  |  |  |
|  | L/10 | L/5 | L/2 | L | L/10 | L/5 | L/2 | L | L/10 | L/5 | L/2 | L |
| ResNet | 0.59 | 0.47 | 0.27 | 0.15 | 0.64 | 0.53 | 0.30 | 0.17 | 0.82 | 0.79 | 0.67 | 0.50 |
| ResNet + MCP | 0.60 | 0.50 | 0.29 | 0.16 | 0.66 | 0.53 | 0.31 | 0.18 | 0.83 | 0.80 | 0.70 | 0.52 |

**Table S7:** The contact map prediction precision for the two representative cases: 5aym and 4e1t.

| Contact prediction precision for 5aym |  |  |  |  |  |  |  |  |  |  |  |  |
| --- | --- | --- | --- | --- | --- | --- | --- | --- | --- | --- | --- | --- |
| Methods | Short |  |  |  | Medium |  |  |  | Long |  |  |  |
|  | L/10 | L/5 | L/2 | L | L/10 | L/5 | L/2 | L | L/10 | L/5 | L/2 | L |
| CCMpred | 0.16 | 0.09 | 0.05 | 0.04 | 0.07 | 0.05 | 0.04 | 0.03 | 0.34 | 0.18 | 0.10 | 0.08 |
| PSICOV | 0.30 | 0.18 | 0.09 | 0.06 | 0.20 | 0.11 | 0.06 | 0.05 | 0.73 | 0.64 | 0.41 | 0.29 |
| MetaPSICOV | 0.39 | 0.26 | 0.14 | 0.08 | 0.36 | 0.25 | 0.15 | 0.08 | 0.66 | 0.64 | 0.50 | 0.35 |
| MetaPSICOV2 | 0.41 | 0.27 | 0.15 | 0.08 | 0.43 | 0.28 | 0.16 | 0.10 | 0.77 | 0.68 | 0.56 | 0.38 |
| ResNet | 0.48 | 0.32 | 0.16 | 0.09 | 0.55 | 0.35 | 0.20 | 0.11 | 0.86 | 0.76 | 0.58 | 0.48 |
| ResNet + MCP | 0.59 | 0.35 | 0.16 | 0.09 | 0.70 | 0.42 | 0.21 | 0.11 | **0.95** | **0.86** | **0.76** | **0.54** |
| Contact prediction precision for 4e1t |  |  |  |  |  |  |  |  |  |  |  |  |
| Methods | Short |  |  |  | Medium |  |  |  | Long |  |  |  |
|  | L/10 | L/5 | L/2 | L | L/10 | L/5 | L/2 | L | L/10 | L/5 | L/2 | L |
| PSICOV | 0.33 | 0.31 | 0.20 | 0.17 | 0.46 | 0.29 | 0.20 | 0.17 | 0.38 | 0.27 | 0.20 | 0.13 |
| CCMpred | 0.54 | 0.51 | 0.34 | 0.22 | 0.71 | 0.53 | 0.34 | 0.21 | 0.54 | 0.53 | 0.30 | 0.18 |
| MetaPSICOV | 0.79 | 0.82 | 0.59 | 0.38 | 0.83 | 0.82 | 0.68 | 0.49 | 0.58 | 0.43 | 0.36 | 0.23 |
| MetaPSICOV2 | 0.88 | 0.82 | 0.66 | 0.39 | 1.00 | 0.94 | 0.77 | 0.56 | 0.71 | 0.61 | 0.43 | 0.28 |
| ResNet | 1.00 | 0.96 | 0.79 | 0.44 | 1.00 | 0.98 | 0.91 | 0.66 | 1.00 | 0.92 | 0.80 | 0.56 |
| ResNet + MCP | 1.00 | 0.98 | 0.81 | 0.44 | 1.00 | 1.00 | 0.96 | 0.66 | **1.00** | **0.98** | **0.83** | **0.63** |

**Table S8:** The RMSD of the top five structure prediction models with respect to the crystal structures for the two representative cases: 5aym and 4e1t.

| 5aym |  |  |
| --- | --- | --- |
| Model | RMSD (Å) |  |
|  | Without MCP | With MCP |
| 1 | **4.55** | **3.25** |
| 2 | 4.73 | 3.60 |
| 3 | 5.33 | 3.61 |
| 4 | 5.48 | 3.73 |
| 5 | 6.00 | 3.86 |
| 4e1t |  |  |
| Model | RMSD (Å) |  |
|  | Without MCP | With MCP |
| 1 | **4.12** | **3.41** |
| 2 | 4.32 | 4.05 |
| 3 | 4.43 | 4.09 |
| 4 | 4.56 | 4.10 |
| 5 | 6.33 | 5.25 |

**Table S9:**
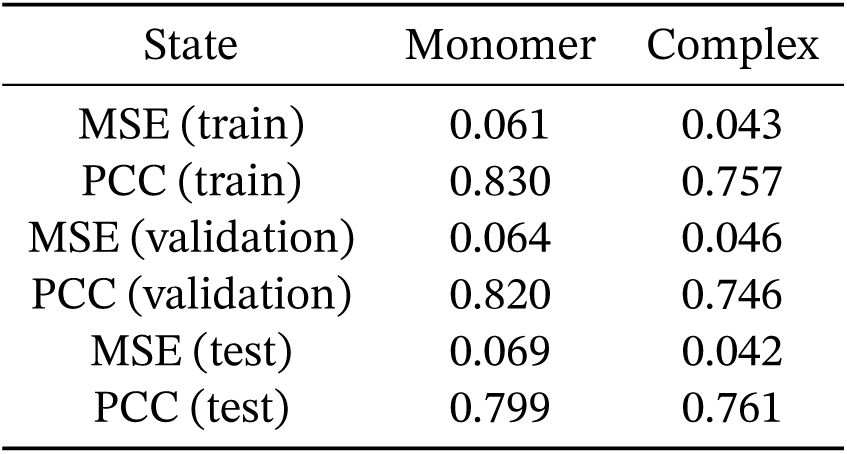
The performance of our MCP predictor for different oligomeric states.

**Table S10:**
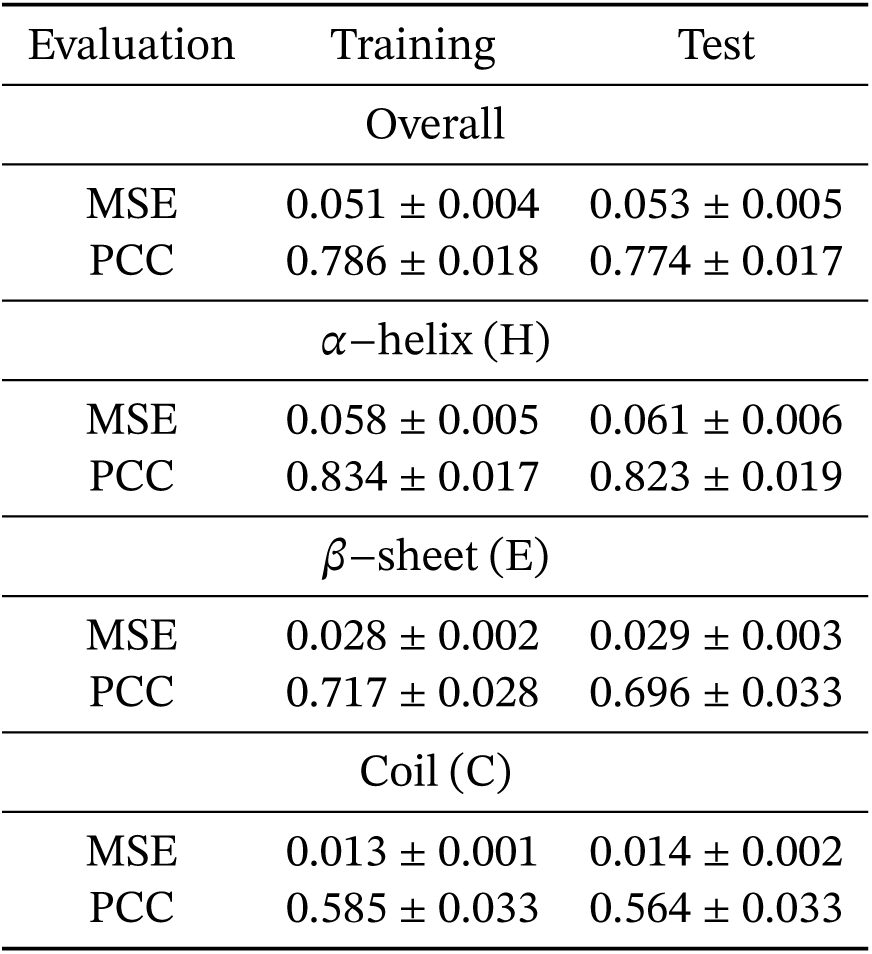
The performance of our MCP predictor in the 10-fold cross-validation using the MCP-Large dataset.

**Table S11:**
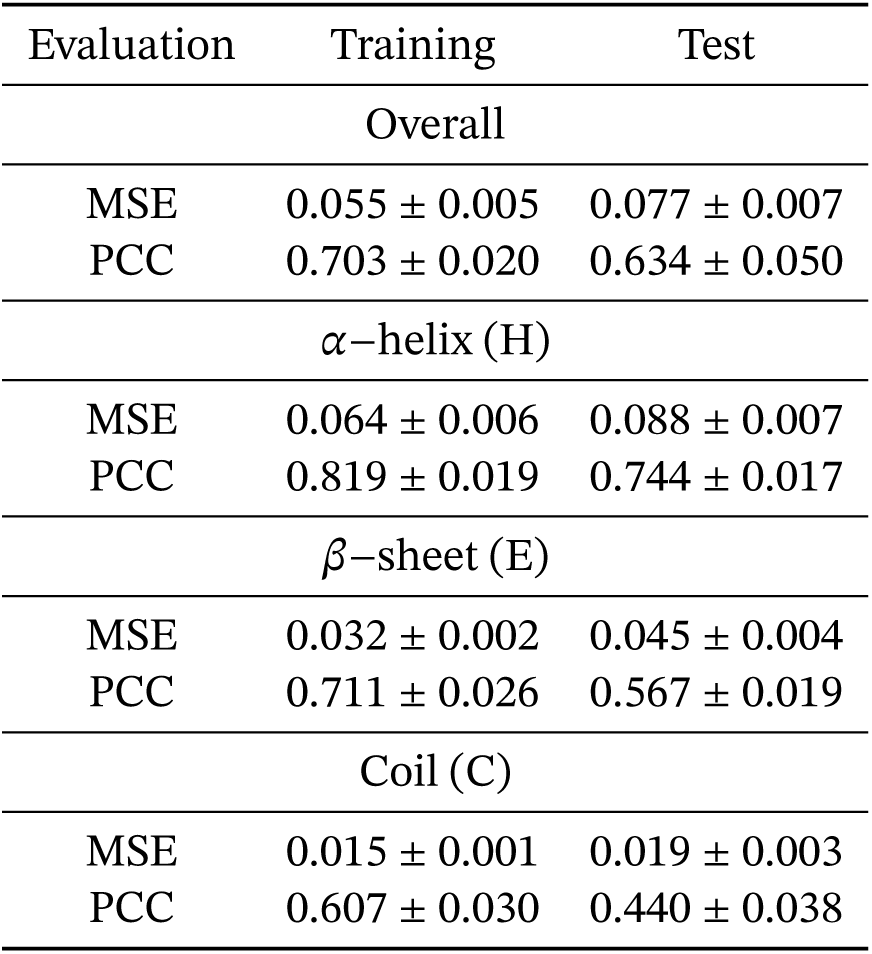
The performance of our MCP predictor in the 10-fold cross-validation using the MCP-Small dataset.

**Table S12:** The basic statistics of the datasets according to the database SCOPe [19, 20].

| Dataset | All alpha | All beta | Alpha/beta | Alpha+beta | Others | Sequence identity |
| --- | --- | --- | --- | --- | --- | --- |
| 10054 proteins | 641 | 604 | 676 | 934 | 7199 | 25% |
| 327 test proteins | 21 | 26 | 36 | 25 | 219 | 25% |
| 495-protein dataset | 22 | 32 | 21 | 28 | 392 | 25% |

**Table S13:** The performance (precision of medium- and long-range contact) of our contact map predictor in the 10-fold cross-validation.

| Methods | Medium |  |  |  | Long |  |  |  |
| --- | --- | --- | --- | --- | --- | --- | --- | --- |
|  | L/10 | L/5 | L/2 | L | L/10 | L/5 | L/2 | L |
| ResNet | 0.795<br>$\pm 0.007$ | 0.697<br>$\pm 0.010$ | 0.476<br>$\pm 0.011$ | 0.296<br>$\pm 0.006$ | 0.852<br>$\pm 0.007$ | 0.805<br>$\pm 0.007$ | 0.707<br>$\pm 0.010$ | 0.550<br>$\pm 0.011$ |
| ResNet+MCP | 0.832<br>$\pm 0.007$ | 0.730<br>$\pm 0.008$ | 0.496<br>$\pm 0.008$ | 0.306<br>$\pm 0.005$ | **0.895**<br>$\pm 0.007$ | **0.860**<br>$\pm 0.007$ | **0.758**<br>$\pm 0.006$ | **0.608**<br>$\pm 0.006$ |

**Table S14:** The normalized confusion matrix of the contact map prediction (cutoff=0.5).

| Dataset | Labels |  | Observation |  |
| --- | --- | --- | --- | --- |
|  |  |  | 0 | 1 |
| 327 test proteins | Prediction<br>(ResNet) | 0 | 0.993 | 0.662 |
|  |  | 1 | 0.007 | 0.338 |
|  | Prediction<br>(ResNet+MCP) | 0 | 0.995 | **0.595** |
|  |  | 1 | 0.005 | **0.405** |
| 495-protein dataset | Prediction<br>(ResNet) | 0 | 0.993 | 0.649 |
|  |  | 1 | 0.007 | 0.351 |
|  | Prediction<br>(ResNet+MCP) | 0 | 0.993 | **0.633** |
|  |  | 1 | 0.007 | **0.367** |

**Table S15:**
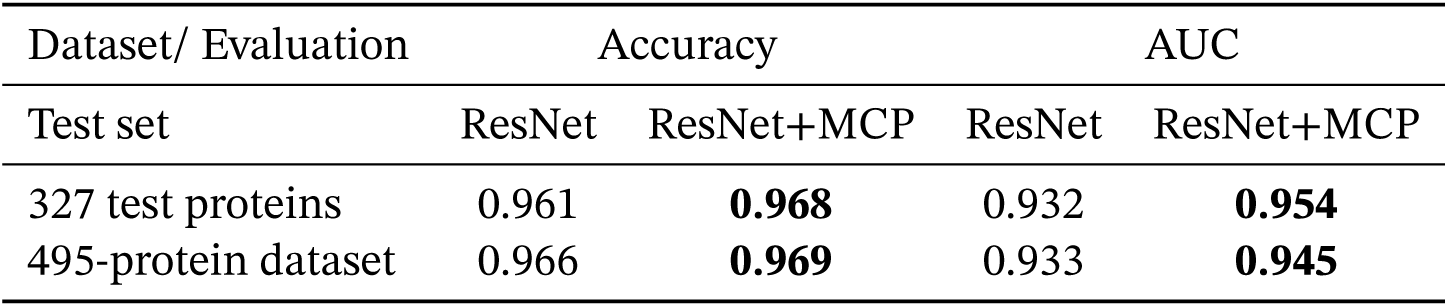
The accuracy (cutoff=0.5) and AUC of the contact map prediction.

### Supplementary Figures

**Figure S1:**
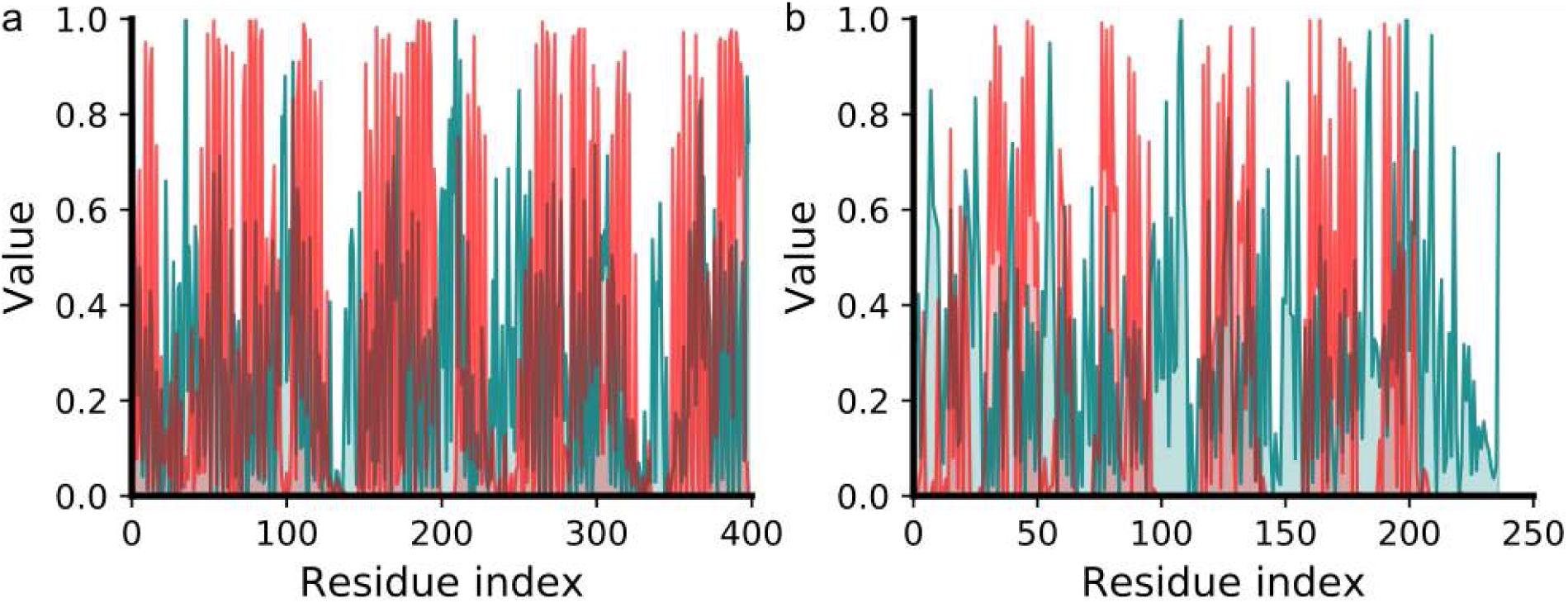
Comparison of the MCP prediction and the exposure (indicated by the relative solvent accessibility calculated by DSSP) for the two representative membrane proteins. The MCP and outer exposure results are shown as red and teal lines, respectively. According to DSSP [21], the buried residues are defined to have an RSA value of 0-0.1 [22, 23], so the outer surface residues have a RSA values of 0.1-1. We calculated the percentage of membrane-contacting residues with high MCP values (>0.5) lying in the outer residues with RSA > 0.1. The value is 84.0% for 5aym, and 88.9% for 4e1t. Therefore, most of the membrane-contacting residues predicted by MCP are outer-surface residues.

**Figure S2:**
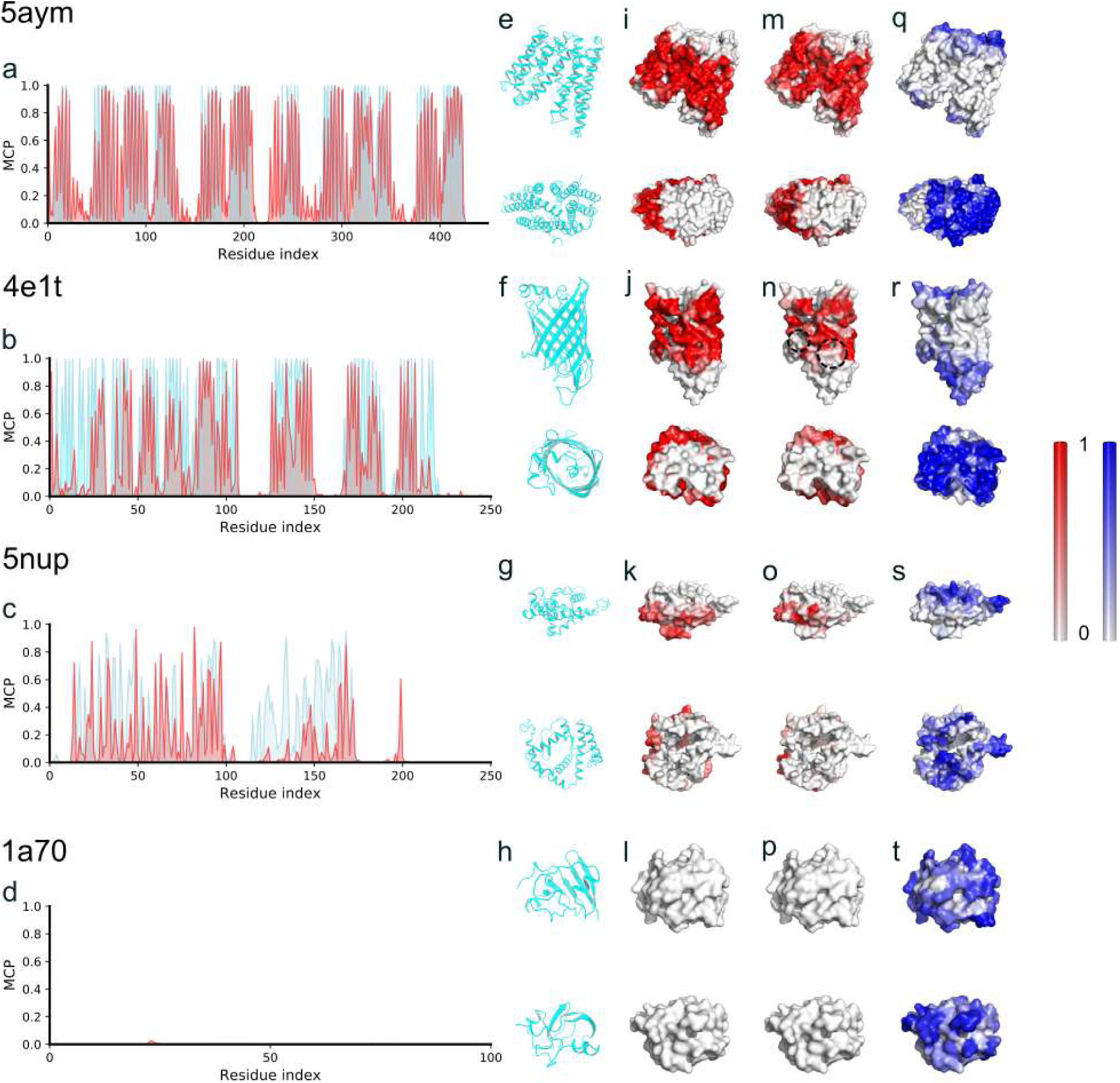
Membrane contact probability (MCP) of four representative proteins with the predictor trained by the MCP-Small dataset. (a-d) Comparison between the observation (cyan) and the prediction (red) of the MCPs. (e-h), Side and top views of the four representative proteins. (i-l), The outer surface of the representative proteins, colored according to the observed MCP values obtained from MD simulations. (m-p), Similar to (i-l), but colored according to the predicted MCP values. (q-t), Similar to (i-l), but colored according to the predicted SA values by RaptorX.

**Figure S3:**
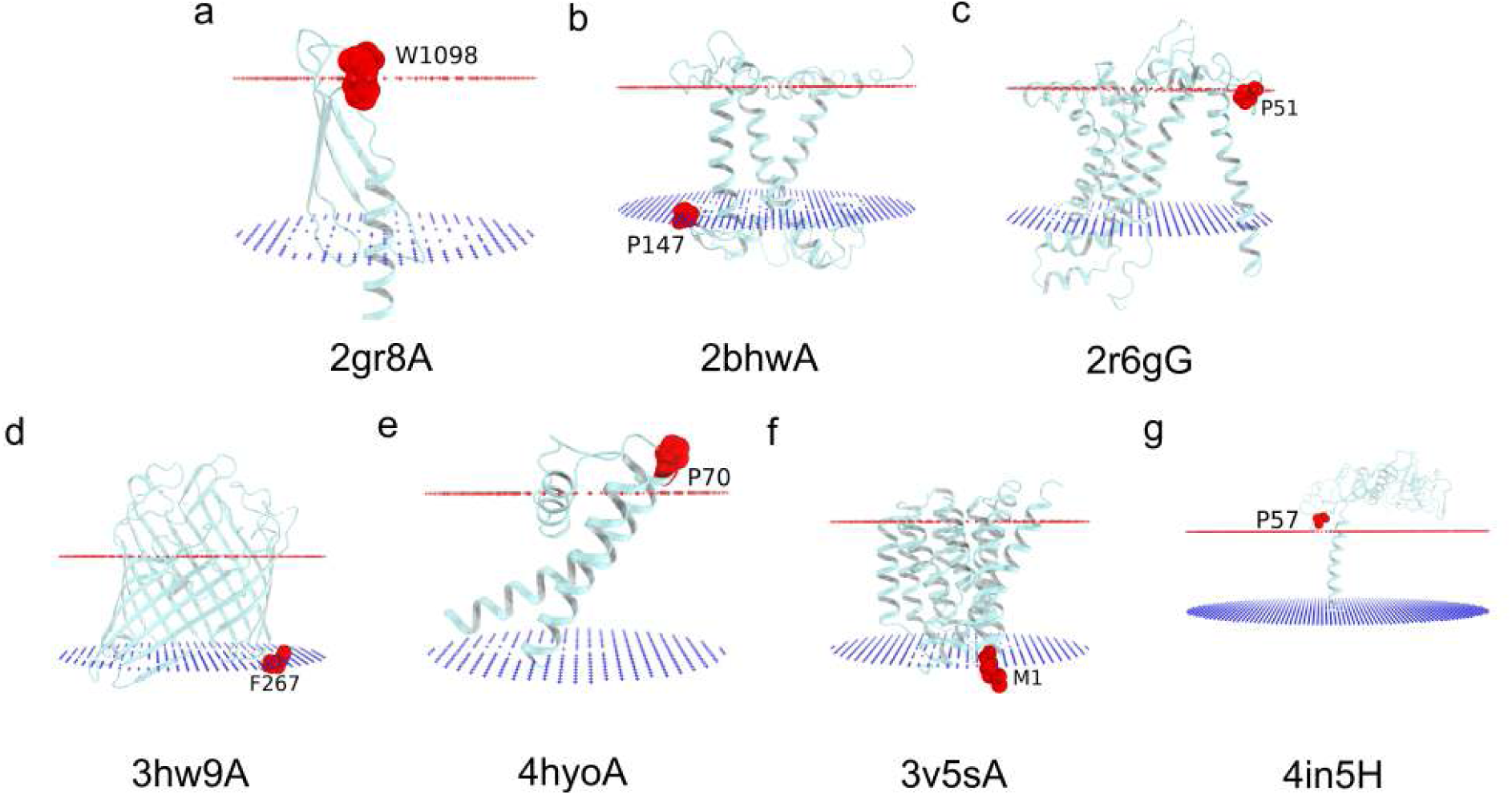
The structures of the membrane proteins showing where the residues in region I of Figure 3a are. The hydrophobic boundaries of the lipid bilayer are represented by the red and blue pseudoatoms, indicating the outer and inner surfaces of the bilayer, respectively.

**Figure S4:**
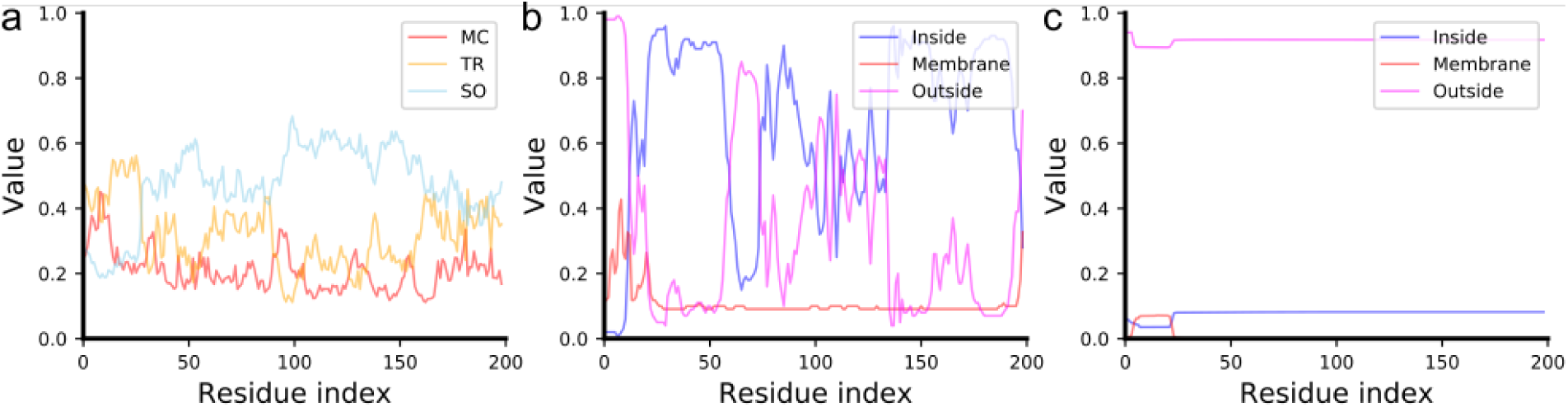
The prediction results of BCL::Jufo9D, TMHMM, and OCTOPUS for the Sar1 sequence. (a) The results of BCL::Jufo9D (red for membrane core (MC), sky blue for transition region (TR), and orange for solution (SO)). (b) The results of OCTOPUS (red for membrane, blue for inside, and fuchsia for outside). (c) The results of TMHMM (red for membrane, blue for inside, and fuchsia for outside).

**Figure S5:**
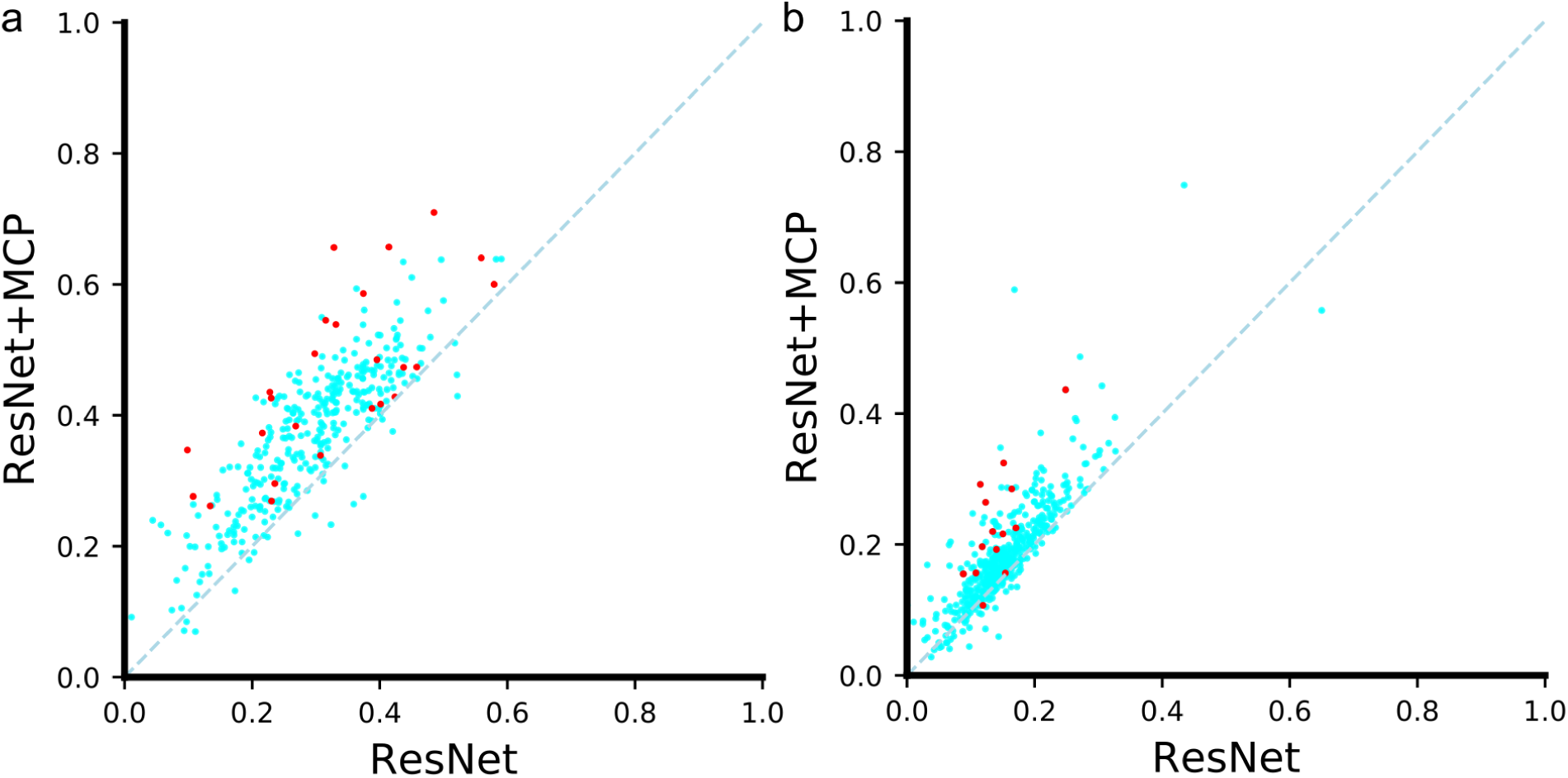
Comparison of the PCCs between the predictions and the native contact maps for the original ResNet predictor and our MCP-incorporated ResNet predictor. The original ResNet predictor (X-axis) vs our MCP-incorporated ResNet predictor (Y-axis) for the 327-protein dataset (a) and 495-protein dataset (b), respectively. The dashed line is the linear fit to the function *y* = *x*. Each point represents a test protein, with red points for membrane proteins.

**Figure S6:**
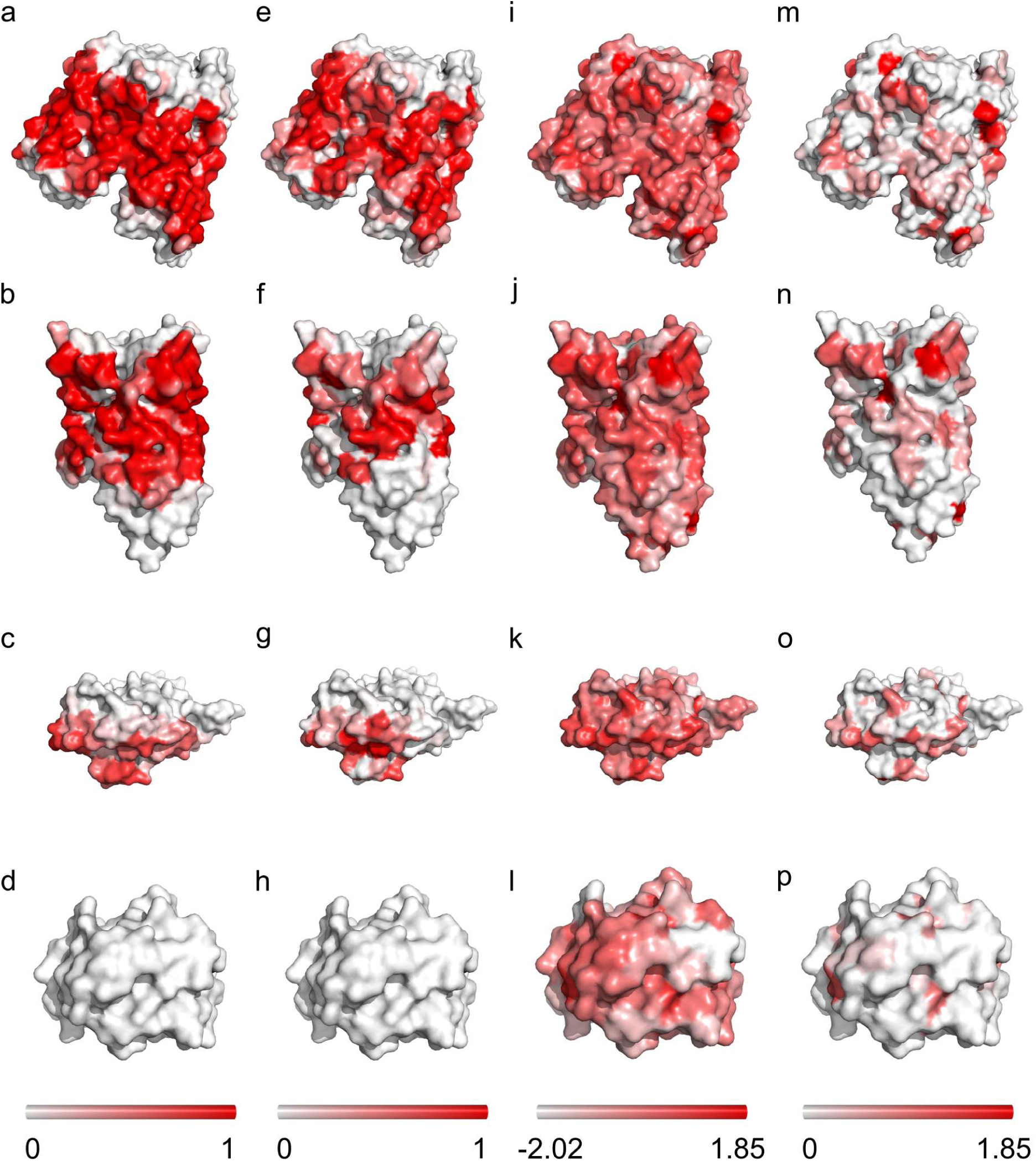
The protein surfaces colored according to the MCP and the Wimley-White hydrophobicity scales. (a-d), The colored outer surfaces of the four representative proteins presented in Fig. 1 & Fig. 3, according to the observed MCP values from MD simulations. (e-h), Similar to (a-d), but colored according to the predicted MCP values. (i-l), Similar to (a-b), but colored according to the value of the Wimley-White hydrophobicity scales. (m-p), Similar to (i-l), but with a different color bar.

**Figure S7:**
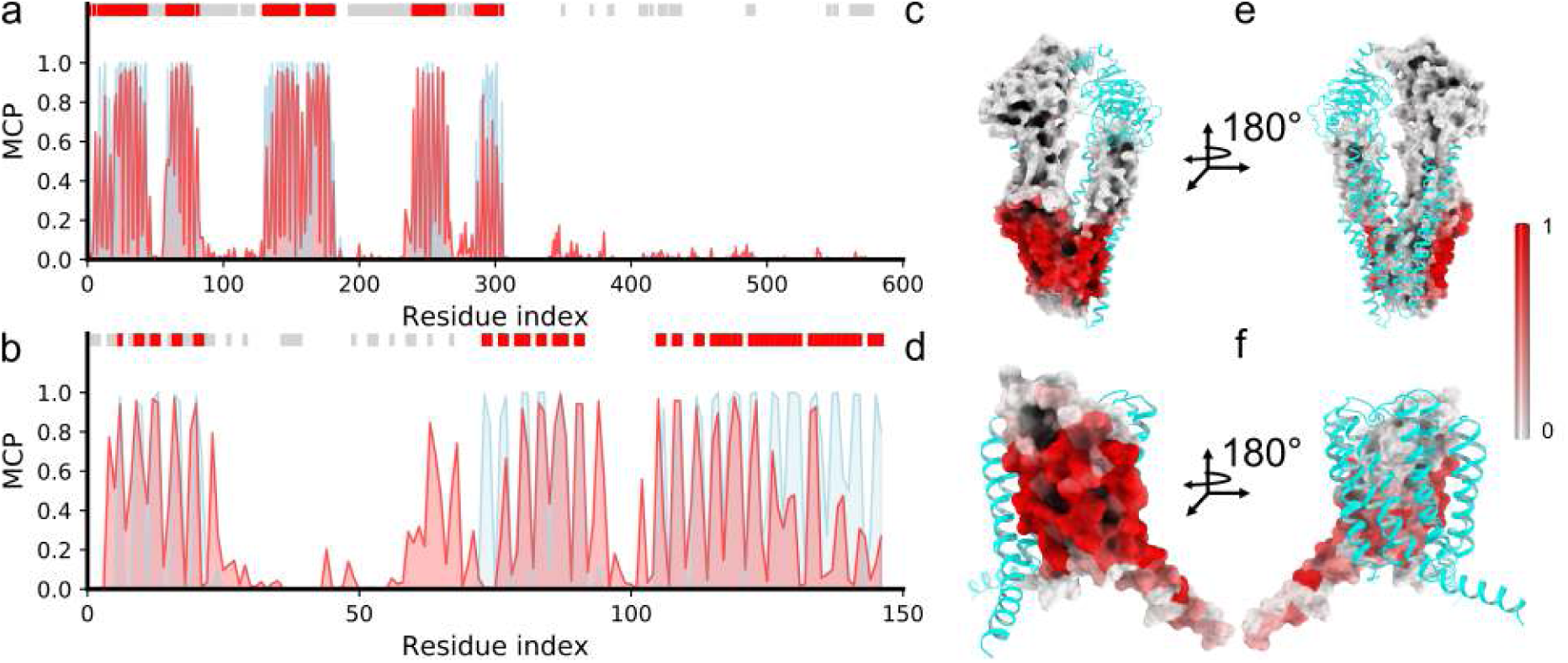
The MCP prediction results for two membrane-embedded complexes (PDB IDs: 5mkk and 2pno). The two complex proteins were in the test dataset with the overall prediction PCCs of 0.84 and 0.61, representing one of the good and one of the poor predictions, respectively. (a-b) Comparison between the observation (light blue) and the prediction (red) of MCP. The horizontal bars on the top of the panels indicate the regions of the protein-membrane (red) and protein-protein interfaces (gray). (c-d) The outer surface of the two proteins colored according to the predicted MCP values. (e-f) Similar to (c-d), but from another view. As can be seen, the protein-protein interface residues (gray bar) overall show low MCP values than those at the protein-membrane interfaces (red bar). The protein-membrane interface residues were defined by MCP > 0.2, while the protein-protein interface residues were defined with the script ‘InterfaceResidues’ of Pymol.

**Figure S8:**
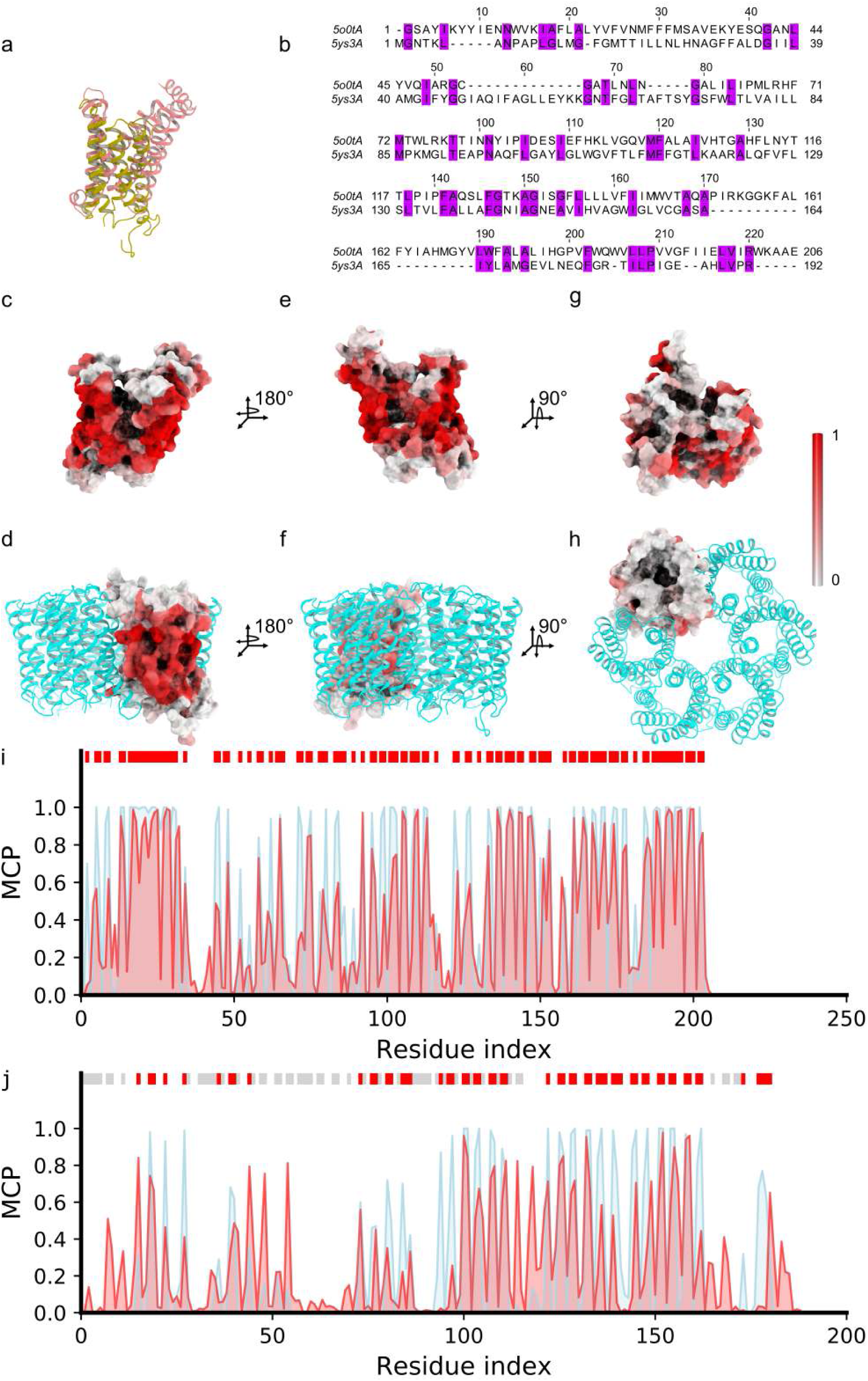
The MCP prediction results for two membrane proteins with similar folds but different oligo states, salmon for the monomeric protein (PDB ID: 5o0t) and olive for the oligomeric protein (PDB ID: 5ys3). (a) The structure alignment of the two proteins showed a similar fold (TM-score = 0.45, normalized by the length of 5ys3). (b) The sequence alignment of the two proteins (sequence similarity = 20.9%, calculated by MUSCLE [24]). (c-d) The outer surface of the two proteins colored according to the predicted MCP values. (e-f) Similar to (c-d), but from another side view. (g-h) Similar to (c-d), but from the top view.(i-j) Comparison between the MD observation (light blue) and the prediction (red) of MCP. The horizontal bars on the top of the panels indicate the regions of the protein-membrane (red) and protein-protein interfaces (gray). As can be seen, the protein-protein interface residues (gray bar) show overall low MCP values than those at the protein-membrane interfaces (red bar) in the transmembrane region. The protein-membrane interface residues were defined by MCP > 0.2, while the protein-protein interface residues were defined with the script ‘InterfaceResidues’ of Pymol.

**Figure S9:**
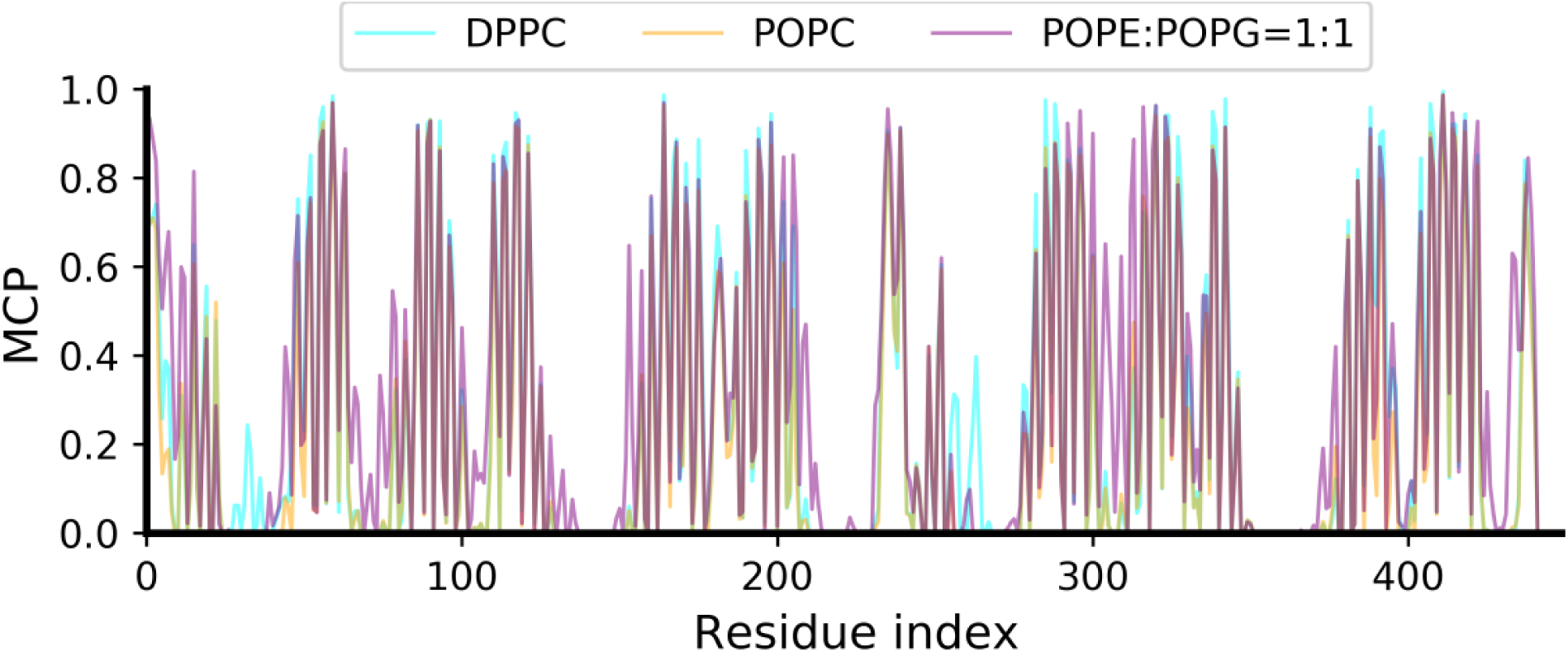
Comparison of the observed MCP in MD simulations with different types of lipids in membranes: cyan, orange, and purple lines for the membranes composed of DPPC (100%), POPC (100%), and mixed POPE (50%) and POPG (50%), respectively. These results show that the saturation, head group and net charge of the lipids have minor impacts on the observed MCP in the hydrophobic core region, and the differences are mostly located at the membrane-water interfaces.

**Figure S10:**
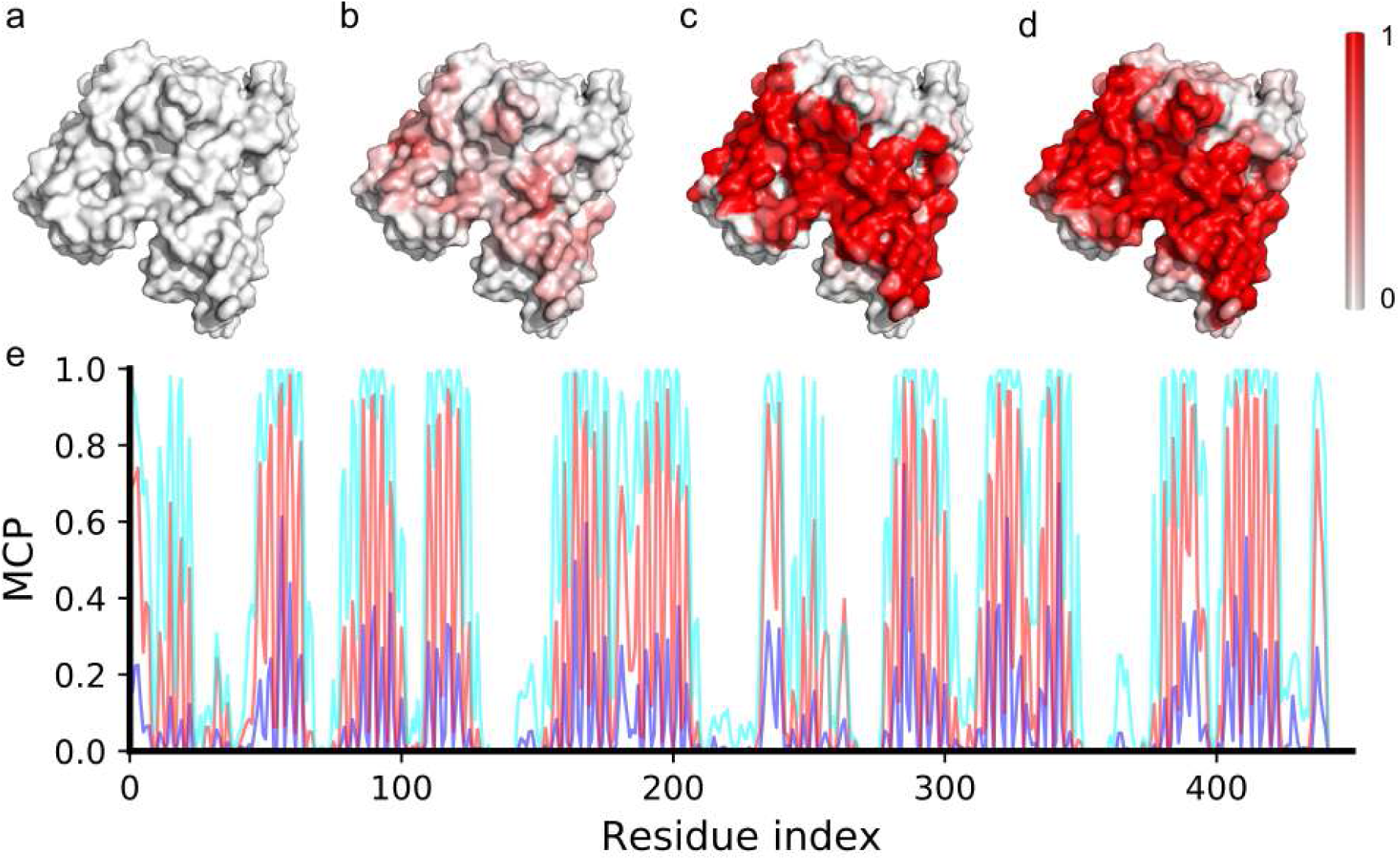
Comparison of the results of MCP analysis with different cutoff values and lipid types. (a-d), The colored outer surface of the protein (PDB ID: 5aym) according to the observed MCP values obtained from MD simulations with different cutoff values of 4, 5, 6, and 8 Å, respectively. As can be seen, a cutoff value smaller than 6 Å would lead to weak signals, while a cutoff value larger than 8 Å would start to overestimate the transmembrane region. (e), Comparison between the observed MCPs with different cutoff values; black, blue, red, and cyan lines for 4, 5, 6, and 8 Å, respectively.

**Figure S11:**
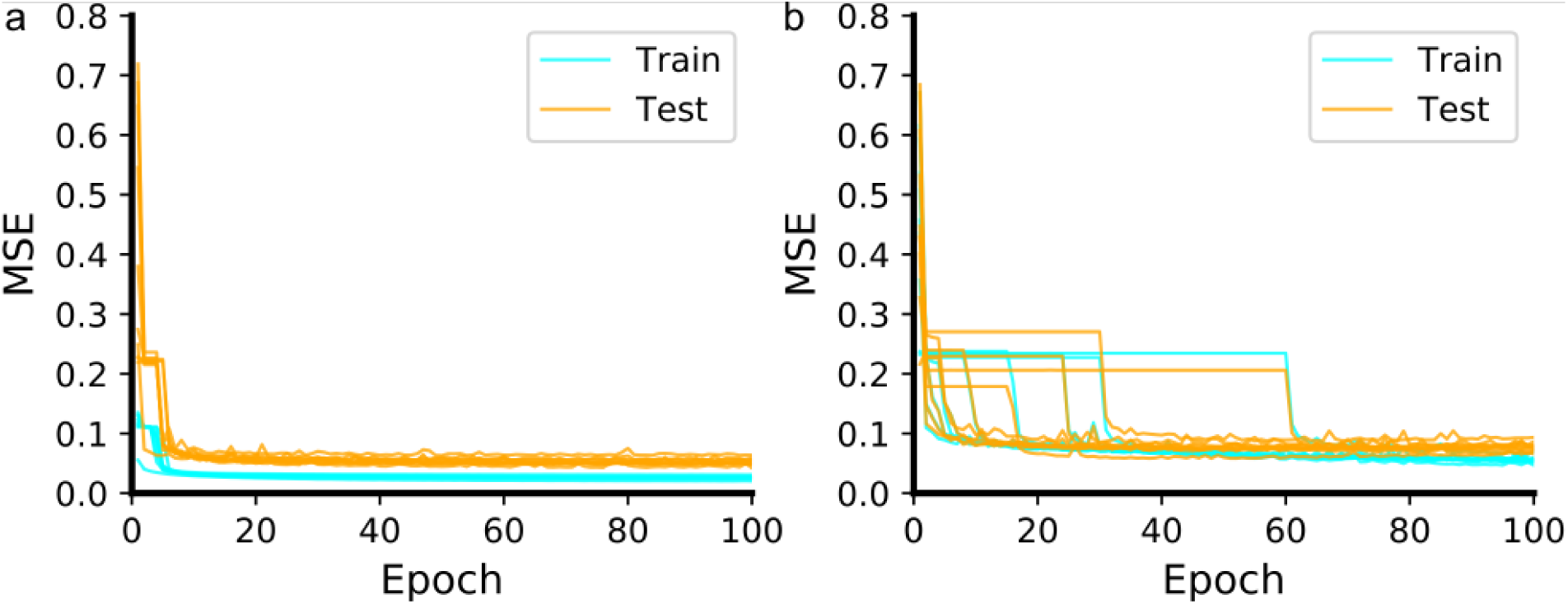
MSE curves of the MCP predictor in the 10-fold cross-validation. The left panel was obtained with the MCP-Large dataset, and the right panel with the MCP-Small dataset.

**Figure S12:**
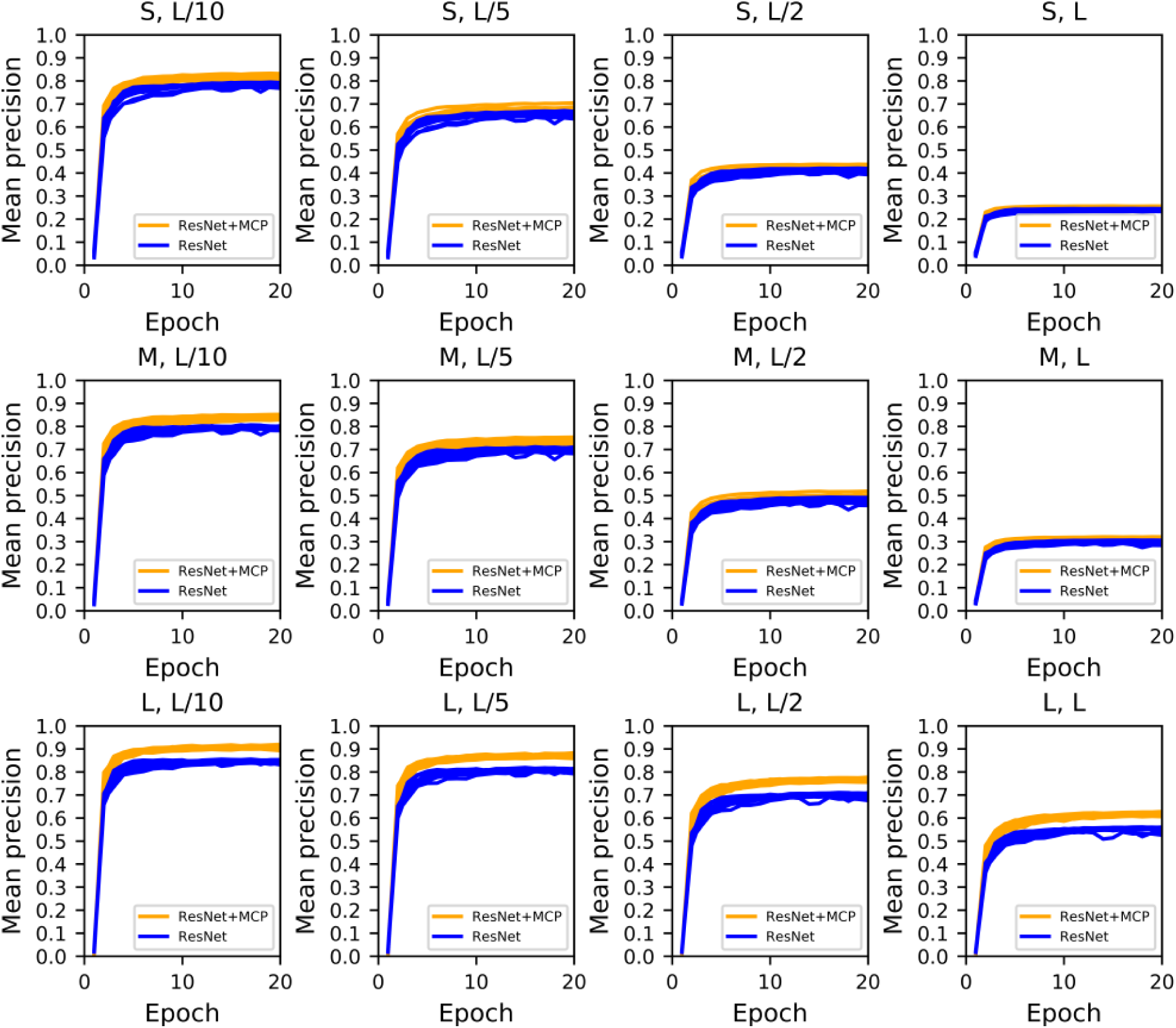
Precision curves in the test procedure of the contact map predictor in the 10-fold cross-validation: blue and orange lines for the results without and with MCP incorporated, respectively.

