## Supplementary material for "Membrane contact probability: an essential and predictive character for the structural and functional studies of membrane proteins": Dataset_Used_in_this_Work

### Dataset Used in the Work

#### 1. The large dataset for the MCP (Membrane Contact Probability) model

##### 1.1 The training set (5000 sequences)

1a0sP 1af6A 1afoA 1aigH 1aigL 1aigM 1aijH 1aijL 1aijM 1ap9A
1ar1A 1ar1B 1at9A 1ay2A 1b9uA 1bccB 1bccC 1bccD 1bccE 1bccF
1bccG 1bccH 1bccJ 1bctA 1be3A 1be3C 1be3E 1be3F 1be3G 1be3H
1be3I 1be3J 1bh3A 1bhaA 1bm1A 1brdA 1brrA 1brxA 1bt9A 1bxwA
1by5A 1c0vA 1c17A 1c17M 1c3wA 1c8rA 1c8sA 1c99A 1ds8H 1ds8L
1ds8M 1dv3L 1dv6H 1dv6L 1dv6M 1dx7A 1dxrC 1dxrM 1dxzA 1dzeA
1e0pA 1e12A 1e14M 1e54A 1e6dL 1e6dM 1e7pA 1e7pB 1e7pC 1ehkA
1ehkB 1ehkC 1ek9A 1eysC 1eysL 1eysM 1ezvA 1ezvB 1ezvC 1ezvD
1ezvE 1ezvF 1ezvG 1ezvH 1ezvI 1f4zA 1f6gA 1f6nH 1f6nM 1f88A
1fbbA 1fbkA 1fepA 1fftA 1fftB 1fftC 1fjkA 1fnpH 1fnpL 1fnpM
1fnqH 1fnqL 1fqyA 1fw2A 1fx8A 1g90A 1gfmA 1gfnA 1gfoA 1gfpA
1gfqA 1gu8A 1gzmA 1h68A 1h6iA 1h6s1 1hxtA 1hxuA 1hxxA 1i78A
1ih5A 1iijA 1ijdA 1ildA 1ilzA 1im0A 1iw6A 1iw9A 1iwgA 1izl0
1izlA 1izlB 1izlC 1izlD 1izlE 1izlK 1j95A 1jb0C 1jb0D 1jb0F
1jb0I 1jb0J 1jb0L 1jb0M 1jb0X 1jgjA 1jgwH 1jgwL 1jgwM 1jgxH
1jgxM 1jgyH 1jgyL 1jgyM 1jgzH 1jgzM 1jh0H 1jh0L 1jh0M 1jq1A
1jq2A 1jvmA 1k24A 1k4cC 1k4dC 1k6lH 1k6lL 1k6lM 1k6nH 1k6nL
1k6nM 1kb9A 1kb9D 1kb9E 1kb9G 1kb9H 1kb9I 1kbyH 1kbyL 1kbyM
1kf6A 1kf6B 1kf6C 1kf6D 1kfyB 1kfyC 1kfyD 1kg9A 1kgbA 1kmeA
1kmoA 1kmpA 1kpkA 1kplA 1kqfB 1kqfC 1kqgB 1kqgC 1kyoA 1kyoC
1kyoD 1kyoF 1kyoI 1l0lA 1l0lB 1l0lC 1l0lD 1l0lE 1l0lF 1l0lI
1l0lK 1l0nA 1l0nC 1l0nD 1l0nE 1l0nG 1l0nH 1l0nJ 1l0nK 1l0vB
1l0vC 1l0vD 1l6tA 1l7vC 1l9bC 1l9bH 1l9bL 1l9bM 1l9hA 1l9jL
1l9jM 1ldaA 1ldfA 1lghA 1lghB 1ln6A 1m0kA 1m0mA 1m3xH 1m3xL
1m3xM 1m56A 1m56B 1m56C 1m56D 1m57A 1m57B 1m57C 1m57D 1malA
1mgyA 1mm5A 1mpfA 1mpmA 1mpnA 1mpqA 1mpsH 1mpsL 1mpsM 1nekA
1nekB 1nekC 1nekD 1nenD 1nkzA 1nkzB 1nqeA 1nqfA 1nqgA 1nqhA
1ntkA 1ntkC 1ntkD 1ntkG 1ntkH 1ntkJ 1ntmC 1ntmD 1ntmE 1ntmF
1ntmH 1ntmJ 1ntmK 1ntzA 1ntzC 1ntzD 1ntzE 1ntzF 1ntzG 1ntzJ
1o0aA 1occA 1occB 1occC 1occD 1occE 1occF 1occG 1occH 1occI
1occK 1occL 1occM 1ocoA 1ocoB 1ocoD 1ocoG 1ocoI 1ocoK 1ocoL
1ocoM 1ocrA 1ocrB 1ocrD 1ocrG 1ocrH 1ocrI 1ocrK 1ocrL 1ocrM
1oczB 1oczC 1oczD 1oczG 1oczI 1oczJ 1oczL 1oczM 1oedA 1oedC
1ogvH 1oh2P 1oh2Q 1okcA 1opfA 1orsC 1osmA 1otsA 1ottA 1otuA
1oy6A 1oy8A 1oy9A 1oydA 1p49A 1p4tA 1p7bA 1p84A 1p84C 1p84D
1p84E 1p84G 1p84H 1p8iA 1pcrH 1pcrL 1pcrM 1phoA 1pnzA 1po0A
1pp9A 1pp9C 1pp9D 1pp9E 1pp9F 1pp9G 1pp9H 1pp9I 1pp9J 1prcC

1prcH 1prcM 1prnA 1pssM 1pstL 1pstM 1pv6A 1pv7A 1pw4A 1pxrA
1pxsA 1py6A 1q16B 1q16C 1q5iA 1q90A 1q90B 1q90C 1q90D 1q90G
1q90L 1q90N 1q90R 1q9fA 1qcrC 1qcrD 1qcrE 1qcrF 1qcrH 1qcrI
1qcrJ 1qcrK 1qd5A 1qd6C 1qhjA 1qj8A 1qj9A 1qjpA 1qkoA 1qkpA
1ql2A 1qlbA 1qlbC 1qleA 1qleB 1qleC 1qleD 1qm8A 1qo1A 1qo1D
1qo1G 1qo1J 1qo1K 1qovL 1r2cC 1r2cH 1r2cL 1r2cM 1r2nA 1r3iC
1r3kC 1r3lC 1r84A 1rc2A 1rg5L 1rg5M 1rgnH 1rgnL 1rgnM 1rh5A
1rh5B 1rh5C 1rhzA 1rhzB 1rhzC 1rqkL 1rqkM 1rvjH 1rvjL 1rwtA
1ry5H 1ry5M 1rzhH 1rzhM 1rzzH 1s00H 1s00L 1s00M 1s51A 1s52A
1s53A 1s5la 1s5lb 1s5ld 1s5le 1s5lh 1s5li 1s5lj 1s5lk 1s5ll
1s5lm 1s5lo 1s5lu 1s5lv 1s5lx 1s5lz 1s8lA 1siwC 1sorA 1sqbA
1sqbB 1sqbD 1sqbF 1sqbG 1sqbJ 1sqbK 1sqpA 1sqpC 1sqpD 1sqpE
1sqpF 1sqpG 1sqpJ 1sqpK 1sqqA 1sqqC 1sqqD 1sqqE 1sqqF 1sqqG
1sqqJ 1sqqK 1sqvA 1sqvC 1sqvD 1sqvE 1sqvF 1sqvG 1sqvJ 1sqvK
1sqxA 1sqxC 1sqxD 1sqxE 1sqxF 1sqxG 1sqxJ 1sqxK 1t1lA 1t9uA
1t9vA 1t9wA 1t9xA 1t9yA 1thqA 1tlwA 1tlyA 1tlzA 1tn0A 1tn5A
1tqqA 1u19A 1u77A 1u7cA 1u7gA 1uazA 1ucqA 1umxH 1umxL 1uynX
1uyoX 1v54A 1v54B 1v54C 1v54D 1v54E 1v54G 1v54I 1v54K 1v54L
1v54M 1v55A 1v55B 1v55C 1v55D 1v55G 1v55I 1v55J 1v55K 1v55M
1vf5A 1vf5B 1vf5C 1vf5D 1vf5E 1vf5F 1vf5G 1vf5H 1vgoA 1vjmA
1vrnH 1vrnL 1vrnM 1vryA 1wrgA 1x0i1 1x0k1 1x0sA 1xfhA 1xioA
1xjiA 1xkhA 1xl4A 1xl6A 1xmeB 1xmeC 1xrdA 1y4zB 1y4zC 1y5iB
1y5iC 1y5lB 1y5lC 1y5nB 1y5nC 1yc9A 1ycea 1yewA 1yf6H 1yf6L
1yf6M 1ymgA 1yq3A 1yq3B 1yq3C 1yq3D 1yq4B 1yq4C 1yq4D 1ystH
1ystL 1ystM 1z9jA 1z9jB 1z9jC 1z9kB 1z9kC 1zcdA 1zllA 1zoyA
1zoyB 1zoyC 1zp0C 1zp0D 1zrtC 1zrtD 1zrtE 1zzaA 2a06C 2a06D
2a06E 2a06F 2a06G 2a65A 2a9hA 2a9hE 2aczB 2aczC 2aczD 2akhC
2akiA 2akiB 2at9A 2axtb 2axtc 2axtd 2axte 2axtf 2axth 2axti
2axtj 2axtk 2axtl 2axtm 2axto 2axtt 2axtu 2b2fA 2b2iA 2b2jA
2b5fA 2b6oA 2b6pA 2b76A 2b76C 2bbjA 2bccA 2bccC 2bccD 2bccE
2bccF 2bccG 2bccJ 2begA 2bg9A 2bg9B 2bg9C 2bg9E 2bhwA 2bnpA
2bnpB 2bnsA 2bnsC 2bobC 2bozH 2bozM 2bs2A 2bs2B 2bs2C 2bs3C
2bs4C 2c0xA 2c32A 2cfpA 2cfqA 2cydA 2d2cA 2d2cB 2d2cE 2d2cF
2d2cG 2d57A 2db4A 2dhhA 2dr6A 2drdA 2dw3A 2dwdC 2dweC 2dyrA
2dyrB 2dyrC 2dyrD 2dyrG 2dyrI 2dyrJ 2dyrK 2dyrL 2dyrM 2dysA
2dysC 2dysD 2dysG 2dysI 2dysK 2dysL 2dysM 2e74A 2e74B 2e74C
2e74F 2e74G 2e75A 2e75B 2e75C 2e75D 2e75E 2e75F 2e75G 2e75H
2e76A 2e76B 2e76C 2e76D 2e76F 2e76G 2e76H 2ei4A 2eijA 2eijB
2eijC 2eijD 2eijI 2eijJ 2eijL 2eijM 2eikA 2eikB 2eikD 2eikG
2eikI 2eikJ 2eikK 2eikL 2eikM 2eilB 2eilC 2eilD 2eilG 2eilI
2eilJ 2eilL 2eilM 2eimA 2eimC 2eimD 2eimG 2eimI 2eimJ 2eimK
2eimL 2eimM 2einA 2einB 2einC 2einD 2einG 2einI 2einJ 2einK
2einL 2einM 2ervA 2evuA 2exyA 2ez0A 2f1cX 2f1tA 2f1vA 2f2bA
2f95A 2f95B 2fbwA 2fbwB 2fbwC 2fecA 2fedA 2feeA 2fgqX 2fkwB

2fynA 2fynB 2fynC 2fyuA 2fyuD 2fyuE 2fyuF 2fyuG 2fyuJ 2fyuK
2ge4A 2gmrH 2gmrM 2gnuH 2gnuM 2gr7A 2gr8A 2grxA 2grxC 2gskA
2gskB 2gsmA 2gsmB 2gufA 2h2pA 2h2sA 2h88A 2h88B 2h88C 2h89B
2h89C 2h89D 2h8aA 2hdfA 2hdiA 2hdiB 2hg3H 2hg3L 2hg3M 2hg9L
2hg9M 2hh1H 2hh1L 2hh1M 2hhkH 2hhkL 2hhkM 2hi2A 2hi7A 2hi7B
2hitH 2hitL 2hitM 2hj6L 2hj6M 2hjfC 2hpyA 2hqcA 2hqfA 2hqgA
2ht2A 2ht3A 2ht4A 2htkA 2hvkC 2hydA 2hynA 2i1xA 2i20A 2i21A
2i35A 2i37A 2i5nC 2i5nH 2i5nL 2i5nM 2i6wA 2ibzA 2ibzC 2ibzE
2ibzF 2ibzG 2ibzI 2ic8A 2irvA 2itaA 2itdC 2iubA 2iwvA 2iwwA
2j1nA 2j4uP 2j8cH 2j8cM 2j8dH 2j8dL 2j8dM 2j8sD 2jafA 2jagA
2jblC 2jblH 2jblL 2jblM 2jiyH 2jiyL 2jiyM 2jj0H 2jj0M 2jk4A
2jk5C 2jlnA 2jmmA 2jp3A 2jqyA 2jwaA 2k0lA 2k1aA 2k1kA 2k1lA
2k4tA 2k58B 2k73A 2k74A 2k8jX 2k9jA 2k9jB 2k9pA 2k9yA 2ka1A
2ka2A 2kdcA 2kihA 2kixA 2kluA 2klvA 2kncA 2kncB 2kpeA 2kpfA
2ks1A 2ks1B 2ks9A 2ksaA 2ksbA 2ksdA 2kseA 2ksfA 2ksjA 2ksrA
2ksyA 2kwxA 2kyhA 2kyvA 2l0jA 2l2tA 2l35A 2l35B 2l5bA 2l6wA
2l6xA 2l8sA 2l91A 2latA 2lckA 2lcxA 2legB 2lhfA 2lifA 2lj2A
2ljbA 2ljcA 2llmA 2llyA 2lm2A 2lnlA 2lohA 2lopA 2loqA 2lorA
2losA 2lp1A 2ltqA 2ly0A 2lz4A 2lzrA 2m06A 2m0bA 2m20A 2m3eA
2m3gA 2m59A 2m67A 2m6bA 2m6iA 2m6xA 2mawA 2mc7A 2metA 2meuA
2mgyA 2mjoA 2mjza 2mk9A 2mkaA 2mkvA 2mlhA 2mmuA 2mnhA 2mofA
2momA 2mozA 2mpnA 2mtsA 2mxbA 2n02A 2n1pA 2n2aA 2n2mA 2n4xA
2n5sA 2n70A 2n7qA 2n7rA 2n9yA 2na8A 2na9A 2ndjA 2nljC 2nmrA
2nopA 2nowA 2npcA 2npdA 2npeA 2npgA 2npjA 2npkA 2nq2A 2nq2C
2nr9A 2nrfA 2nrgA 2ntuA 2ntwA 2nuuA 2nuuG 2nwxA 2o011 2o012
2o013 2o014 2o01D 2o01E 2o01F 2o01G 2o01H 2o01I 2o01J 2o01L
2o01N 2o7lA 2o9dA 2o9eA 2o9gA 2oa0A 2oarA 2occA 2occB 2occC
2occG 2occI 2occM 2odjA 2omfA 2onjA 2onkA 2onkC 2onkE 2p7tC
2pnoA 2porA 2prcH 2prcL 2prcM 2prnA 2q67A 2q69A 2q6hA 2q72A
2q7mA 2q7rA 2qb4A 2qeiA 2qfiA 2qi9A 2qi9C 2qi9F 2qjkA 2qjkB
2qjkC 2qjpA 2qjpB 2qjpC 2qjuA 2qjyA 2qjyB 2qjyC 2qksA 2qpdA
2qpdB 2qpdC 2qpeA 2qpeC 2qtkA 2qtoA 2qtsA 2r4lA 2r4oA 2r4pA
2r4rA 2r4sA 2r6gA 2r6gF 2r6gG 2r89A 2r9hA 2r9rA 2r9rB 2rcrH
2rcrM 2rddA 2rlfA 2rmzA 2rn0A 2uuhA 2uuiA 2uwtH 2uwtL 2uwtM
2uwuH 2uwuL 2uwuM 2uwwH 2uwwL 2uwwM 2ux3H 2ux3M 2ux4L 2ux4M
2ux5L 2ux5M 2uxjH 2uxjL 2uxkH 2uxkL 2uxkM 2uxlH 2uxlL 2uxlM
2uxmH 2uxmL 2uxmM 2v8nA 2vdeA 2vdfA 2vl0A 2voyA 2voyB 2voyE
2voyF 2voyG 2voyH 2voyI 2voyJ 2voyK 2vpwB 2vpwC 2vpzC 2vqiA
2vt4A 2vv5A 2w0fC 2w1pA 2w2eA 2w5jA 2wdqB 2wdqC 2wdqD 2wdrB
2wdrC 2wdvB 2wdvC 2wdvD 2wgma 2wieA 2witA 2wjkA 2wjlA 2wjmC
2wjmH 2wjmL 2wjnC 2wjnM 2wjqA 2wjrA 2wlhA 2wliB 2wljA 2wlkA
2wllA 2wllB 2wlnA 2wp9B 2wp9C 2wp9D 2wpdA 2wpdD 2wpdH 2wpdI
2wqyB 2wqyC 2wqyD 2ws3B 2ws3C 2wsc1 2wsc2 2wsc3 2wsc4 2wscA
2wscC 2wscE 2wscG 2wscH 2wscI 2wscJ 2wscK 2wscL 2wscN 2wse1

2wse2 2wse3 2wse4 2wseA 2wseF 2wseG 2wseH 2wseJ 2wseK 2wseL
2wseN 2wsf1 2wsf2 2wsf3 2wsf4 2wsfA 2wsfF 2wsfH 2wsfI 2wsfK
2wsfN 2wswA 2wsxA 2wu2B 2wu2C 2wu2D 2wu5B 2wu5C 2wu5D 2wvpA
2ww9A 2ww9B 2ww9H 2ww9I 2ww9J 2ww9K 2ww9L 2ww9N 2ww9O 2wwbB
2wwbC 2wwbJ 2wx5H 2wx5L 2wx5M 2x27X 2x2vA 2x55A 2x5uC 2x5uH
2x5uL 2x5uM 2x5vC 2x5vM 2x6aA 2x6bA 2x6cA 2x72A 2x79A 2x9kA
2xe1A 2xe2A 2xe3A 2xe5A 2xg6A 2xkmA 2xmnA 2xokD 2xokG 2xokH
2xokK 2xovA 2xowA 2xq2A 2xq3A 2xq4A 2xq5A 2xq6A 2xq7A 2xq8A
2xq9A 2xqaA 2xqsA 2xqtA 2xquA 2xtuA 2xtvA 2y01A 2y02A 2y03A
2y04A 2y0hA 2y0kA 2y0lA 2y2xA 2y5yA 2y69A 2y69B 2y69C 2y69D
2y69I 2y69J 2y69K 2y69L 2y69M 2ycwA 2ycxA 2ycyA 2yczA 2ydoA
2ydvA 2yevB 2yevC 2yiuA 2yiuB 2yiuC 2yn6A 2yn9B 2ynkA 2yoeA
2yptA 2ysuA 2ysuB 2yvxA 2yxqA 2yxrA 2z55A 2zd9A 2zfeA 2zfgA
2ziyA 2zldA 2zqpE 2zt9A 2zt9B 2zt9E 2zt9F 2zt9G 2zt9H 2zupB
2zuqA 2zw3A 2zxeB 2zxeG 2zxwA 2zxwB 2zxwC 2zxwD 2zxwF 2zxwG
2zxwI 2zxwJ 2zxwK 2zxwM 2zzlA 3a0ba 3a0bb 3a0bc 3a0bd 3a0be
3a0bf 3a0bh 3a0bk 3a0bl 3a0bm 3a0bt 3a0bx 3a0bz 3a2sX 3a3yB
3a3yG 3a7kA 3abkB 3abkG 3abkI 3abkJ 3abkK 3abkL 3abkM 3ablA
3ablB 3ablC 3ablD 3ablI 3ablK 3ablL 3ablM 3abmA 3abmB 3abmD
3abmG 3abmI 3abmJ 3abmK 3abmL 3abvB 3abvC 3abvD 3abwA 3ae1D
3ae2D 3ae3C 3ae3D 3ae4B 3ae4C 3ae5B 3ae5C 3ae5D 3ae6B 3ae6C
3ae6D 3ae7B 3ae7C 3ae7D 3ae8B 3ae8D 3ae9B 3aeaB 3aeaC 3aeaD
3aebB 3aebC 3aebD 3aecB 3aecC 3aedB 3aedC 3aedD 3aeeB 3aeeD
3aefB 3aefC 3aefD 3aegB 3aegD 3aehA 3ag1A 3ag1B 3ag1C 3ag1D
3ag1G 3ag1I 3ag1J 3ag1L 3ag1M 3ag2A 3ag2B 3ag2C 3ag2D 3ag2I
3ag2J 3ag2K 3ag2L 3ag2M 3ag3A 3ag3D 3ag3G 3ag3J 3ag3K 3ag3L
3ag3M 3ag4A 3ag4B 3ag4C 3ag4D 3ag4G 3ag4I 3ag4J 3ag4K 3ag4L
3ag4M 3am6A 3anza 3aoaA 3aobA 3aocA 3aouA 3aqpA 3asnA 3asnB
3asnD 3asnG 3asnI 3asnJ 3asnK 3asnL 3asoA 3asoB 3asoC 3asoD
3asoF 3asoG 3asoI 3asoJ 3asoK 3asoL 3asoM 3ayfA 3aymA 3aynA
3b29A 3b44A 3b45A 3b4rA 3b5wA 3b5xA 3b5yA 3b5zA 3b61A 3b8eB
3b8eG 3b9wA 3b9yA 3ba6A 3bccA 3bccD 3bccE 3bccF 3bccG 3bccJ
3behA 3bhsA 3bo0A 3bo0B 3bo0C 3bo1A 3bryA 3brzA 3bs0A 3bvdA
3bvdB 3bvdC 3c02A 3c1gA 3c1iA 3c9mA 3capA 3chxA 3chxB 3cirA
3cirB 3cirC 3cllA 3cn5A 3cn6A 3cocA 3codA 3cslC 3cwbA 3cwbB
3cwbC 3cwbE 3cwbF 3cwbH 3cwbJ 3cx5C 3cx5D 3cx5G 3cx5H 3cx5I
3cxhC 3cxhD 3cxhE 3cxhG 3cxhH 3d31A 3d31C 3d38C 3d38H 3d38L
3d38M 3d4sA 3d5kA 3d9sA 3ddlA 3ddrC 3dhwA 3dhwC 3dinC 3dinD
3dknB 3dqbA 3dsyL 3dsyM 3dtaH 3dtaL 3dtaM 3dtrH 3dtrL 3dtrM
3dtsH 3dtsL 3dtsM 3dtuA 3dtuB 3du2H 3du2L 3du2M 3du3M 3duqL
3duqM 3dwnA 3dwwA 3dzmA 3e83A 3e86A 3e89A 3e8fA 3e8gA 3e8hA
3efmA 3egwB 3egwC 3eh3A 3eh3B 3eh3C 3eh4A 3eh4B 3eh4C 3eh5A
3eh5B 3eh5C 3ehbA 3ehbB 3ehzA 3ejyA 3ejzA 3emlA 3emnX 3emoA
3f3cA 3f3dA 3f3eA 3f4iA 3f4jA 3f7yC 3fb6C 3fh6A 3fh6F 3fh6G

3fhhA 3fi1A 3fidA 3fipA 3fyeA 3fyiA 3fyiB 3fyxA 3g5uA 3g60A
3g7fC 3g7fH 3g7fL 3g7fM 3gb7C 3gd8A 3gi8C 3giaA 3gjcA 3gjdA
3gp6A 3gwuA 3gwvA 3gwwA 3h1hD 3h1hE 3h1hF 3h1hG 3h1hJ 3h1iA
3h1iC 3h1iD 3h1iE 3h1iF 3h1iG 3h1iJ 3h1jA 3h1jC 3h1jD 3h1jE
3h1jF 3h1jJ 3h1kA 3h1kB 3h1kC 3h1kD 3h1kE 3h1kF 3h1kG 3h1kH
3h1kJ 3h1lD 3h1lE 3h1lF 3h1lG 3h1lJ 3h90A 3h9vA 3hanA 3haoA
3haqA 3harA 3hb3A 3hb3B 3hd6A 3hd7A 3hd7C 3hd7D 3hfxA 3hw9A
3hwbA 3hzqA 3i4dH 3i4dL 3i4dM 3i5dA 3ifxA 3ij4A 3ipdA 3ipdB
3ir5B 3ir6A 3ir6B 3ir6C 3ir7B 3iyzA 3iz1A 3iz2A 3j08A 3j41A
3j41E 3j45E 3j45G 3j45T 3j45U 3j45Y 3j5pA 3j5rA 3j9jA 3j9pA
3jacA 3jadA 3jaeA 3jafA 3jc21 3jc22 3jcfA 3jtyA 3jycA 3k03A
3k04A 3k06A 3k07A 3k08A 3k0dA 3k0iA 3k19A 3k3fA 3k3gA 3kbcA
3kcuA 3kcvA 3kdpB 3kdpG 3kg2A 3kj6A 3klyA 3klzA 3kp9A 3ksoA
3kvnA 3kziA 3kziB 3kziC 3kziD 3kziE 3kziF 3kziH 3kziI 3kziK
3kziL 3kziM 3kziO 3kziT 3kziU 3kziX 3kziy 3kziZ 3l1lA 3l72A
3l72C 3l72D 3l72F 3l72G 3l72J 3l74A 3l74C 3l74D 3l74F 3l74G
3l74J 3l75A 3l75C 3l75D 3l75E 3l75F 3l75J 3lddA 3leoA 3llqA
3lrbA 3lrcA 3lsvA 3lw51 3lw52 3lw53 3lw54 3lw5A 3lw5D 3lw5E
3lw5F 3lw5G 3lw5H 3lw5I 3lw5J 3lw5K 3lw5L 3m4eA 3m6eA 3m71A
3m72A 3m73A 3m74A 3m76A 3m77A 3m7bA 3m7cA 3m7eA 3m7lA 3m8bA
3m8dA 3m9iA 3mbvA 3me1A 3mk7A 3mk7C 3mktA 3mkuA 3mp7A 3mp7B
3mpnA 3mpqA 3n23A 3n23B 3ncyA 3nd0A 3ne2A 3ne5A 3ne5B 3njtA
3nk5A 3nmoA 3nocD 3nogD 3ns0A 3nsgA 3ny9A 3nyaA 3o0rC 3o7pA
3oduA 3oe0A 3oe8A 3oe9A 3ogcC 3ohnA 3om3B 3omaA 3omaB 3omiA
3omiB 3omnA 3or6C 3orgA 3oufA 3ousA 3p03A 3p0gA 3p4pB 3p4pC
3p4pD 3p4qB 3p4qC 3p4qD 3p4rB 3p4rC 3p4rD 3p4sB 3p4sD 3p4wA
3p50A 3p5nA 3pblA 3pcqA 3pcqI 3pcqJ 3pcqL 3pcqM 3pcqX 3pdsA
3pf1A 3pgrA 3pguA 3pikA 3pjsK 3pl9A 3pouA 3poxA 3pqrA 3prnA
3puvA 3puvE 3puvF 3puvG 3puwF 3puwG 3puxA 3puxG 3puyA 3puyF
3puyG 3pv0A 3pv0E 3pv0F 3pv0G 3pwhA 3q17A 3q7kA 3qapA 3qbgA
3qbkA 3qblA 3qdcA 3qe7A 3qf4A 3qf4B 3qjqA 3qjqB 3qjrA 3qjrB
3qjrC 3qjsA 3qjsB 3qjsC 3qjtA 3qjtB 3qjtC 3qjuA 3qjuB 3qjuC
3qjvC 3qlbA 3qnqA 3qq2A 3qraA 3qrcA 3qs4A 3qs6A 3r65A 3rceA
3reyA 3rfmA 3rfrA 3rfrB 3rfrC 3rfuA 3rfzA 3rfzB 3rfzC 3rgbA
3rgbB 3rgmA 3rgnA 3ri5A 3riaA 3rifA 3rkoC 3rkoD 3rkoF 3rkoG
3rlbA 3rlfF 3rlfG 3rquA 3rqwA 3rvyA 3rvzA 3rzeA 3s33A 3s33B
3s38B 3s39A 3s39B 3s39C 3s3aA 3s3aC 3s3bA 3s3bB 3s3bC 3s3cB
3s3cC 3s3dA 3s3dB 3s3dC 3s3wA 3s3xA 3s3xD 3s8fA 3s8fB 3s8fC
3s8gA 3s8gC 3sfdB 3sfdC 3sfdD 3sfeB 3sfeC 3sfeD 3sljA 3sloA
3sltA 3sn6A 3sn6B 3sn6G 3spcA 3spgA 3sphA 3spiA 3spjA 3stlC
3sy7A 3sy9A 3sybA 3sycA 3syoA 3sypA 3syqA 3sysA 3szdA 3t0sA
3t1cA 3t20A 3t24A 3t2mA 3t4dA 3t4zA 3t51A 3t51B 3t53A 3t56A
3t6dC 3t6dH 3t6dL 3t6dM 3t6eC 3t6eH 3t6eL 3t6eM 3t9nA 3tcuA
3tdpA 3tdrA 3tdsA 3te0A 3te2A 3tetA 3tguA 3tguC 3tguD 3tguE

3tguF 3tguG 3tguI 3tguJ 3tlsA 3tltA 3tluA 3tlvA 3tt1A 3tt3A
3tu0A 3tuiA 3tuiC 3tujA 3tujC 3tuzA 3tuzC 3tx3A 3u2fK 3u2yK
3u32K 3ubbA 3ud0K 3udcA 3ug9A 3ukmA 3um7A 3uonA 3uq4A 3usgA
3usiA 3usjA 3uskA 3uslA 3usmA 3usoA 3uspA 3utvA 3utwA 3utxA
3uu2A 3uu4A 3uu6A 3uu8A 3uubA 3ux4A 3uzaA 3uzcA 3v2wA 3v2yA
3v3yH 3v3yL 3v3yM 3v3zH 3v3zL 3v3zM 3v5sA 3v5uA 3v89A 3v89B
3v8fA 3v8xA 3v8xB 3vg9A 3vhzA 3vmrA 3vmsA 3vmtA 3vr8A 3vr8B
3vr8C 3vr8D 3vr9B 3vr9D 3vraB 3vraC 3vraD 3vrbB 3vrbC 3vvkA
3vvnA 3vvoA 3vvpA 3vvrA 3vw7A 3vy8X 3vztX 3vzuX 3vzwX 3w4tA
3w5aC 3w9tA 3wajA 3wbnA 3wdoA 3wfbB 3wfbC 3wfcB 3wfcC 3wfdC
3wfeC 3wg7A 3wg7B 3wg7C 3wg7D 3wg7G 3wg7I 3wg7J 3wg7K 3wg7L
3wg7M 3wguB 3wguE 3wgvB 3wgvE 3wi5A 3wmeA 3wmfA 3wmgA 3wmm1
3wmmC 3wmmL 3wmmM 3wo6A 3wqjA 3wu2a 3wu2b 3wu2c 3wu2d 3wu2e
3wu2f 3wu2h 3wu2j 3wu2k 3wu2l 3wu2m 3wu2o 3wu2x 3wu2y 3wu2z
3wvfA 3wxvA 3wxwA 3x29A 3x29B 3x2qA 3x2qC 3x2qD 3x2qG 3x2qI
3x2qJ 3x2qK 3x2qL 3x2qM 3x2rA 3x3bA 3x3cA 3zd0A 3zdqA 3ze3A
3ze4A 3ze5A 3zevA 3zjzA 3zk1A 3zk2A 3zkrA 3zmiA 3zmjA 3zo6A
3zojA 3zotA 3zpqA 3zrsA 3zumH 3zumL 3zumM 3zuwH 3zuwM 3zuxA
3zuyA 4a01A 4a2nB 4a4mA 4a82A 4a97A 4a98A 4ac5C 4ac5H 4ac5M
4ageA 4agfA 4aipA 4aiqA 4al0A 4amiA 4apsA 4aq5A 4aq5B 4aq5E
4aq9A 4aq9B 4aq9C 4aq9E 4atvA 4av3A 4av6A 4aw6A 4aytA 4aywA
4ayxA 4azlA 4b61A 4b7oA 4bbjA 4bemA 4bemJ 4bevA 4bezA 4bgnA
4boiA 4boiB 4boiC 4boiE 4booC 4booE 4borA 4borB 4borC 4borE
4botC 4botE 4bpdA 4bpmA 4brbA 4brrA 4bumX 4buoA 4bv0A 4bvnA
4bw5A 4bwbA 4bwzA 4bygA 4c48A 4c48B 4c48C 4c4vA 4c69X 4c7rA
4c9jA 4c9qA 4cadC 4casA 4casB 4casC 4casD 4cbcA 4cbjA 4cbkA
4cdiA 4cdiC 4cg5A 4cg5B 4cg5C 4cg6A 4cg6B 4cg6C 4cg7B 4chvA
4cjzA 4ck0A 4cskA 4cy4A 4czaA 4czbA 4czbC 4d0aB 4d1aA 4d1bA
4d1cA 4d1dA 4d2bA 4d2cA 4d2dA 4d51A 4d5dA 4d64A 4d6tA 4d6tB
4d6tC 4d6tD 4d6tE 4d6tF 4d6tG 4d6tH 4d6tJ 4d6tO 4d6uA 4d6uC
4d6uE 4d6uF 4d6uG 4d6uN 4dajA 4dblA 4dblC 4dblE 4dcbA 4djhA
4djiA 4djkA 4dklA 4dnrA 4dntA 4dojA 4dopA 4dveA 4dw0A 4dw1A
4dxwA 4eebA 4eedA 4eiyA 4ej4A 4ekwA 4epaA 4ev6A 4ezcA 4ezdA
4f35A 4f4cA 4f4lA 4f4sA 4f8hA 4fa7A 4fa7B 4fa7C 4faaB 4faaC
4fbzA 4fc4A 4fe1I 4fe1J 4fe1L 4fe1M 4fe1X 4fg6A 4fi3A 4fmsA
4fozA 4fpdA 4fqeA 4frtA 4fsoA 4ft6A 4ftpA 4fuvA 4fxzA 4fy0A
4g1uA 4g70A 4g70B 4g70C 4g71A 4g71B 4g71C 4g72C 4g7qA 4g7qB
4g7qC 4g7rC 4g7sA 4g7sB 4g7vS 4g7yS 4g80I 4gbyA 4gbzA 4gc0A
4gd3A 4gd3J 4gd3Q 4geyA 4gf4A 4gp4A 4gp4C 4gp5A 4gp5B 4gp5C
4gp8A 4gp8C 4gpoA 4gx0A 4gx1A 4gx5A 4gycA 4gycB 4h0lA 4h0lB
4h0lC 4h0lD 4h0lE 4h0lF 4h0lG 4h0lH 4h13A 4h13C 4h13E 4h13F
4h1dA 4h1wB 4h33A 4h37A 4h44A 4h44B 4h44C 4h44D 4h44E 4h44F
4h44G 4h56A 4h99H 4h99M 4h9lH 4h9lM 4hbhH 4hbhL 4hbjH 4hbjL
4hbjM 4he8A 4he8C 4he8D 4he8E 4he8F 4he8G 4he8I 4hea2 4hea4

4hea5 4hea6 4hea7 4hea9 4heaA 4heaH 4heaJ 4heaK 4heaL 4heaM
4heaN 4heaW 4hfbA 4hfcA 4hfdA 4hfeA 4hfhA 4hfiA 4hg6B 4hkrA
4hksA 4hodA 4hqjB 4hqjE 4htsA 4hukB 4hulA 4humA 4hunA 4huqA
4huqB 4huqS 4huqT 4hw9A 4hwaA 4hwlA 4hycA 4hydA 4hygA 4hyjA
4hyoA 4hytA 4hytB 4hytE 4hyxA 4hz3A 4hzuA 4hzuS 4i0uA 4i7zA
4i7zB 4i7zC 4i7zD 4i7zE 4i7zF 4i7zG 4i7zH 4i9wA 4ia4A 4ib4A
4ikvA 4ikwA 4ikxA 4ikyA 4il3A 4il4A 4il6a 4il6b 4il6c 4il6d
4il6e 4il6f 4il6j 4il6k 4il6l 4il6m 4il6R 4il6t 4il6x 4il6y
4il6z 4il9A 4ilbA 4ilcA 4in5H 4in5L 4in5M 4in6H 4in6L 4in6M
4in7L 4in7M 4ireA 4iu9A 4ixqb 4ixqc 4ixqd 4ixqe 4ixqf 4ixqg
4ixqh 4ixqi 4ixqj 4ixqk 4ixql 4ixqm 4ixqo 4ixqt 4ixqx 4ixqz
4ixra 4ixrb 4ixrc 4ixrd 4ixre 4ixrf 4ixrg 4ixri 4ixrj 4ixrk
4ixrl 4ixrm 4ixrt 4ixrx 4ixrz 4j4qA 4j72A 4j7cA 4j7cI 4j7yA
4j9uA 4j9uE 4ja3A 4ja4A 4jbwA 4jbwF 4jbwG 4jbwM 4jc6A 4jc7A
4jfbA 4jkvA 4jq6A 4jr8A 4jr9A 4jreA 4jrzA 4jtcB 4jtdB 4k0jA
4k1cA 4k3cA 4k5yA 4k7qA 4k7rA 4kfmA 4kfmB 4khzA 4khzG 4ki0A
4ki0G 4kjpA 4kjqA 4kjrA 4kjsA 4kjwA 4kk5A 4kk6A 4kk8A 4kkaA
4kkbA 4klyA 4knfA 4kppA 4kr4A 4kraA 4ksbA 4kscA 4ksdA 4kt0D
4kt0F 4kt0K 4kt0M 4kx6B 4kx6D 4kytB 4l6v0 4l6v6 4l6v7 4l6v8
4l6v9 4lbeC 4lczA 4ldsA 4lepA 4llhA 4lmjA 4lmkA 4lmlA 4louA
4lp8A 4lseA 4lsgA 4lshA 4lsiA 4ltoA 4ltpA 4ltqA 4ltrA 4lwyH
4lwyL 4lwyM 4lxjA 4lz6A 4lz9A 4m1mA 4m2tA 4m58A 4m5bA 4m5cA
4mbsA 4md1A 4md2A 4mesA 4mhwA 4mlbA 4mm4A 4mm5A 4mm6A 4mm7A
4mm8A 4mm9A 4mmbA 4mmcA 4mmdA 4mmeA 4mmfA 4mndA 4mqsA 4mqtA
4mqxA 4mrnA 4mrpA 4ms2A 4mt0A 4mt1A 4mt4A 4mtgA 4mtoA 4muuA
4mvmA 4mvoA 4mvqA 4mvrA 4mvsA 4mvuA 4mvzA 4mw3A 4mw8A 4myhA
4n4rA 4n4wA 4n4yB 4n4yC 4n75A 4n7kH 4n7kL 4n7kM 4n7lH 4n7lL
4n7lM 4n7wA 4n7xA 4nefA 4nh2A 4njpA 4nppA 4npqA 4ntaA 4ntfA
4ntjA 4nv2A 4nv5A 4nv6A 4nykA 4o6mA 4o6nA 4o6yA 4o79A 4o7gA
4o93A 4o93B 4o9pB 4o9rA 4o9tA 4o9uA 4o9uE 4oaaA 4od4A 4od5A
4ogqA 4ogqB 4ogqC 4ogqE 4ogqF 4ogqG 4oh3A 4oj2X 4oo9A 4or2A
4ov0A 4oxsA 4oyeA 4oyfA 4p19A 4p24A 4p2zA 4p30A 4p3jA 4p6hA
4p6vA 4p6vB 4p6vC 4p6vD 4p6vE 4p6vF 4p79A 4p9oA 4p9pA 4pa4A
4pa6A 4pa7A 4pa9A 4pb2A 4pbua 4pbub 4pbuc 4pbud 4pbue 4pbuf
4pbuh 4pbui 4pbuj 4pbuk 4pbul 4pbut 4pbux 4pbuz 4pd4A 4pd4C
4pd4D 4pd4E 4pd4I 4pd5A 4pd6A 4pd7A 4pd8A 4pdaA 4pdlA 4pdmA
4pdrA 4pdvA 4pe5A 4pe5B 4pgrA 4pgsA 4pguA 4pgwA 4pi0B 4pi0C
4pi2A 4pi2C 4pirA 4pj0a 4pj0c 4pj0d 4pj0e 4pj0f 4pj0h 4pj0i
4pj0j 4pj0m 4pj0r 4pj0t 4pj0x 4pj0y 4pj0z 4pl0A 4popA 4povA
4pr7A 4pv1A 4pv1B 4pv1C 4pv1E 4pv1F 4pv1H 4pxfA 4pxkA 4pxzA
4py0A 4pypA 4q2gA 4q35A 4q35B 4q4aA 4q4aB 4q4hA 4q4hB 4q4jA
4q65A 4q79A 4q7cA 4q9hA 4q9iA 4q9jA 4q9lA 4qe7A 4qe9A 4qh1A
4qh4A 4qh5A 4qidA 4qimA 4qinA 4qiqA 4qkyA 4ql0A 4qnzA 4qo0A
4qo2A 4qryA 4qtnA 4quvA 4r0cA 4r50A 4r6zA 4r7cA 4r8cA 4r9uA

4r9uC 4raiA 4rcrH 4rcrL 4rcrM 4rdqA 4rdrA 4rdtA 4resA 4resB
4resE 4retA 4retB 4retE 4rfsT 4rhbA 4ri2A 4ri3A 4rku1 4rku2
4rku3 4rku4 4rkuD 4rkuE 4rkuH 4rkuI 4rkuJ 4rkuL 4rl8A 4rl9A
4rlcA 4ro2A 4rp9A 4rueA 4rufA 4rvwA 4rwaA 4rwdA 4rwsA 4ry2A
4ryjA 4rymA 4rynA 4ryqA 4ryrA 4s0fA 4s0vA 4tkqA 4tkrA 4tl3A
4tllA 4tllB 4tlmA 4tlmB 4tnha 4tnhb 4tnhc 4tnhd 4tnhf 4tnhg
4tnhh 4tnhl 4tnhm 4tnhx 4tnhz 4tnib 4tnic 4tnid 4tnie 4tnif
4tnig 4tnii 4tnij 4tnik 4tnit 4tnix 4tnja 4tnjb 4tnjc 4tnje
4tnjf 4tnjg 4tnjh 4tnji 4tnjj 4tnjk 4tnjl 4tnjm 4tnjt 4tnjx
4tnjz 4tnwA 4tphA 4tpjA 4tq3A 4tq4A 4tq5A 4tq6A 4tqqH 4tqqL
4tqqM 4tquN 4tquQ 4tquS 4tqvA 4tqvB 4tqvC 4tsyA 4twdA 4twfA
4twhA 4twkA 4u14A 4u15A 4u16A 4u1wA 4u1xA 4u1yA 4u2qA 4u3fA
4u3fC 4u3fD 4u3fE 4u3fF 4u4tA 4u4vA 4u4wA 4u9lA 4u9nA 4ub6a
4ub6b 4ub6c 4ub6e 4ub6h 4ub6i 4ub6j 4ub6k 4ub6l 4ub6m 4ub6x
4ub6z 4uc1A 4uc2A 4uc3A 4ug2A 4uhrA 4umvA 4up6A 4uqjA 4us4A
4uujC 4uv3A 4uvmA 4ux2B 4uxwA 4uxxA 4uxzA 4v1fA 4v1gA 4v3gA
4v3hA 4w6vA 4wabA 4wavA 4wd7A 4wd8A 4wfgA 4wfhA 4wgvA 4wgwA
4wibA 4wisA 4witA 4wmzA 4wo1A 4wolA 4ww3A 4x2sA 4x31A 4x5mA
4x5nA 4x5tA 4x88A 4x89A 4x8aA 4xdjA 4xdkA 4xdlA 4xe5A 4xe5B
4xe5G 4xeeA 4xesA 4xigM 4xigN 4xigS 4xk81 4xk82 4xk83 4xk84
4xk8d 4xk8e 4xk8f 4xk8g 4xk8h 4xk8i 4xk8j 4xk8k 4xk8l 4xniA
4xnjA 4xnkA 4xnlA 4xnvA 4xnwA 4xnxA 4xp1A 4xp4A 4xp5A 4xp6A
4xpaA 4xpbA 4xpfA 4xphA 4xptA 4xt1A 4xt1B 4xt3A 4xt3B 4xtcM
4xtcN 4xtcS 4xtlA 4xtnA 4xtoA 4xu5A 4xu6A 4xxjA 4xydA 4xydB
4y1kA 4y25A 4y3uA 4y3uB 4y7jA 4yayA 4yb9D 4ybqA 4yeuA 4yk5A
4yl0A 4yl1A 4yl3A 4ymkA 4ymsA 4ymtB 4ymuA 4ymuC 4ymvC 4ymwC
4ysxA 4ysxB 4ysxC 4ysxD 4ysyB 4ysyC 4ysyD 4yszB 4yszC 4yszD
4yt0C 4ytmB 4ytmD 4ytnB 4ytnC 4ytnD 4ytpB 4ytpC 4ytpD 4yxdB
4yxdC 4yzfA 4yziA 4z35A 4z36A 4z3pA 4z7fA 4z90A 4z91A 4z9gA
4zbmA 4zdyA 4ze3A 4zj8A 4zp0A 4zp2A 4zr0A 4zr1A 4zudA 4zw9A
4zwbA 4zwcA 4zyoA 4zyrA 4zzbA 4zzcA 5a1sA 5a2nA 5a2oA 5a40A
5a43A 5a44A 5a45A 5a63A 5a63B 5a63C 5a63D 5a6gB 5a6uB 5a6uG
5a8eA 5aexA 5af1A 5ah3A 5ahyA 5ahzA 5aidA 5ajiA 5an8A 5araA
5araG 5araI 5araJ 5araS 5araT 5araV 5araW 5ariJ 5ariT 5ariV
5ariW 5avqB 5avrB 5avrG 5avtB 5avtG 5avuG 5avvB 5avvG 5avwB
5avwG 5avxB 5avxG 5avyB 5avyG 5avzB 5avzG 5aw0B 5aw0G 5aw1B
5aw1G 5aw2B 5aw2G 5aw3B 5aw4B 5aw4G 5aw5B 5aw5G 5aw6B 5aw6G
5aw7B 5aw7G 5aw8G 5aw9B 5aw9G 5awwE 5awwY 5awzA 5ax0A 5ax1A
5ayoA 5aywA 5aywB 5aywC 5aywD 5azbA 5azcA 5azdA 5azoA 5azpA
5azsA 5b0wA 5b1aA 5b1aB 5b1aC 5b1aD 5b1aG 5b1aI 5b1aK 5b1aL
5b1aM 5b1bA 5b1bB 5b1bD 5b1bG 5b1bI 5b1bJ 5b1bK 5b1bL 5b1bM
5b34A 5b35A 5b3sA 5b3sB 5b3sC 5b3sG 5b3sI 5b3sJ 5b3sK 5b3sL
5b3sM 5b57A 5b57C 5b58A 5b58C 5b6vA 5b6xA 5b6yA 5b6zA 5bn2A
5bpsA 5bq6A 5bqaA 5bqhA 5bqiA 5br2A 5br5A 5butA 5butI 5bz2A

5c1mA 5c2tB 5c2tC 5c2tD 5c3jB 5c3jC 5c58A 5c58B 5c5xA 5c65A
5c6oA 5c78A 5cbfA 5cbgA 5cbhA 5cfbA 5cgdA 5ch4E 5ch4G 5ch4Y
5ckrA 5ctgA 5cthA 5cxvA 5d0oB 5d0oC 5d0oD 5d0qA 5d0qC 5d0qD
5d0qE 5d3mA 5d3mC 5d57A 5d58A 5d59A 5d6iA 5d6kA 5d91A 5da0A
5dgyA 5dhgA 5dhhA 5djqA 5djqC 5djqN 5dl5A 5dl8A 5dn6A 5dn6D
5dn6G 5dn6H 5dn6I 5dn6J 5dn6X 5dn6Z 5do7A 5do7B 5doqA 5doqB
5doqC 5dqqA 5dqxA 5dwkA 5dwyA 5dyeA 5dysA 5e1jA 5eabA 5eacA
5eadA 5eaeA 5eafA 5eagA 5eahA 5eblC 5ec1C 5ee7A 5eg1A 5eikA
5ej1A 5ej1B 5ekpA 5ekqD 5ekqE 5en0A 5eqbA 5eqgA 5eqhA 5eqiA
5eseA 5esfA 5eshA 5esiA 5eskA 5eslA 5esnA 5etzA 5eulA 5ezmA
5f15A 5f5bA 5f5dA 5f5gA 5f5jA 5f5kA 5f8uA 5fgnA 5fijJ 5fijT
5fijU 5fijW 5fl7D 5fl7G 5fl7H 5fl7K 5fn2A 5fn2C 5fn3B 5fn3C
5fn3D 5fn4A 5fn4B 5fn4C 5fn5A 5fn5B 5fn5C 5fn5D 5fokA 5fp2A
5fq6A 5fq6E 5fq7A 5fq7B 5fq7E 5fq7G 5fq8A 5fq8B 5fq8G 5fufA
5fvnA 5fxhA 5fxhB 5fxiB 5fxjA 5fxjB 5fxkA 5g28A 5g2aA 5g2dA
5g36A 5g53C 5gjvA 5gjvB 5gjvC 5gjvE 5gjvF 5gjwA 5gjwE 5gkoA
5glhA 5gmyA 5gpjA 5gthc 5gthd 5gthe 5gthh 5gthi 5gthj 5gthl
5gthm 5gtho 5gthR 5gtht 5gthx 5gthy 5gthz 5gtia 5gtib 5gtic
5gtid 5gtie 5gtih 5gtii 5gtij 5gtik 5gtil 5gtim 5gtiR 5gtit
5gtiy 5gtiz 5gxbA 5h1qA 5h2fa 5h2fb 5h2fc 5h2fd 5h2fe 5h2ff
5h2fh 5h2fi 5h2fk 5h2ft 5h2fx 5h2fy 5h2fz 5h2hA 5h2iA 5h2jA
5h2kA 5h2mA 5h2nA 5h2oA 5h2pA 5h35C 5h3oA 5hcjA 5hcmA 5hegA
5hehA 5hejA 5heuA 5hewA 5hi9A 5hk1A 5hk6A 5hk7A 5hv9A 5hvdA
5hvxA 5hwxA 5hxcA 5hxeA 5hxhA 5hxrA 5hxsA 5hyaA 5i1mV 5i32A
5i6cA 5i6zA 5i71A 5i73A 5i74A 5i75A 5i9kA 5ia9A 5id3A 5ideA
5ideB 5iofA 5iouB 5ir6C 5irxA 5irzB 5is0B 5itcA 5iteA 5iu4A
5iu7A 5iu8A 5iuaA 5iubA 5iuxA 5iuyA 5iwkA 5iwnA 5iwoA 5iwpA
5iwrA 5iwsA 5iwtA 5iy5A 5iy5C 5iy5D 5iy5G 5iy5I 5iy5J 5iy5K
5iy5L 5iy5M 5iyuA 5j0zA 5j4iA 5j4nA 5jagA 5jdfA 5jdgA 5jdhA
5jdlA 5jdnA 5jdqA 5jefA 5jeqA 5jgpA 5jjeA 5jkiA 5jnqA 5jqhA
5jr0A 5jrfA 5jrwA 5jsiA 5jtbA 5jwyA 5jynA 5k0iA 5k47A 5k7lA
5k7lB 5kbnA 5kbsA 5kbtA 5kbuA 5kbvA 5kbwA 5kc0A 5kc4A 5khnA
5khsA 5kk2A 5kk2E 5kkzA 5kkzB 5klbA 5klgA 5kliA 5kliB 5kliC
5klsA 5kmfA 5kmhA 5ko2A 5komA 5koyA 5kpdA 5kpiA 5kpjA 5kufA
5kukA 5kumA 5kxiA 5kxiB 5l1bA 5l1eA 5l1gA 5l1hA 5l24A 5l25A
5l2aA 5l2bA 5l47A 5l4eA 5l4hA 5l75F 5l7dA 5l7iA 5l8r1 5l8r2
5l8r3 5l8r4 5l8rD 5l8rE 5l8rF 5l8rG 5l8rH 5l8rI 5l8rJ 5l8rK
5lbjA 5ldvA 5levA 5lidA 5lilA 5lj6A 5ljoB 5ljoC 5ljoD 5ljoE
5llmA 5lm4A 5lqxA 5lqxG 5lqxH 5lqxI 5lqxK 5lqxU 5lqxV 5lqxW
5lqyA 5lqyK 5lqyV 5lqyY 5lriH 5lriL 5lriM 5lseH 5lseL 5lseM
5lv6A 5lweA 5lwyA 5lxaA 5lxgA 5lzrA 5m7jA 5m7jB 5m7jD 5m7kA
5m7kC 5m7kD 5m7lA 5m7lB 5m7lC 5m7lD 5m87A 5m8jA 5m94A 5mdqA
5mdrA 5mg3C 5mg3E 5mg3F 5mg3G 5mg3Y 5mjuA 5mkeA 5mkkA 5mkkB
5mm0A 5mm1A 5mmtA 5mrwA 5mrwB 5mrwC 5mrwD 5mt6A 5mt7A 5muoA

5murA 5mvmA 5mvnA 5mwvA 5mx2a 5mx2b 5mx2c 5mx2d 5mx2e 5mx2h
5mx2i 5mx2k 5mx2l 5mx2m 5mx2o 5mx2t 5mx2x 5mx2z 5mzpA 5mzqA
5mzrA 5mztA 5n2rA 5n6hA 5n6lA 5n6mA 5n9yA 5nbdA 5nc4A 5nc5F
5ndcA 5ndcB 5ndcC 5nddA 5ndzA 5nj3A 5njyA 5nlxA 5nm4A 5nmiA
5nmiC 5nmiE 5nmiF 5nmiG 5nmiH 5nmiJ 5nr2A 5nupA 5nupD 5nuqA
5nuqG 5nurA 5nurB 5nv9A 5nvaA 5nxnA 5nxuA 5o4cC 5o4cH 5o4cL
5o5eA 5o64C 5o64L 5o64M 5o65A 5o67A 5o67B 5o68A 5o77A 5o78A
5o79A 5o8fA 5o8oA 5o9cA 5ob4A 5oc0A 5oc9A 5ochA 5odwC 5ofpA
5ogeA 5ogkA 5ojmA 5okdA 5okdB 5okdC 5okdD 5okdG 5okdI 5okdJ
5oloA 5olvA 5olzA 5om1A 5om4A 5onuA 5oonA 5oqtA 5oxlA 5oxmA
5oxnA 5oxoA 5oxpA 5oybA 5oygA 5oykA 5prcC 5prcL 5prnA 5sv9A
5svjA 5svkA 5svlA 5svmA 5svpA 5svqA 5svrA 5svsA 5sxuA 5sxvA
5sy1A 5sy1C 5sytA 5t04A 5t1aA 5t36A 5t37A 5t3rA 5t4dA 5t5nA
5tcxA 5te3A 5te5A 5tgzA 5tinA 5tioA 5tj6A 5tjiA 5tl9A 5tqqA
5tr1A 5tsaA 5tsbA 5ttpA 5tuaA 5tv4A 5tvnA 5tzrA 5u1dA 5u1dB
5u1dX 5u1lA 5u1vA 5u1wA 5u1xA 5u1yA 5u6oA 5u6pA 5u70A 5u73A
5u74A 5u9wA 5uarA 5uenA 5uhsA 5uigA 5uiwA 5uiwB 5uj9A 5ujaA
5uk6A 5ul0A 5ul7A 5ul9A 5uldA 5uleA 5un1A 5un1B 5unfA 5ungB
5unhA 5uniA 5uniB 5uowA 5uowB 5uowD 5up2A 5up2B 5up2D 5uz7A
5uz7B 5uz7G 5uz7R 5v2sA 5v33H 5v33M 5v4sA 5v57A 5v6nA 5v6pA
5v7pD 5v8kA 5va1A 5va2A 5va3A 5vaiG 5vaiP 5vaiR 5vb2A 5vdhA
5vdiA 5vewA 5vexA 5vg9A 5vhwA 5vhxA 5vhyA 5vhzA 5vmsB 5vn9A
5votA 5votE 5vouA 5vouE 5vovA 5vovE 5vpnA 5vpnB 5vpnC 5vpnD
5vraA 5vrgA 5w3sA 5w81A 5wehA 5wehB 5wehC 5wehD 5welA 5wemA
5wenA 5weoA 5wiuA 5wivA 5wj5A 5wj9A 5wktA 5wkuA 5wkvA 5wkyA
5woaA 5wpqA 5wptA 5wpvA 5wqcA 5ws3A 5ws5a 5ws5b 5ws5c 5ws5d
5ws5e 5ws5f 5ws5h 5ws5i 5ws5j 5ws5k 5ws5m 5ws5R 5ws5t 5ws5x
5ws5y 5ws5z 5ws6a 5ws6b 5ws6c 5ws6d 5ws6e 5ws6f 5ws6h 5ws6j
5ws6k 5ws6l 5ws6t 5ws6x 5ws6y 5ws6z 5wtrA 5wucA 5wudA 5wueA
5wufA 5wzyA 5x0mA 5x19A 5x19B 5x19D 5x19G 5x19I 5x19J 5x19K
5x19M 5x1bA 5x1bB 5x1bC 5x1bD 5x1bJ 5x1bK 5x1bL 5x1bM 5x1fA
5x1fB 5x1fC 5x1fD 5x1fG 5x1fI 5x1fJ 5x1fK 5x1fL 5x1fM 5x33A
5x3xa 5x41A 5x41E 5x41F 5x5yA 5x5yF 5x7dA 5x87A 5x9hA 5x9rA
5xamA 5xarA 5xatA 5xdnA 5xdoA 5xdqA 5xdqC 5xdqD 5xdqG 5xdqI
5xdqJ 5xdqK 5xdqM 5xdxA 5xdxB 5xdxC 5xdxD 5xdxG 5xdxJ 5xdxK
5xdxL 5xdxM 5xezA 5xf1A 5xhqA 5xj6A 5xj7A 5xj8A 5xjjA 5xjmA
5xjyA 5xlsA 5xnn1 5xnn3 5xnn4 5xno1 5xno3 5xno4 5xpdA 5xprA
5xr8A 5xraA 5xsyA 5xszA 5xteB 5xteC 5xteE 5xteF 5xteG 5xteL
5xu1A 5xu1M 5xw6A 5y50A 5y78A 5y79A 5y83A 5yc8A 5yckA 5ydzA
5ye1A 5ye5A 5yluA 5ylvA 5ylvB 5yqzP 5yqzR 5ys3A 5ys8A 5yveA
5yw7B 5ywdB 5z1lA 5z1wA 5z62A 5z62B 5z62C 5z62D 5z62E 5z62F
5z62G 5z62H 5z62I 5z62K 5z62L 5z62M 5z62N 5z84A 5z84C 5z84D
5z84G 5z84J 5z84K 5z84L 5z84M 5z85A 5z85D 5z85I 5z85J 5z85K
5z85L 5z86C 5z86D 5z86G 5z86I 5z86J 5z86K 5z86L 5z86M 5z96A

5zazA 5zbgA 5zbhA 5zbqA 5zcoB 5zcoD 5zcoH 5zcoI 5zcoK 5zcoL
5zcoM 5zcpA 5zcpB 5zcpI 5zcpJ 5zcpK 5zcpM 5zfpA 5zfuA 5zgb1
5zgb2 5zgb3 5zgbC 5zgbD 5zgbI 5zgbJ 5zgbK 5zgbL 5zgbO 5zgh1
5zgh2 5zghF 5zghI 5zghJ 5zghK 5zghM 5zghO 5zilA 5zimA 5zinA
5zk3A 5zk8A 5zkbA 5zkpA 5zleA 5ztyA 5zugA 6a2jA 6a6nA 6a70A
6a70B 6a90A 6a91B 6a93A 6a94A 6a96A 6a96B 6agfA 6ajgA 6akfA
6akfB 6akxA 6al2A 6aqfA 6aveA 6awfB 6awfC 6awfD 6awoA 6awqA
6b21A 6b24A 6b24C 6b2bA 6b2dA 6b3jA 6b3jB 6b3jG 6b3jP 6b3jR
6b5vA 6b73A 6b85A 6b87A 6barA 6basA 6batA 6bavA 6bbfA 6bbgA
6bbhA 6bbiA 6bbjA 6bd4A 6be1A 6bgiA 6bgjA 6bh8A 6bhpA 6bhuA
6bl6A 6bmiA 6bmsA 6bo4A 6bo5A 6bo8A 6bo9A 6boaA 6bobA 6bplA
6bpnA 6bppA 6bqgA 6bqoA 6bqrA 6btmA 6btmC 6btmD 6btmF 6bu5A
6bugA 6bugC 6buhA 6buhC 6buiA 6bvgA 6bw6A 6bwfA 6bwmA 6bx4A
6by2C 6by3C 6byoA 6c08C 6c0vA 6c14B 6c1eA 6c1kA 6c1qB 6c261
6c262 6c263 6c264 6c265 6c26A 6c26C 6c3iA 6c6lA 6c6lB 6c6lC
6c6lD 6c6lE 6c6lM 6c6lN 6c6lO 6c70A 6c8fA 6c8gA 6c8hA 6c9aA
6caaA 6cb2A 6cfwA 6cfwB 6cfwC 6cfwD 6cfwE 6cfwF 6cfwG 6cfwH
6cfwI 6cfwJ 6cfwK 6cfwM 6cfwN 6ci0A 6ci0B 6cjqA 6cjtA 6cm4A
6cmoA 6cmoB 6cmoR 6cnaA 6cnaB 6cnjA 6cnjB 6cnkA 6cnkC 6cnmE
6cnnA 6cnnE 6cnoA 6coyA 6cp77 6cp78 6cp7J 6cp7K 6cp7X 6cp7Z
6cpvA 6cseC 6csfC 6csmA 6csxA 6ctdA 6cudA 6cvlA 6cvlC 6cvlE
6cx0A 6cxhA 6cxhB 6cxhC 6d0jA 6d0jC 6d0kA 6d0kC 6d0nA 6d0nC
6d1sA 6d1wA 6d26A 6d27A 6d32A 6d35A 6d3rA 6d3sA 6d6uB 6d6uE
6d7oA 6d7pA 6d7qA 6d7sA 6d7tA 6d7vA 6d7wA 6d7xA 6d91A 6d9wA
6d9zA 6ddeA 6ddeB 6ddeC 6ddeR 6ddfR 6dg7A 6dg8A 6djbA 6djzA
6dk0A 6dk1A 6dm0A 6dm1A 6dmoA 6dmrA 6dmuA 6dmwA 6dmwE 6dmyA
6dmyB 6dnfA 6do1A 6drjA 6drkA 6drxA 6dryA 6ds0A 6dt0A 6du8A
6dvyA 6dvzA 6dw0A 6dw0B 6dw0D 6dz1A 6dz7A 6e0hA 6e14C 6e14D
6e14G 6e15C 6e15D 6e15G 6e15H 6e1hA 6e1hC 6e1kA 6e1kC 6e1nA
6e1pA 6e2fE 6e2gA 6e3yA 6e3yB 6e3yE 6e3yG 6e3yR 6e59A 6e7pA
6e7yA 6e7zA 6e8qA 6e8wA 6ebkA 6ebkB 6ebmB 6edqA 6ehcA 6ehdA
6eheA 6ehfA 6eiaA 6eidA 6eigA 6ej6A 6ejhA 6emxA 6eneA 6etiA
6eusA 6eyuA 6eznB 6eznC 6eznD 6eznE 6eznF 6eznH 6f0iA 6f0jA
6f0kA 6f0kC 6f0kD 6f0kF 6f0kH 6f0mA 6f0nA 6f0rA 6f0uA 6f0vA
6f0zA 6f10A 6f11A 6f13A 6f15A 6f16A 6f34A 6f36A 6f36M 6f36N
6f46A 6f7aA 6f7hA 6feqA 6ffcA 6ffvA 6fk6A 6fk8A 6fk9A 6fkaA
6fkdA 6fkfa 6fkfA 6fkfb 6fkfe 6fkfg 6fkfG 6fkfp 6fkha 6fkhg
6fkhG 6fkhp 6fkiA 6fkib 6fkiG 6fkip 6fl9A 6fliA 6fmrA 6fmsA
6fmtA 6fmwA 6fmyA 6fn1A 6fn4A 6fo6A 6fo6B 6fo6C 6fo6D 6fo6F
6fo6G 6fo6J 6fos2 6fosD 6fosE 6fosF 6fosK 6fosM 6fosO 6fsuA
6fufA 6fufB 6fvqA 6fvrA 6fvsA 6fwfA 6fwzA 6g1kA 6g79A 6g79B
6g79G 6g7hA 6g7iA 6g7jA 6g7kA 6g7lA 6g7oA 6g8zA 6g9lA 6g9oA
6gciA 6gctA 6gdgA 6gdgC 6gdgD 6gdiA 6ghjA 6gieA 6gs1A 6gs4A
6gs7A 6gv1A 6gxcA 6gz9A 6h3iB 6h3iF 6h7dA 6h7fA 6h7jA 6h7jE

6h7lA 6h7lE 6h7mA 6h7mE 6h7nA 6h7nE 6h7oA 6h7oE 6h7uA 6hawA
6hawB 6hawC 6hawD 6hawE 6hawG 6hawH 6hawJ 6hbuA 6hcoA 6hcpA
6hd1A 6hijA 6hinA 6hioA 6hiqA 6hisA 6hllA 6hloA 6hlpA 6hqbD
6hqbE 6hqbF 6hqbI 6hqbJ 6hqbL 6hqbM 6hraA 6hraB 6hraC 6hraD
6hrbC 6hrbD 6hugA 6hugB 6hugC 6hujA 6hujB 6hujC 6hukA 6hukB
6hukC 6humA 6humB 6humC 6humD 6humE 6humG 6humH 6humJ 6humK
6humL 6humN 6humO 6humP 6humQ 6huoA 6huoB 6huoC 6hupA 6hupB
6hy9A 6hyaA 6hyrA 6hyvA 6hywA 6hyxA 6hyzA 6hz0A 6hz3A 6hzpA
6hzwA 6i08A 6i2jA 6i53A 6i53B 6i53C 6i6bA 6i6hA 6i6jA 6iblA
6ic4A 6ic4G 6ic4I 6ic4K 6idfA 6idfB 6idfC 6idfD 6idfE 6idpA
6idsA 6iedA 6iglA 6igz1 6igz2 6igz3 6igz7 6igz9 6igzD 6igzE
6igzG 6igzH 6igzI 6igzJ 6igzK 6igzM 6iiuA 6iivA 6ijzA 6iraA
6irfA 6iu3A 6iu4A 6iycB 6iycC 6iycD 6metB 6mg8A 6mgvA 6mgwA
6mhoA 6mhqA 6mhwA 6mhxA 6mm9A 6mm9B 6mmaA 6mmgA 6mmgB 6mmjA
6mmkA 6mmnB 6mmsB 6mmtA 6msmA 6mvvA 6mvwA 6mwbB 6mwdB 6mwgB
6mx4A 6mx4B 6mx4C 6n23A 6n24A 6n25A 6n26A 6n27A 6n28A 6n3qA
6n3qB 6n3qC 6n3qE 6n3qF 6n3tA 6n40A 6n4bA 6n4bC 6n4bR 6n4iE
6n4qA 6n4qE 6n4rA 6n4rE 6n51A 6n52A 6nbqA 6nbqB 6nbqC 6nbqD
6nbqE 6nbqH 6nbqI 6nbqJ 6nbqK 6nbqL 6nbqM 6nbqN 6nbqO 6nbqP
6nbxA 6nbxC 6nbxD 6nbxE 6nbxF 6nbxG 6nbxH 6nbxI 6nbxK 6nbxL
6nbxN 6nbxQ 6nd1A 6nd1B 6nd1C 6nd1E 6nd1F 6nhwA 6nhyA 6niyA
6niyR 6nt3A 6nt4A 6nt4B 6nziA 6nzwA 6nzzA 6prcC 6prcH 6prcL
6prcM 6prnA 6q5eA 6qeeA 6qexA 7prcC 7prcL 7prcM 7prnA 8prnA

#### 1.2 The test set (500 sequences)

1a91A 1bccA 1be3B 1be3D 1bl8A 1dv3H 1e14H 1e14L 1fnqM 1gueA
1h2sA 1h2sB 1hzxA 1ijdB 1ijpA 1ixfA 1izlF 1j4nA 1jb0E 1jfpA
1jgzL 1kb9C 1kyoE 1kyoG 1kyoH 1l0lH 1l0nF 1l9jH 1m0lA 1mm4A
1nenC 1ntmG 1ocrC 1oczA 1oczK 1oedB 1oedE 1oyeA 1p84I 1p8hA
1pjfA 1po3A 1pssL 1pstH 1r3jC 1rqkH 1rzhL 1s5hC 1s5lc 1s5lf
1s5lt 1s8jA 1sqbC 1sqbE 1t9tA 1v54J 1v55L 1xmeA 1xqeA 1yewB
1z98A 1z9kA 1zoyD 2a06J 2ahyA 2ahzA 2akhB 2axtz 2bl2A 2bnpC
2bocC 2bozL 2brdA 2d2cC 2d2cD 2d2cH 2dysB 2dysJ 2eikC 2eilK
2eimB 2f93A 2fbwD 2fkwA 2g87A 2gmrL 2hacA 2hj6H 2hn2A 2htlA
2i36A 2iahA 2ibzD 2itcC 2j5dA 2l16A 2l9uA 2lmeA 2lzlA 2muwA
2mv6A 2n28A 2n90A 2n9yB 2nwwA 2o01C 2oauA 2occK 2pilA 2q68A
2qpeB 2r8aA 2ux3L 2ux4H 2uxjM 2vddA 2wdrD 2wjmM 2wjnH 2wpdJ
2wseI 2wsfL 2ww9M 2wwbA 2x56A 2x5vL 2y00A 2y69E 2yksA 2zjsE
2zjsY 2zt9C 2zt9D 2zz9A 3a0bj 3abkC 3abkD 3ablG 3ablJ 3abmC
3abmM 3ae1B 3ae1C 3ae2C 3ae4D 3ae9C 3ae9D 3aecD 3asnC 3asnM
3aygA 3b5dA 3b9zA 3cwbD 3cwbI 3cxhA 3dknA 3dknC 3dsyH 3du3H
3du3L 3duqH 3dwoX 3e9jC 3eamA 3f48A 3g61A 3h1hA 3h1hC 3h1lC
3hasA 3hkkA 3ir5C 3ir7C 3ixzB 3j5qB 3jd8A 3k1bA 3kssA 3l74E
3l75G 3ldeA 3lnmB 3lw5N 3m75A 3m78A 3mk7B 3nkaA 3nkcA 3ny8A
3o0eA 3oaxA 3ob6A 3odjA 3om3A 3omnB 3p4sC 3pcqF 3pcvA 3pgsA
3pjzA 3porA 3puxF 3pxoA 3qakA 3qbiA 3qjqC 3qjvB 3qs5A 3rgbC
3rkoA 3rkoB 3rlfA 3rw0A 3s38A 3s38C 3s3aB 3sn6R 3syaA 3szvA
3tdoA 3tdxA 3te1A 3tijA 3tlwA 3txtA 3uq5A 3uq7A 3uu5A 3vgaA
3vr9C 3vrbD 3vy9X 3wmmH 3wu2i 3x2qB 3zebA 4ainA 4al1A 4amjA
4aq5C 4auiA 4b4aA 4booA 4booB 4botA 4botB 4c9gA 4chwA 4cofA
4d2eA 4d5bA 4d6uJ 4e1sA 4e1tA 4faaA 4fe1F 4fi3C 4frxA 4fz0A
4fz1A 4g72B 4g7sC 4gbrA 4gcqA 4gcsA 4h13G 4h99L 4hea1 4hmkA
4ikzA 4il6h 4il6i 4in7H 4iu8A 4ixqa 4ixrh 4izmA 4j05A 4jtaB
4kt0C 4kt0J 4l35A 4lsfA 4m2sA 4m48A 4m64A 4mrvA 4njnA 4ntbA
4o9pA 4ogqD 4ogqH 4p1aA 4pa3A 4pd4G 4pj0b 4pj0k 4pj0l 4pj0u
4px7A 4q2eA 4q4jB 4r1iA 4rfsS 4rhbB 4rkuG 4rlbA 4rngA 4ryiA
4ryoA 4tnhi 4tnht 4tnih 4tnil 4u3fG 4u3fH 4u4fA 4ub6f 4ub6t
4umwA 4us3A 4x32A 4xp9C 4xu4A 4y9hA 4ytmC 4yxdD 4z34A 4zjcA
4zowA 5a6uA 5aezA 5araH 5araU 5avsG 5avuB 5aw8B 5awwG 5aymA
5b1aJ 5b2nA 5b6wA 5bqjA 5c6nA 5c76A 5d0oE 5d5aA 5d6lA 5d92A
5dirA 5djqB 5dl6A 5edlA 5eh6A 5ekeA 5ekqA 5esmA 5eulY 5g26A
5g2cA 5g53A 5gthb 5gthf 5gthk 5gtix 5h36A 5hd8A 5heoA 5ir6A
5ir6B 5iy5B 5j7aA 5jjeB 5jmnD 5l27A 5l75A 5l75G 5lg3A 5lqxD
5m7jC 5m8aA 5m95A 5mdoA 5mdpA 5mdsA 5mkfA 5mlzA 5mx2f 5naoA
5nj6A 5nm2A 5nx2A 5o0tA 5o4uA 5okdE 5okdF 5or1A 5osbA 5oxkA
5sv1A 5svtA 5t77A 5tzyA 5u09A 5uhqA 5uviA 5v56A 5vblB 5vrhA
5wekA 5wf6A 5wkxA 5ws5l 5x19L 5x1bG 5x3xm 5x5yG 5xdxI 5xj9A
5xteD 5xteH 5xteJ 5ye2A 5yluB 5z62J 5z84I 5z85C 5z85G 5z85M

5zcoG 5zcpC 5zcpD 5zcpL 5zgbM 5zgh3 5zghL 5zlgA 5ztlA 6a69A
6a91A 6ak3A 6akyA 6awfA 6awpA 6ayfA 6aygA 6bauA 6bcjA 6bwiA
6bx5A 6c1rB 6c96A 6cjuA 6co7A 6cp7U 6drzA 6dvwA 6e14F 6e2fA
6eu6A 6euqA 6eznA 6f0kE 6f12A 6f2dA 6f2dF 6f2dG 6fkfB 6fosJ
6g79S 6gpsA 6gpxA 6hqbK 6hrbB 6humI 6humM 6humS 6hupC 6idrA
6igz6 6igzC 6igzF 6igzL 6iraB 6iycE 6meoB 6meoG 6mhyA 6mmtD
6mmwB 6mmxB 6n4iA 6nbqF 6nbqG 6nbqS 6nbxB 6nd1D 6o00A 6q81A

#### 1.3 The validation set (400 sequences)

1by3A 1dv3M 1dxrH 1eysH 1f6nL 1jgxL 1kg8A 1kyoW 1kzuA 1kzuB
1l0lG 1l0lJ 1l7vA 1ldiA 1mpoA 1mprA 1nenB 1ntkE 1ntkF 1ntkK
1ntmA 1occJ 1ocoC 1ocoJ 1ocrJ 1ogvM 1p8uA 1prcL 1pssH 1q90M
1qcrA 1qcrG 1qovH 1qovM 1rg5H 1rklA 1rvjM 1ry5L 1rzzL 1rzzM
1s54A 1siwB 1t16A 1umxM 1vrnC 1xkwA 1xqfA 1zp0B 2abmA 2akhA
2akiC 2axta 2b2hA 2b76B 2bnsB 2c3eA 2cpbA 2cpsA 2e74D 2e74E
2e74H 2eijG 2eijK 2eilA 2f93B 2fgrA 2fyuC 2hg9H 2hlfA 2hqdA
2hvjC 2j8cL 2jj0L 2jo1A 2kv5A 2lz3A 2mfrA 2mg2A 2mprA 2muvA
2n2lA 2nwlA 2o4vA 2o9fA 2occD 2occJ 2occL 2pedA 2q6aA 2r4nA
2r6gE 2r88A 2rcrL 2rddB 2rh1A 2ux5H 2wjnL 2wliA 2wpdG 2ws3D
2wscF 2wsfG 2wsfJ 2wwbN 2x5vH 2xutA 2y69F 2z73A 2zqpY 2zxwL
2zy9A 3a0bi 3a0by 3abkA 3ae2B 3ae3B 3ae8C 3aeeC 3aegC 3ag1K
3ag2G 3ag3B 3ag3C 3ag3I 3aodA 3b60A 3bccC 3bo1B 3bo1C 3c1hA
3c1jA 3c9lA 3chxC 3cirD 3cwbG 3cx5A 3cx5E 3cxhI 3detA 3dinE
3e8bA 3effK 3ei0A 3fyeB 3h1lA 3hapA 3hd7B 3igaC 3j45y 3k0gA
3l72E 3ldcA 3m4dA 3n23E 3o0rB 3o7qA 3or7C 3poqA 3puwA 3qjvA
3rhwA 3s0xA 3s33C 3s3cA 3s8gB 3t45A 3uu3A 3v8gA 3vvsA 3wfdB
3wfeB 3wkvA 3wu2t 3zmhA 3zprA 3zuwL 4afkA 4au5A 4c9hA 4cg7A
4cg7C 4cu4A 4cz8A 4d5uA 4d6uD 4eneA 4fe1A 4fi3F 4fz0M 4g1uC
4g72A 4g7rA 4g7rB 4gx2A 4h13B 4h13H 4h44H 4h9lL 4hbhM 4hea3
4hukA 4hzuT 4ilaA 4j7tA 4jczA 4k0eA 4kfmG 4khzF 4ki0F 4kkcA
4kklA 4kt0E 4kx6C 4ky0A 4m8jA 4mmaA 4mrrA 4mrsA 4mtfA 4mycA
4n4dA 4n4rB 4n4yA 4o9tB 4o9uB 4p00B 4pb1A 4pbuy 4pd4H 4pgvA
4phuA 4pi0A 4pi2B 4pv1G 4q9kA 4qi1A 4qncA 4qndA 4rp8A 4tnhe
4tnhj 4tnhk 4tniz 4tnjd 4tnvA 4tpgA 4u3fJ 4ub6d 4ub6R 4ub6y
4ux1B 4xpgA 4y7kA 4ycrA 4ymsC 4ymtA 4ymwA 4yt0D 4ze1A 4zwjA
5a3rA 5aw3G 5aynA 5aywE 5b1bC 5b3sD 5b3sF 5bqgA 5bz3A 5c3jD
5c6pA 5c73A 5cfyA 5d3mB 5d3mD 5d5bA 5dl7A 5ebwC 5esgA 5esjA
5eulE 5fjgA 5fl7A 5fn2B 5fn2D 5fn4D 5fxkB 5gtha 5gtif 5gufA
5h2fj 5h2fl 5hs1A 5i20A 5ijiA 5irxE 5iy51 5iy5H 5kkhA 5kkzC
5kteA 5l22A 5l26A 5l8rL 5ldtA 5lqxJ 5lqxY 5lx9A 5m7kB 5m8kA
5mg3D 5nkqA 5nmiD 5nxrA 5o4cM 5o64H 5oekA 5olgA 5osaA 5prcH
5prcM 5sv0A 5uakA 5v33L 5v7pA 5vkvA 5vmsA 5wieB 5ws4A 5ws6i
5ws6m 5ws6R 5x19C 5x1bI 5x3xq 5xasA 5xdqB 5xdqL 5xteA 5xteK
5z1fA 5z84B 5z85B 5z86A 5zcoA 5zcoC 5zcoJ 5zcpG 5zfvA 5zgbE
5zgbF 5zkcA 5zsuA 6a90B 6awnA 6ayeA 6b3uA 6bqhA 6btmE 6bwdA
6bwjA 6c14A 6cfwL 6cnmA 6ctfA 6d6uA 6d80A 6dlzA 6e14H 6e1mA
6e1oA 6ehbA 6exsA 6f34C 6fk7A 6fkfd 6fkhb 6fkia 6fm9A 6fmvA
6fmxA 6fo6E 6fos4 6fosL 6hawF 6hrbA 6humF 6hy5A 6igkA 6igz0
6igz4 6irgA 6irhA 6iycA 6mhsA 6mwaB 6n3qD 6nbxP 7ahlA 7prcH

#### 2. The small dataset for the MCP (Membrane Contact Probability) model

##### 2.1 The training set (718 sequences)

1a0sP 1a91A 1ar1B 1ay2A 1b9uA 1dxrH 1e54A 1ehkB 1ezvB 1fftA
1fftB 1fftC 1fjkA 1fx8A 1h6s1 1iijA 1ijdA 1ijdB 1izl0 1izlA
1izlC 1izlD 1jb0M 1jb0X 1k24A 1kb9A 1kf6A 1kf6B 1kf6C 1kf6D
1kpkA 1kqfC 1kyoW 1l0lK 1l7vC 1l9bC 1lghA 1m56A 1m56D 1mm4A
1mprA 1nekD 1nqeA 1occB 1occH 1p49A 1p7bA 1pjfA 1pw4A 1q16B
1q90B 1q90G 1q90L 1q90M 1q90N 1q90R 1qleC 1qo1J 1rh5C 1rklA
1s5lh 1s5li 1s5lj 1s5ll 1s5lm 1s5lt 1s5lu 1s5lx 1sqbA 1sqbB
1t16A 1uynX 1vf5B 1vf5D 1vf5E 1vf5F 1vf5H 1wrgA 1xioA 1xkwA
1xl4A 1yc9A 1yewC 1zzaA 2a65A 2a9hE 2bl2A 2c0xA 2ervA 2evuA
2f1cX 2f93A 2f95B 2fynC 2grxC 2hacA 2hdiB 2hi7A 2hydA 2iahA
2ibzF 2ibzI 2j5dA 2j8sD 2jlnA 2jp3A 2jwaA 2k0lA 2k73A 2k9pA
2k9yA 2kixA 2kluA 2kncA 2ks9A 2ksfA 2kv5A 2kyhA 2l0jA 2l2tA
2l35A 2l5bA 2l8sA 2l9uA 2lckA 2lhfA 2lifA 2lmeA 2lnlA 2lorA
2lzlA 2m20A 2m59A 2m67A 2mc7A 2mjoA 2mmuA 2mofA 2mpnA 2mxbA
2n1pA 2n4xA 2na8A 2ndjA 2nq2A 2nq2C 2nr9A 2nrgA 2nuuG 2o4vA
2oauA 2porA 2qfiA 2r6gA 2r6gF 2rddB 2voyA 2vpwB 2vpwC 2vqiA
2w1pA 2witA 2wjqA 2wpdI 2wscC 2wscE 2wscG 2wscH 2wscI 2wscJ
2wscN 2wswA 2ww9A 2ww9B 2ww9H 2ww9I 2ww9J 2ww9L 2ww9N 2ww9O
2x27X 2x2vA 2xokD 2xokG 2xq2A 2xzbB 2y5yA 2y69D 2y69G 2y69I
2y69J 2y69K 2y69L 2y69M 2yevB 2yevC 2ynkA 2yvxA 2z73A 2zjsE
2zt9E 2zxeB 3a0bt 3a2sX 3a7kA 3b4rA 3b5dA 3b9wA 3bryA 3chxB
3cn5A 3cslC 3d31A 3d31C 3d5kA 3ddlA 3dinE 3dzmA 3eh3A 3emnX
3emoA 3fhhA 3gi8C 3hd6A 3hd7A 3hd7B 3hd7C 3hd7D 3hw9A 3iyzA
3j41E 3j45T 3j45Y 3jtyA 3jycA 3k3fA 3kj6A 3kp9A 3kvnA 3llqA
3m71A 3mk7A 3mk7B 3mk7C 3mp7A 3ne5B 3njtA 3o0rB 3o7pA 3orgA
3pjsK 3pjzA 3q7kA 3qf4A 3qf4B 3qnqA 3qq2A 3qraA 3rkoA 3rkoB
3rkoC 3rkoD 3rkoF 3rkoG 3rlbA 3sybA 3tdoA 3tuiC 3ug9A 3ukmA
3um7A 3ux4A 3v5sA 3v8xB 3vmqA 3vr8B 3w5aC 3wdoA 3wkvA 3wmfA
3wmm1 3wmmC 3wmmM 3wo7A 3x29B 3x2rA 3x3bA 3zd0A 3zevA 3zuxA
4a2nB 4aq5B 4aq5C 4atvA 4aywA 4b4aA 4bgnA 4bpmA 4c00A 4c48C
4c9jA 4cadC 4chvA 4cy4A 4cz8A 4czbA 4d5bA 4d6tE 4d6tH 4d6tJ
4dblE 4djiA 4dxwA 4ezcA 4fc4A 4fe1X 4fqeA 4frxA 4fsoA 4fz0M
4g7vS 4gbyA 4gd3A 4gd3J 4gd3Q 4geyA 4gx0A 4h56A 4he8A 4he8D
4he8G 4he8I 4hea1 4hea2 4hea4 4hea5 4hea7 4hea9 4hukA 4hukB
4huqA 4huqS 4huqT 4hw9A 4hycA 4hyjA 4hyoA 4ikvA 4il3A 4il6t
4in5L 4ixqk 4ixqo 4j05A 4jbwM 4k1cA 4knfA 4kt0F 4kt0M 4ldsA
4m58A 4m64A 4mndA 4mqsA 4mrnA 4mt4A 4n4rB 4njnA 4o6mA 4o6yA
4o9pA 4o9uB 4od4A 4ogqC 4oh3A 4p6vA 4p6vB 4p6vC 4p6vD 4p6vE
4p6vF 4pbut 4pgrA 4pi0A 4pj0r 4pl0A 4px7A 4q2eA 4qndA 4qtnA
4quvA 4rdqA 4rfsS 4rl8A 4rl9A 4rlcA 4rp8A 4rwsC 4ryiA 4tkrA
4tq3A 4tquM 4tquN 4tquQ 4tsyA 4u4tA 4uc1A 4us3A 4v1fA 4w6vA

4wd7A 4wmzA 4x5mA 4xnkA 4xnvA 4xt1A 4xt1B 4xu4A 4xxjA 4xydB
4y7jA 4yayA 4ymkA 4ymsA 4ymsC 4ytpC 4z3pA 4z7fA 4zr0A 4zw9A
5a1sA 5a40A 5a63B 5a63C 5a63D 5araG 5araH 5araI 5araS 5araT
5araU 5araV 5araW 5awwG 5awwY 5awzA 5azbA 5b2nA 5b57A 5b57C
5c73A 5cbfA 5cfbA 5ctgA 5d0oB 5d0oC 5d0oE 5d3mA 5d3mC 5dirA
5djqN 5dl5A 5dl7A 5dn6G 5dn6H 5dn6I 5dn6J 5dn6X 5dn6Z 5do7B
5doqA 5doqB 5doqC 5dqxA 5dwyA 5eikA 5ek0A 5eulE 5ezmA 5fl7A
5fq6A 5fq6E 5fq7G 5gjvB 5gjvC 5gkoA 5gufA 5h1qA 5h2ft 5h36A
5hk1A 5i32A 5i6cA 5i6zA 5i9kA 5id3A 5ijiA 5irxE 5iwsA 5j4iA
5jkiA 5jnqA 5jrwA 5jsiA 5kbnA 5kc0A 5kkzB 5l22A 5l24A 5l25A
5l75F 5l75G 5lbjA 5ldvA 5lqxI 5lqxJ 5lqxU 5lqxY 5lweA 5lx9A
5m87A 5mdoA 5mg3C 5mg3D 5mg3E 5mg3Y 5mkkA 5mkkB 5mlzA 5mrwA
5mrwD 5n6mA 5n9yA 5naoA 5nj3A 5nupD 5nv9A 5o0tA 5o4uA 5o65A
5o8oA 5oc0A 5oc9A 5odwC 5oekA 5onuA 5oqtA 5osaA 5sv0A 5sv9A
5svjA 5t3rA 5t77A 5tcxA 5tsaA 5ttpA 5u1dX 5uz7R 5v56A 5v6pA
5v7pA 5v8kA 5vaiP 5vkvA 5vmsA 5wpqA 5wqcA 5wtrA 5wucA 5wufA
5x3xa 5x3xm 5x41F 5x5yA 5x5yF 5x5yG 5xlsA 5xnn4 5xu1M 5xw6A
5y78A 5y83A 5yckA 5zgbI 5zgbM 5zleA 6a2jA 6akfA 6aveA 6b3jA
6b3jG 6b3jR 6b85A 6b87A 6barA 6be1A 6bh8A 6bhpA 6bmsA 6btmA
6btmD 6btmE 6btmF 6bugA 6bugC 6bw6A 6c08C 6c14A 6c14B 6c261
6c262 6c263 6c265 6c26C 6c3iA 6c61C 6c61D 6c61M 6c61N 6c61O
6c70A 6c96A 6cb2A 6cfwA 6cfwB 6cfwC 6cfwD 6cfwF 6cfwH 6cfwK
6cfwL 6cfwN 6cjqA 6cmoA 6cp78 6cp7J 6cp7U 6cvlE 6d0jA 6dt0A
6e14C 6e14F 6e14G 6e14H 6e1kC 6e8wA 6edqA 6eu6A 6euqA 6eusA
6eyuA 6f0kA 6f0kC 6f0kD 6f0kE 6f0kH 6f2dA 6f34C 6f36A 6f36M
6f36N 6f46A 6f7hA 6ffvA 6fkfa 6fkfb 6fkfe 6fkfg 6fkfp 6fl9A
6fos2 6fos4 6gieA 6gpsA 6gs1A 6h3iB 6h3iF 6h7dA 6hd1A 6hraB
6hraC 6hraD 6hrbD 6hugC 6humA 6humB 6humD 6humE 6humF 6humG
6humI 6humK 6humN 6humO 6i6hA 6ic4G 6ic4I 6ic4K 6igz1 6igz3
6igz6 6igz9 6igzD 6igzF 6igzM 6iiuA 6iu3A 6mhyA 6mx4A 6mx4B
6mx4C 6n3qA 6n3qB 6n4iE 6nbxQ 6nd1A 6nhwA 6nt4B

#### 2.2 The test set (90 sequences)

1e7pC 1ezvH 1jb0L 1p4tA 1q16C 1rh5B 1s5lf 1tlwA 1xmeC 2b2fA
2hdfA 2ibzG 2k8jX 2kncB 2ksdA 2kseA 2latA 2losA 2lp1A 2n28A
2n90A 2onkE 2pnoA 2q7mA 2wscK 2wscL 2ww9M 2wwbC 2y69E 2y69F
2ysuB 2zxeG 3c02A 3dinD 3p5nA 3s0xA 3sljA 3tx3A 3vr8A 3vr8C
3w9tA 3wguE 4aq5A 4bwzA 4d6tG 4d6uD 4dblA 4e1tA 4epaA 4f35A
4heaW 4hkrA 4j9uE 4kppA 4mlbA 4q65A 4r1iA 4ri2A 4y25A 5aexA
5aymA 5azoA 5d0oD 5da0A 5edlA 5fgnA 5i20A 5u6oA 5uiwB 5v4sA
5wieB 5z62N 6a90B 6b3jP 6c6lB 6cfwG 6cmoR 6cnmA 6cp7Z 6cseC
6ddeR 6dmyB 6f0kF 6f2dG 6g79S 6gdgA 6idfE 6idpA 6meoG 6n3qF

#### 2.3 The validation set (90 sequences)

1c17M 1i78A 1kqfB 1nekC 1s5lb 1s5le 1s5lz 1tqqA 2bs2A 2bs2B
2k1kA 2l16A 2l6wA 2loqA 2m3eA 2mfrA 2mgyA 2mk9A 2mlhA 2nmrA
2r9rA 2voyJ 2wsc2 2ww9K 2xokH 2xutA 2yiuA 2zt9H 3anza 3cwbF
3dhwA 3dwoX 3fidA 3kziy 3mktA 3pl9A 3sn6B 3sy9A 3udcA 3uq4A
3ze3A 4bemJ 4dveA 4h33A 4he8C 4hzuS 4in5H 4j72A 4j7cI 4kjrA
4kt0K 4lepA 4o9uE 4p79A 4rngA 4ytpD 5b58T 5butA 5c6oA 5dl8A
5do7A 5ekeA 5gjvE 5kk2E 5mg3F 5mg3G 5n6hA 5ogeA 5oykA 5sy1A
5u1dB 5xj6A 5xnn3 5ys3A 5zazA 5zgbO 6cfwE 6cfwI 6cfwM 6cp77
6fkfd 6gctA 6humC 6humL 6humM 6humS 6ic4A 6n3qE 6n4bR 6nbqP

#### 3. The 327-protein dataset (327 sequences)

1czlA 1d1zA 1deoA 1dmtA 1e2bA 1e58A 1e7zA 1e8jA 1eayC 1edmB
1elwA 1f20A 1fe8A 1g8tA 1gmxA 1h0hB 1h0xA 1h2sB 1h4xA 1hqz1
1i1jA 1i3qL 1i4jA 1ijbA 1itvA 1j5pA 1jwpA 1k4jA 1khyA 1kqrA
1kt9A 1la6A 1lm5A 1m0uA 1m4jA 1mfzA 1mj4A 1mu7A 1nbaA 1nekC
1nofA 1nqdA 1occD 1of1A 1ow4A 1pc3A 1qd6C 1r4xA 1r77A 1regX
1rw1A 1s5lB 1t3jA 1td5A 1uajA 1uf4A 1uunA 1uv0A 1uw4B 1v54C
1v5vA 1v8eA 1vbkA 1vcaA 1vr5A 1wvgA 1ww9A 1xg1A 1xl3C 1ym0B
1yp0A 1yrkA 1yulA 1yvbI 1z0mA 1zv1A 2a67A 2a9yA 2acfA 2anvA
2auwA 2b0vA 2bhwA 2bjiA 2bl2A 2blnA 2bmjA 2c2qA 2c4jA 2ds2B
2fp8A 2fweA 2g02A 2gr8A 2hivA 2i6xA 2ijgX 2iuwA 2j58A 2j8bA
2jcbA 2jcvA 2jpsA 2jsoA 2kx6A 2mlrA 2nloA 2nooA 2ntkA 2o2wA
2o3iA 2p4xA 2p97A 2qorA 2r6gG 2rdcA 2rfqA 2rgxA 2v3dA 2vpwC
2w12A 2wfiA 2wieA 2wjqA 2x1qA 2xciA 2xd7A 2xqhA 2xykA 2y8wA
2zfnA 2zgiA 2ztbA 3a0tA 3a4kA 3a4tA 3as8A 3b8oA 3bduA 3c8cA
3cgmA 3cl6A 3cnbA 3cojA 3cq4A 3cwnA 3cxnA 3cz8A 3dr7A 3dtzA
3dwwA 3dxeB 3eikA 3emnX 3evaA 3f0hA 3f2oA 3fdhA 3fx3A 3g88A
3getA 3gmgA 3h9pA 3hm5A 3hw9A 3hzbA 3igqA 3igyB 3ih6A 3j5va
3k6oA 3ktoA 3ky8A 3l5oA 3la9A 3lg2A 3lgoA 3lm2A 3m6oA 3m8uA
3mdnA 3mvnA 3n0xA 3n1gC 3n2oA 3n6sA 3ne8A 3ngyA 3o15A 3o7hA
3og4B 3pgjA 3pmcA 3pt8B 3q47B 3q7bA 3qbtB 3qekA 3qp9A 3qvaA
3qvkA 3r2dA 3rauA 3rjzA 3rxyA 3s5qA 3sonA 3tlcA 3tn2A 3uc2A
3ungC 3up0A 3v5sA 3vusA 3vv1A 3w6kB 3w7vA 3w9sA 3wdcA 3wh4A
3wjdA 3wu2A 3wxmB 3ze3A 3zlcA 3zwzA 4abkA 4adgA 4afyA 4b21A
4b28A 4bjhB 4cbpA 4cccA 4ck4A 4cnfA 4czdB 4d01A 4d8fA 4dgtA
4e11A 4eacA 4emjB 4fdwA 4fmrA 4fmyA 4fs7A 4g10A 4g5aA 4gaxA
4ggvA 4guxA 4gvfA 4h2wA 4hc8A 4hd1A 4hesA 4hh8A 4hkiA 4hyoA
4ia5A 4ihfG 4im9A 4in5H 4infA 4ioxA 4iqzA 4iwnA 4ix7A 4iztA
4j1qA 4je9A 4jjjA 4jprA 4km8A 4l63A 4l8kA 4l8pA 4lclA 4lerA
4lkuA 4lpiA 4lreA 4lrjA 4lscA 4luqA 4luqC 4m2pA 4m6gA 4m7rA
4mcxB 4n0kA 4nhrA 4nlmA 4nooA 4nptA 4nssA 4nuaA 4odxA 4ohuA
4oloA 4otsA 4p0yA 4p3zA 4p7wA 4pk5A 4pnaA 4pwuA 4pzoA 4pzuA
4q0yA 4q1qA 4q4v3 4q53A 4q6mA 4qe0A 4qrzA 4r03A 4r9oA 4tshA
4u13A 4umlA 4weqA 4wuvA 4wwhA 4wxeA 7ahlA

#### 4. The 102 Pfam proteins (102 sequences)

1a3aA 1a6mA 1a70A 1aapA 1abaA 1ag6A 1aoeA 1atlA 1atzA 1avsA
1bdoA 1bebA 1behA 1brfA 1bsgA 1c44A 1c52A 1c9oA 1ckeA 1cxyA
1cznA 1d1qA 1dbxA 1dixA 1dlwA 1dsxA 1ej0A 1ek0A 1f6bA 1fk5A
1fl0A 1fnaA 1fqtA 1fvgA 1fvkA 1fx2A 1g9oA 1gbsA 1gmxA 1guuA
1gz2A 1gzcA 1h0pA 1h2eA 1h4xA 1h98A 1hfcA 1i1jA 1i1nA 1i4jA
1i5gA 1ihzA 1iibA 1im5A 1iwdA 1jbeA 1jbkA 1jfuA 1jfxA 1jkxA
1jo0A 1jo8A 1jvwA 1jwqA 1kidA 1kqrA 1ktgA 1kw4A 1lm4A 1lpyA
1m4jA 1m8aA 1mugA 1nb9A 1ne2A 1npsA 1nrvA 1ny1A 1o1zA 1pchA
1qf9A 1ql0A 1r26A 1roaA 1rw1A 1rybA 1smxA 1svyA 1tqhA 1tzvA
1vfyA 1vhuA 1vjkA 1vp6A 1xdzA 2cuaA 2hs1A 2phyA 3borA 3dqgA
4qyxA 5ptpA

#### 5. The dataset for the contact map predictor

##### 5.1 The training set (10054 sequences)

12asA 16vpA 1a12C 1a1xA 1a2xB 1a41A 1a5tA 1a62A 1a7jA 1a8lA
1a92B 1a9xF 1ae9A 1ah7A 1ahoA 1ahsA 1aluA 1alyA 1am7C 1aocB
1aolA 1aqzB 1atgA 1ax8A 1ayoB 1azoA 1b25C 1b33N 1b35A 1b35B
1b35C 1b35D 1b3uA 1b4fG 1b5eA 1b5qA 1b63A 1b77B 1b8kA 1b9wA
1bb1A 1bcoA 1bdfC 1beaA 1bg6A 1bgfA 1bheA 1bjaB 1bkrA 1bl0A
1bm8A 1bm9B 1bquB 1brtA 1bteA 1btkB 1bu8A 1bvyF 1bx7A 1bxyB
1byfA 1byiA 1byrA 1c0pA 1c1dA 1c1kA 1c1yB 1c3cA 1c4oA 1c4qD
1c5eB 1c7kA 1c8nA 1c8uA 1c94A 1c9kB 1cb8A 1cc8A 1ccwC 1ccwD
1cczA 1cfbA 1cfrA 1cfzC 1cg2D 1chdA 1chmB 1ci4A 1cjwA 1ckmB
1cnt3 1colB 1cq3B 1cr1A 1cr5A 1cshA 1ctfA 1cukA 1cv8A 1cwvA
1cxqA 1cxzB 1cy5A 1cywA 1d0dA 1d0qA 1d2nA 1d2oA 1d2sA 1d2tA
1d2zD 1d3bB 1d3bI 1d3yA 1d4aD 1d4oA 1d5tA 1d8hB 1d8wD 1d9cB
1dabA 1dbhA 1dbwB 1dcsA 1dd9A 1dekA 1devD 1dfuP 1dg6A 1dj0A
1dj7A 1dj8D 1dk8A 1dleB 1dm9A 1dmgA 1dmuA 1dowB 1dp4A 1dpjB
1dqgA 1dqpB 1ds1A 1dtdB 1dunA 1dvkB 1dvoA 1dwkE 1dxgB 1dynB
1dypA 1dzfA 1dzkA 1e0bB 1e2wA 1e30A 1e44B 1e5kA 1e6uA 1e7lA
1e9rF 1eajB 1eaqB 1eazA 1eb6A 1ed1A 1eejA 1eerA 1eerC 1eexA
1eexB 1eexM 1ef1C 1efdN 1egwC 1ehyB 1ei5A 1ei7B 1ei9A 1ej8A
1ejfB 1ek9B 1ekqA 1el6A 1elkB 1eluA 1em9B 1eokA 1epfB 1epuA
1es5A 1es9A 1escA 1et1B 1eteD 1euvA 1euvB 1euwA 1ev7B 1evlD
1evsA 1ew4A 1ewfA 1ez3B 1ezgB 1ezjA 1f00I 1f0kB 1f0lB 1f0xB
1f1eA 1f1mC 1f2lA 1f2tA 1f2tB 1f32A 1f3uC 1f3uH 1f3vA 1f46B
1f5nA 1f5qB 1f5vB 1f86A 1f8eA 1f9vA 1fc3B 1fc6A 1fcqA 1fcyA
1fg7A 1fi8C 1fi8D 1fjhB 1fjrB 1fm0D 1fm0E 1fm2B 1fn9A 1fnnB
1fo0B 1fo8A 1fobA 1fp2A 1fpoB 1fpzC 1fqjC 1fs0G 1fs7A 1fsgC
1ft5A 1ftrD 1fviA 1fxkC 1fyeA 1g12A 1g2rA 1g2yC 1g31E 1g3kA
1g5aA 1g5hA 1g5tA 1g60B 1g61A 1g66A 1g6xA 1g8aA 1g8eB 1g8mB
1g8pA 1ga6A 1gakA 1gciA 1gheA 1gjwA 1gk6A 1gk9A 1gk9B 1gl4A
1gmiA 1gmlC 1gmuA 1gnlB 1gnyA 1go3F 1go3M 1gp0A 1gp6A 1gpjA
1gppA 1gprA 1gqeA 1gs5A 1gs9A 1gsaA 1gsmA 1gttC 1gu2B 1gu3A
1gu4A 1gu7A 1guiA 1gutE 1gv9A 1gveB 1gvnB 1gvpA 1gweA 1gwmA
1gxmB 1gxrA 1gxuA 1gxyA 1gy7B 1gyxA 1gzsD 1h12A 1h21B 1h2bA
1h2cA 1h2kS 1h30A 1h3oA 1h3oB 1h4aX 1h5wB 1h6dK 1h6fB 1h6gA
1h6hA 1h6yB 1h80B 1h8pB 1h97B 1h99A 1hcuB 1hdoA 1he1B 1hf2C
1hfeT 1hh8A 1hi9B 1hj6A 1hk8A 1hlvA 1hp1A 1hq0A 1hr6F 1hruA
1htmE 1htrP 1htwA 1hulA 1huwA 1hw7A 1hx6B 1hx8A 1hxiA 1hxnA
1hxrB 1hyhA 1hyoA 1hz4A 1hz6A 1hztA 1i0rA 1i12B 1i1qA 1i1rB
1i24A 1i27A 1i2kA 1i2tA 1i36B 1i3jA 1i4dA 1i4uB 1i58A 1i5pA
1i60A 1i71A 1i7dA 1i8aA 1i8nB 1i9zA 1iapA 1iarB 1id0A 1id1B
1idpC 1ifrA 1igqC 1ii5A 1iktA 1im3P 1in4A 1inlD 1io0A 1iomA
1iq4A 1iq5B 1iqzA 1irqB 1is3A 1isuA 1it2B 1itbB 1ithB 1iujA
1iuqA 1ix9B 1ixhA 1ixlA 1ixvA 1izcA 1izmA 1j0pA 1j1tA 1j24A

1j27A 1j2jB 1j3aA 1j3wB 1j5uA 1j5wB 1j5yA 1j6rB 1j77A 1j7dA
1j7xA 1j83A 1j8bA 1jadB 1jayA 1jb0D 1jb0E 1jb0F 1jb0I 1jb0J
1jb0K 1jb0L 1jb0M 1jb0X 1jb3A 1jcdA 1jdcA 1jdhB 1jdwA 1je5B
1jetA 1jeyA 1jeyB 1jf3A 1jfbA 1jfuB 1jg1A 1jh6B 1jhcA 1jhgA
1jhjA 1jhsA 1ji7C 1jiwI 1jixA 1jkeB 1jl1A 1jm1A 1jmkC 1jmvA
1jniA 1jnrA 1jo0B 1jofC 1jogD 1josA 1jovA 1jqlB 1jr2B 1jr7A
1js8B 1jsuC 1jtvA 1ju2A 1juhD 1juvA 1jw9B 1jx6A 1jy2Q 1jy2R
1jy2S 1jyeA 1jyhA 1jykA 1jyoD 1jztA 1k0iA 1k1fF 1k1xB 1k3sA
1k3xA 1k4iA 1k4nA 1k4zB 1k5cA 1k5nA 1k5nB 1k6kA 1k77A 1k78I
1k7jA 1k7wC 1k8kC 1k8kD 1k8kE 1k8kF 1k8kG 1k8uA 1k8wA 1ka1A
1kaeA 1kafC 1kcfA 1kcmA 1kdgA 1kgdA 1kjnA 1kjqA 1kkoB 1klxA
1kmtB 1kngA 1knyB 1ko7A 1koeA 1kolA 1kp6A 1kpgC 1kptB 1kq3A
1kq6A 1kqpB 1kt6A 1ku3A 1kveB 1kveC 1kwfA 1kxgC 1kxoA 1kxpD
1kyfA 1kyqC 1kzqA 1l0sB 1l2pA 1l3kA 1l3pA 1l4dB 1l6pA 1l6rB
1l7aA 1l8dA 1l9lA 1l9xC 1lb6A 1lbqB 1lbuA 1lc0A 1lc5A 1lf6B
1lf7A 1lfpA 1lfwA 1lj2A 1lj8A 1lkiA 1llfA 1lmiA 1lmlA 1ln1A
1lniB 1lo7A 1lpbA 1lq9A 1lqtB 1lr0A 1lr5B 1lrzA 1ls1A 1lu0A
1lu4A 1luaB 1lucA 1luzB 1lv7A 1lwbA 1lxjA 1ly1A 1lzlA 1m0dC
1m0kA 1m0wB 1m15A 1m1eB 1m1fA 1m1qA 1m1sA 1m1zB 1m22A 1m2dA
1m32D 1m4iB 1m4lA 1m4uA 1m55A 1m56D 1m5iA 1m5qO 1m6yB 1m70D
1m9zA 1maiA 1mc2A 1mgtA 1mijA 1mixA 1mj5A 1mjnA 1mk0A 1mk4B
1mkfB 1mkkB 1mn8C 1mnnA 1mo9B 1mpgA 1mr7A 1mrzA 1ms9A 1mscA
1mspB 1mtpA 1mtyE 1mtzA 1mujC 1munA 1musA 1muwA 1mvlA 1mw9X
1mwqB 1mwwB 1mxrA 1my7B 1mz9E 1mzwB 1n08B 1n0wB 1n12C 1n1cB
1n1fA 1n26A 1n3lA 1n4kA 1n57A 1n62A 1n62C 1n67A 1n71B 1n7kB
1n7sC 1n7vA 1n7zA 1n81A 1n8vB 1n93X 1n9pA 1na6A 1nbwD 1nc7D
1ne2B 1nepA 1nezG 1nfpA 1ng6A 1nh1A 1nigA 1nijA 1njhA 1njrA
1nkdA 1nkgA 1nkiA 1nkoA 1nkrA 1nkzF 1nlfA 1nlqB 1nnfA 1nnlB
1nnwB 1nnxA 1np0B 1np6A 1nqkA 1nqzA 1nr0A 1nrjA 1nrjB 1nszA
1ntvA 1nu0A 1nu7H 1nulA 1nuuA 1nuyA 1nv8B 1nxmA 1nxuB 1nycA
1nz0C 1nzjA 1nznA 1o13A 1o22A 1o4wA 1o50A 1o57B 1o5lA 1o5uA
1o66D 1o69A 1o6dA 1o6vA 1o75B 1o7fA 1o7iA 1o7jC 1o7zA 1o88A
1o8xA 1o98A 1o9gA 1o9yA 1oa8B 1oaiA 1oc0B 1ocyA 1od3A 1odmA
1oeyJ 1of8B 1ogdD 1ogoX 1oh0B 1oh2Q 1oh4A 1ohuB 1oi0D 1oi2A
1oi7A 1oisA 1oj5A 1ojhH 1ok0A 1ok3B 1okcA 1oksA 1oltA 1olzB
1omzB 1on2B 1oo0A 1oo0B 1ooeB 1oq1D 1oqjA 1or4B 1orsC 1oruB
1otkA 1ou8B 1ov9B 1ow1A 1oygA 1oyzA 1oz2A 1oz9A 1p1jA 1p1mA
1p1xB 1p2xA 1p32C 1p35C 1p3cA 1p3dB 1p3hD 1p3qQ 1p4xA 1p57A
1p5vB 1p5zB 1p6oB 1p90A 1p9gA 1p9hA 1p9iA 1p9yB 1paqA 1pbjA
1pbwB 1pcfE 1pdkB 1pdoA 1peaA 1pf5A 1pg6A 1pixA 1pkhB 1pl5A
1pm4B 1pmhX 1pn0D 1po5A 1pocA 1poiD 1pp0D 1pp7U 1ppjI 1ppjO
1ppjT 1psrB 1pswA 1pucA 1pujA 1pv5A 1pvgB 1pvmB 1pxzB 1pytA
1pz4A 1pzwA 1q08A 1q0pA 1q0rA 1q15C 1q1fA 1q2hC 1q33A 1q35A
1q40D 1q42A 1q5yC 1q5zA 1q67A 1q6oB 1q6zA 1q74D 1q7eA 1q7fA

1q7lB 1q7lC 1q87B 1q8bA 1q8cA 1q8dA 1qamA 1qazA 1qcsA 1qcxA
1qd1B 1qexA 1qf8A 1qfhA 1qfjC 1qftB 1qg8A 1qgrB 1qhlA 1qhxA
1qjpA 1qkrA 1qksA 1qlwA 1qnrA 1qq5A 1qqp1 1qqp2 1qqp4 1qqrC
1qr0A 1qsaA 1qsmC 1qstA 1quuA 1qv1A 1qv9A 1qw2A 1qw9B 1qwdB
1qwgA 1qwrA 1qyiA 1qynA 1qysA 1qz7B 1qzmA 1r0dH 1r0mD 1r0uA
1r29A 1r2jA 1r44A 1r45A 1r4vA 1r5mA 1r6dA 1r6jA 1r6wA 1r6xA
1r75A 1r7jA 1r89A 1r8eA 1r8gB 1r9lA 1r9wA 1ra0A 1rcwB 1reqB
1rf6C 1rfyB 1rg8B 1rgxB 1rh6A 1ri5A 1ri6A 1rifA 1rj7M 1rjdA
1rjuV 1rk6A 1rk8C 1rkiA 1rkqB 1rktB 1rkuB 1rl0A 1rl6A 1rliB
1rlzA 1rmgA 1ro2A 1ro5A 1rocA 1rp0A 1rp3F 1rr7A 1rsgB 1rssA
1rt8A 1rtqA 1rttA 1rtwB 1rutX 1rv9A 1rwzA 1rxqD 1ry9C 1ryiB
1rylB 1ryoA 1rypN 1ryqA 1rz3A 1rz4A 1rzhM 1s0pB 1s12D 1s1dA
1s21A 1s29A 1s2oA 1s2wA 1s2xA 1s3cA 1s4cC 1s4kB 1s5uF 1s7iA
1s7zA 1s98B 1s99A 1s9rA 1s9uA 1sauA 1sazA 1sbxA 1sbyB 1scfC
1sd4B 1sdiA 1sdoA 1se8A 1sedC 1sefA 1seiB 1senA 1sfpA 1sfsA
1sfuA 1sg4C 1sg6B 1sgmA 1sgwA 1sh8A 1shuX 1sj1A 1smxB 1so7A
1sp3A 1sq9A 1sqhA 1sqwA 1sr4C 1sr8A 1sraA 1ss4A 1stmA 1stzA
1su1A 1sumB 1svfA 1svfB 1svmA 1svsA 1sw6B 1sz7A 1szhB 1szwA
1t07A 1t08A 1t0bB 1t0fB 1t0iB 1t0pB 1t1jA 1t1uA 1t33B 1t3gA
1t3yA 1t4aB 1t4wA 1t61E 1t6aA 1t6cA 1t6fA 1t6lA 1t6sB 1t6t1
1t6uE 1t77C 1t82D 1t92A 1t9iB 1tafA 1tafB 1tbfA 1tc3C 1te5A
1tfeA 1tfzA 1tg0A 1th7J 1thtB 1tifA 1tigA 1tiqA 1tjlA 1tkeA
1tkjA 1tp6A 1tq5A 1tqgA 1tqyF 1ts9A 1tt8A 1tu1A 1tu9A 1tuaA
1tueF 1tuhA 1tulA 1tuvA 1tuwA 1tv8A 1tvfA 1tvgA 1twdA 1twfF
1twfH 1twfI 1twfJ 1twfK 1twuA 1txgA 1txkA 1tzpB 1u02A 1u07B
1u0sA 1u14A 1u2hA 1u2kA 1u2mC 1u2wA 1u2zB 1u53A 1u5fA 1u5kA
1u5uB 1u60C 1u6tA 1u7bA 1u7gA 1u7iA 1u7kE 1u7lA 1u7pD 1u7zA
1u84A 1u8sA 1u8vC 1u8xX 1u9kA 1u9lB 1u9pA 1ua4A 1ua7A 1ub9A
1ucdA 1ucrA 1ucsA 1ud9B 1uebB 1uekA 1ufhA 1ufiD 1ufoE 1ufyA
1ugiF 1uhvD 1ui0A 1ui5A 1uixA 1uj2A 1uj8A 1ujcA 1ukfA 1uoyA
1upkA 1upsA 1uptF 1urqA 1us0A 1us5A 1uscB 1usgA 1usmA 1usuB
1ut7B 1utgA 1utyA 1uujC 1uuyA 1uv7A 1uvjB 1uvqC 1uw4A 1uwcB
1uwkB 1uwvA 1ux5A 1ux6A 1uxoA 1uz3B 1v05A 1v0aA 1v0wA 1v18B
1v2xA 1v2zA 1v30A 1v33A 1v4aA 1v4eB 1v4pC 1v6pB 1v6tA 1v70A
1v72A 1v74A 1v74B 1v77A 1v7zA 1v84B 1v8dA 1v8hB 1v96A 1v9fA
1v9mA 1v9yB 1vajA 1vbwA 1vccA 1vchE 1vctA 1vdwA 1ve2A 1ve4A
1vf7L 1vg0A 1vgwD 1vh4B 1vh5A 1vhnA 1vhtB 1vhzA 1vi0A 1vi4A
1vi6A 1vimB 1vjgA 1vjlB 1vjqB 1vjvA 1vjxA 1vk3A 1vk4A 1vk6A
1vkeE 1vkfB 1vkhA 1vkiA 1vkkA 1vkwA 1vkyB 1vl5C 1vl7A 1vlmA
1vlyA 1vmbA 1vmgA 1vmhA 1vmoA 1vp7E 1vp8A 1vpmB 1vpqA 1vprA
1vptA 1vqo1 1vqoP 1vqoR 1vqoU 1vqqB 1vqrC 1vqsB 1vr4D 1vr7B
1vr8A 1vraB 1vsrA 1vybA 1vyiA 1vykA 1vyrA 1vz0C 1vzjF 1vzyB
1w07A 1w0hA 1w0nA 1w1hC 1w23A 1w2fB 1w2wI 1w2wN 1w4sA 1w53A
1w5qB 1w5rB 1w66A 1w6sC 1w6sD 1w8iA 1w8sF 1w99A 1wb4B 1wbaA

1wbhC 1wcv1 1wcwA 1wd3A 1wd5A 1wdcB 1wdjC 1wehA 1werA 1whiA
1whzA 1wiwA 1wjxA 1wkcA 1wkqB 1wleB 1wlfA 1wlgA 1wljA 1wluA
1wlzB 1wm1A 1wmhA 1wmhB 1wmiC 1wmiD 1wmxA 1wn2A 1wnaA 1wolA
1wouA 1wpbK 1wpnB 1wq6A 1wqlB 1wrdA 1ws8B 1wt6D 1wteB 1wtjB
1wurA 1wv3A 1wv9A 1wvfA 1wvhA 1wwcA 1wwiA 1wwpB 1wwzA 1wy5B
1wy6A 1wyzB 1wz3A 1wzdB 1wznA 1x0tA 1x19A 1x2iB 1x3kA 1x3lA
1x54A 1x6iB 1x6mD 1x6oA 1x6zA 1x79A 1x7dA 1x8bA 1x8dC 1x8qA
1x91A 1x9iB 1x9zA 1xakA 1xauA 1xawA 1xbiA 1xcrB 1xd3A 1xdnA
1xdpB 1xdyB 1xe1A 1xe7B 1xffA 1xfkA 1xfsA 1xg5D 1xg8A 1xgkA
1xhdA 1xhnB 1xipA 1xiwA 1xiwF 1xjuB 1xjvA 1xk5A 1xk7A 1xkiA
1xkpA 1xkrA 1xksA 1xkwA 1xlqA 1xm7B 1xmkA 1xmtA 1xmxA 1xnfA
1xo0B 1xo5A 1xovA 1xpjD 1xppA 1xqoA 1xqrB 1xr4A 1xrkB 1xruA
1xrxA 1xs0C 1xsvB 1xszA 1xteA 1xttC 1xu9D 1xubA 1xw3A 1xwvB
1xzpB 1y02A 1y08A 1y0hB 1y0kA 1y0nA 1y0uB 1y14A 1y1pA 1y1xA
1y3tB 1y43A 1y43B 1y4mA 1y5hA 1y60C 1y63A 1y66A 1y6xA 1y6zB
1y71A 1y7pC 1y7rA 1y88A 1y8aA 1y8xB 1y96A 1y96B 1y9iD 1y9lA
1y9wB 1yacB 1yadC 1yb2A 1ybkB 1yc9A 1ycdB 1yd0A 1yd7A 1ydxA
1ydyA 1ye8A 1yf2A 1yf3A 1yfqA 1ygtA 1yhtA 1yi9A 1yixA 1yj7A
1yk3A 1ykiB 1yleA 1yllD 1ylmA 1yloD 1ylqA 1ylxA 1yn3A 1ynbB
1ynfA 1ynpB 1yocA 1yodA 1yozB 1ypyA 1yq5B 1yqhA 1yqsA 1yqtA
1yrbA 1yreA 1yrrA 1yrtA 1ys1X 1yt3A 1yt5B 1ytlC 1ytvM 1yu0A
1yw4A 1ywfA 1ywmA 1yx1B 1yymS 1yzvA 1yzyA 1z0pA 1z0sA 1z0wA
1z0xB 1z1sA 1z21A 1z2nX 1z2uA 1z3eA 1z3eB 1z3xA 1z45A 1z67A
1z6nA 1z6oB 1z72B 1z84A 1z85B 1z8uC 1z94D 1z9lA 1za0A 1zarA
1zatA 1zavA 1zavW 1zb1B 1zbpA 1zbsA 1zbxB 1zc3B 1zccB 1zceA
1zd0A 1ze3H 1zeeA 1zelA 1zghA 1zgkA 1zh8B 1zhsE 1zhvA 1zhxA
1zi8A 1zjcA 1zk4A 1zk5A 1zk8B 1zkeC 1zkpA 1zl0A 1zmaA 1zn6A
1zoqD 1zpsA 1zpvA 1zr6A 1zs9A 1zsoA 1zsqA 1zt3A 1ztcC 1ztdB
1zuoA 1zupB 1zvaA 1zvpC 1zvtA 1zwwA 1zx3A 1zx8A 1zxxA 1zy7B
1zybA 1zymA 1zyoA 1zzkA 2a0bA 2a14A 2a15A 2a1iA 2a1kA 2a1vA
2a1xA 2a26A 2a2fX 2a2mA 2a35B 2a3nA 2a4xB 2a5dB 2a5hD 2a5zA
2a65A 2a6aA 2a6qD 2a6sB 2a6zA 2a7kD 2a8yD 2a90A 2a9iA 2a9uA
2aalB 2aamA 2ab5B 2abwB 2ae6C 2aefA 2aegC 2aeuA 2ag4B 2agkA
2ahuB 2aibA 2aj6A 2ajrA 2akfC 2akzA 2al6A 2amhA 2amlB 2aneD
2anrA 2anuD 2ao9I 2ap3A 2aplA 2aq6A 2arcA 2asbA 2asfA 2askB
2atpF 2atzA 2au5A 2auaA 2awiB 2axcA 2axoA 2axpA 2axqA 2axwA
2aydA 2az4A 2azjB 2azwA 2b06A 2b0aA 2b0cA 2b1eA 2b1yA 2b2aB
2b3gB 2b4aA 2b4hB 2b4wA 2b5iD 2b69A 2b78A 2b81C 2b82A 2b8iA
2b8mA 2b97A 2b99D 2b9cA 2b9dB 2b9sB 2b9wA 2ba2C 2bayF 2bb3B
2bbdA 2bbhA 2bbrA 2bcmB 2bcxB 2bdrB 2bdtA 2bdvA 2bf6A 2bfdA
2bfdB 2bfwA 2bhgA 2bhuA 2bibA 2biiA 2bjnA 2bjqA 2bjuA 2bk8A
2bk9A 2bkfA 2bkrA 2bkxA 2bl0A 2bl0B 2bl8C 2bl9A 2bm5A 2bm8K
2bmoA 2bmoB 2bnlE 2bnmB 2bo4E 2bo9B 2bogX 2bolB 2bonA 2bopA
2bryA 2bs2C 2bsjA 2bsyA 2bt9C 2bu3A 2bueA 2bvfB 2bw3A 2bwrA

2bz1A 2bzvA 2bzwB 2c0cA 2c0gB 2c0nA 2c0zA 2c1lB 2c1mB 2c1vA
2c1wB 2c2iB 2c2uA 2c36A 2c3nB 2c3vA 2c5aA 2c61B 2c6uA 2c71A
2c7nI 2c92E 2c9qA 2ca5A 2carB 2cayB 2cb8A 2cc6A 2ccmB 2ccqA
2ccvA 2cdcB 2ce2X 2cfmA 2cg7A 2ch5C 2ch7A 2chcA 2chgC 2chpD
2cibA 2ciuA 2cj4A 2cjaB 2cjjA 2cjtD 2ckkA 2clbC 2cmgB 2cnqA
2co3A 2covD 2cpgB 2cs7B 2cu3A 2culA 2cvbA 2cvdD 2cveA 2cviB
2cw6C 2cw9A 2cwqB 2cwsA 2cwyA 2cx7B 2cxaA 2cxcA 2cxhA 2cxiA
2cxkD 2cxnB 2cxyA 2cy5A 2cyeC 2cz4C 2czsB 2d00C 2d0bA 2d0oC
2d1gB 2d1lB 2d28C 2d3dA 2d42B 2d48A 2d4gB 2d4pA 2d59A 2d5bA
2d5mA 2d5wA 2d68B 2d6yA 2d7eD 2d7jA 2d7vB 2d81A 2d8dB 2d9rA
2db0A 2db7A 2dbnA 2dbsA 2dbyA 2dc1B 2dc4B 2dchX 2dclA 2ddfB
2ddmB 2ddrB 2dduA 2ddxA 2ddzE 2de3A 2dejA 2dg1F 2dg7A 2dg8A
2dgkD 2dh2A 2dhoA 2djfA 2dkaB 2dkjA 2dlbB 2dm9A 2dohC 2dokB
2dp9A 2dpfD 2dplB 2dpmA 2dqlB 2ds5A 2dskA 2dstA 2dtjA 2dulA
2dvmC 2dwkA 2dwuC 2dxaA 2dxqB 2dxuA 2dy0A 2dy1A 2dyiA 2dyjB
2dyoA 2dyoB 2e0nB 2e11B 2e12A 2e1fA 2e1vA 2e2aB 2e2eA 2e2oA
2e3hA 2e3nA 2e4tA 2e56A 2e5aA 2e5fA 2e5yA 2e6fB 2e6mA 2e6xD
2e7jA 2e7vA 2e8bA 2e8eA 2e8gA 2e8vA 2e9xB 2e9xE 2e9xH 2eaqA
2ebbA 2ebeB 2eceA 2ecuA 2ed6L 2efeC 2efkA 2efvA 2egvA 2eh3A
2ehbD 2ehgA 2ehpB 2ehwC 2ehzA 2ei9A 2eiyC 2ej8A 2ejxA 2ek0B
2ekdE 2elcD 2endA 2engA 2eplX 2eq5B 2erfA 2erlA 2ervA 2es9A
2essA 2et1A 2etbA 2ethB 2etjA 2etsA 2eu9A 2eucA 2ev1B 2ew0A
2ewcC 2ewfA 2ex2A 2ex5A 2eyuA 2ez2A 2ezvB 2f01B 2f06A 2f07B
2f0cB 2f1fB 2f1nA 2f22B 2f23A 2f2bA 2f2fD 2f3lA 2f46B 2f48A
2f4mA 2f4mB 2f4nA 2f4pC 2f4wB 2f5gB 2f5jA 2f5tX 2f5uA 2f60K
2f62A 2f6hX 2f6kA 2f6mA 2f6mD 2f6rA 2f7vA 2f8lA 2f8yA 2f9hA
2f9iA 2fa1B 2fa5B 2fa8A 2faoA 2fb0A 2fb5A 2fb6A 2fbaA 2fbiA
2fbqA 2fcaB 2fcjA 2fckA 2fcoA 2fctB 2fcwA 2fcwB 2fd4A 2fd5A
2fdbN 2fdoA 2fdrA 2feaA 2fefB 2felB 2fexA 2ff4A 2ffgA 2ffsA
2fg9A 2fgqX 2fgtA 2fh5A 2fhpA 2fhzA 2fhzB 2fi0A 2fi1A 2fi9A
2fipE 2fiuB 2fiyB 2fj8A 2fji1 2fjrA 2fkkA 2fl4A 2fm9A 2fmaA
2fmyC 2fnaA 2fnjB 2fnoB 2fnuA 2fotC 2fozA 2fp1A 2fphX 2fpnA
2fq3A 2fq4A 2fqmF 2fqxA 2fr5C 2freA 2frgP 2fsdB 2fsjA 2fsqA
2fsrA 2ft0B 2ftrA 2fu2A 2fueA 2fulB 2fupA 2furA 2fvvA 2fvyA
2fx5A 2fxaD 2fy7A 2fyfB 2fygA 2fyuK 2fzpA 2fztB 2fzvA 2g0cA
2g0wA 2g1uA 2g2cA 2g2qB 2g30A 2g3aA 2g3rA 2g3wB 2g40A 2g45A
2g5gX 2g7gA 2g7lA 2g7oA 2g7sA 2g8lB 2g8sB 2g9wB 2ga1A 2gagB
2gagC 2gagD 2gakA 2ganB 2gauA 2gaxB 2gb4B 2gbbC 2gboA 2gc7H
2gd9B 2gdmA 2gdqB 2ge7B 2gefA 2genA 2geyC 2gf3A 2gf4B 2gf6B
2gffA 2gfhA 2gfqB 2ggcA 2ghsA 2ghvC 2giaA 2giaB 2giyB 2gj2D
2gj3B 2gk4A 2gkeA 2gkgA 2gkpA 2glzB 2gm3E 2gmhB 2gmqA 2gmyF
2gnoA 2gnpA 2go8A 2gomA 2gpeD 2gpiA 2gqtA 2gqwA 2grrB 2grvB
2gs5A 2gscB 2gsoA 2gsvA 2gt1B 2gtiA 2gu3A 2gu9B 2gufA 2guhB
2guiA 2gukB 2gumB 2gupA 2guzK 2gv8B 2gviA 2gwdA 2gwfA 2gwgB

2gwmA 2gxgA 2gxqA 2gyqB 2gz4C 2gz6A 2gzqA 2gzsA 2gzvA 2h1tB
2h1vA 2h21A 2h30A 2h5cA 2h5nB 2h62C 2h6lA 2h7oA 2h88B 2h88C
2h88N 2h88Q 2h8eA 2h8gB 2h9aA 2h9aB 2h9bA 2ha8A 2hb0A 2hbaB
2hboA 2hbvA 2hbwA 2hc8A 2hcfA 2hd0G 2hd9A 2hdiB 2hdoA 2heuB
2hewF 2hf1B 2hf2A 2hfsA 2hhcA 2hhzA 2hi0A 2himA 2hinB 2hiqB
2hiyA 2hj1B 2hjeA 2hkjA 2hkuA 2hkvA 2hl7A 2hljA 2hlyA 2hlzC
2hmaA 2hmvB 2hnfA 2hngA 2hnuC 2hoxD 2hp0A 2hpsA 2hq4B 2hq7B
2hq9A 2hqlD 2hqsA 2hqtD 2hqxB 2hqyA 2hr3C 2hraA 2hsbA 2htdA
2htiA 2hu9B 2hueB 2huhA 2hujA 2hvwC 2hw2A 2hw4A 2hx0A 2hx1B
2hx5A 2hxiB 2hxrA 2hxtA 2hy5B 2hy5C 2hy7A 2hyjA 2hytA 2hzfA
2hzmB 2hzmC 2i00B 2i02A 2i06A 2i0zA 2i10A 2i15B 2i1sA 2i2cA
2i2oB 2i2xG 2i2xJ 2i39B 2i3dB 2i3oC 2i3sF 2i44C 2i49A 2i4lA
2i51B 2i53A 2i5eA 2i5hA 2i5iA 2i5uA 2i5vO 2i6hA 2i6tB 2i6vA
2i71A 2i74B 2i79F 2i7gA 2i7hD 2i7rA 2i8dA 2i8tB 2i9cA 2i9dC
2i9fD 2i9iA 2i9wA 2i9xA 2ia1A 2ia7A 2iabA 2iafA 2iayA 2iazD
2ib0B 2ibdB 2ibnB 2ic2B 2ic6A 2ichA 2id6A 2idlB 2idoD 2ielB
2if6A 2ig6B 2ig8A 2igiB 2igpA 2igsE 2igtB 2ihtD 2ii0A 2ii2A
2iidB 2iihA 2ij2A 2ijlB 2ijqB 2ikkB 2iksA 2il5A 2ilkA 2ilrA
2imfA 2imhB 2imjB 2imqX 2imrA 2in3A 2in5B 2inuA 2inwB 2iojA
2ip1A 2ip6A 2ipqX 2iqcA 2irpB 2isbA 2iskG 2it2A 2it9A 2iu5B
2iumC 2iuyA 2ivfC 2ivnA 2ivyA 2iw1A 2ixaA 2ixmA 2ixsA 2iyvA
2iz6B 2izrA 2izxB 2j0aA 2j0sT 2j1vA 2j2jC 2j3tC 2j3tD 2j3wC
2j3wF 2j43A 2j5bA 2j5gD 2j5yA 2j66A 2j6aA 2j6bA 2j6lE 2j6pE
2j73B 2j7qA 2j85A 2j8gA 2j8kA 2j91B 2j97A 2j9oC 2j9wA 2ja4A
2ja9A 2jaeB 2jbwD 2jbyA 2jc9A 2jdaA 2jdcA 2jdjB 2je6A 2je6B
2je6I 2jekA 2jenA 2jfrA 2jg0A 2jg1C 2jgpA 2jgsB 2jh1A 2jh3C
2jheA 2jhfB 2jhnA 2jhpA 2jiiA 2jisB 2jj7A 2jjuA 2jkhL 2jkuA
2jliA 2mcmA 2mhrA 2mprB 2nl9A 2nlrA 2nlsA 2nlvA 2nlyA 2nmlA
2nn4A 2nn5A 2nnuA 2nogA 2np5D 2npsC 2nptC 2nq2A 2nq3A 2nqwA
2nr4A 2nr5H 2nr7A 2nrhB 2nrjA 2nrkA 2nrrA 2ns0A 2ns6A 2ns9B
2nsaA 2nscA 2nsfA 2nszA 2nteB 2ntpA 2ntxA 2nuhA 2nujA 2nutC
2nvhA 2nvoA 2nvwB 2nw8A 2nwfA 2nwhA 2nwiF 2nwrB 2nx2A 2nx4B
2nx9A 2nxfA 2nxoC 2nxpF 2nxvB 2nxwA 2nyiB 2nykA 2nyvA 2nz7A
2nzcD 2nzlA 2nzxC 2o0aA 2o0jA 2o0mA 2o0qA 2o14A 2o1kB 2o1qB
2o1sB 2o2gA 2o2kA 2o34B 2o3bB 2o3jC 2o4aA 2o4tA 2o57D 2o5fB
2o5hB 2o5nB 2o62A 2o6kB 2o6lA 2o6pB 2o70D 2o71A 2o7iA 2o7rA
2o7tA 2o8gJ 2o8lA 2o8pA 2o8qA 2o8sB 2o90A 2o9uX 2oaaB 2oafB
2ob0B 2ob5A 2ob9A 2obbA 2obdA 2obnC 2obpA 2ocgA 2octB 2od0A
2od4A 2od5A 2od6C 2odfF 2odkB 2odvA 2oebA 2oeeA 2oerA 2oezB
2of3A 2ofkB 2ofyB 2ofzA 2ogfC 2oggA 2ogiB 2oh1D 2oh3A 2ohwA
2oikB 2oitA 2oixA 2oj5E 2ojhA 2ojwE 2okfA 2okmA 2oktA 2okuB
2ol5A 2olmA 2olnA 2olrA 2oltB 2om2B 2om6B 2omlA 2omzB 2onfA
2ooaB 2oocA 2ookA 2opcA 2opeA 2opiB 2opwA 2oqbB 2oqmD 2orwA
2oryB 2osoA 2osxA 2ou1K 2ou3A 2ou6A 2ougB 2ouxB 2ov0A 2ov9B

2ovgA 2ovjA 2ovsB 2owlB 2ownA 2owpA 2ox6D 2ox7B 2oxgD 2oxlA
2oxoA 2oy9B 2oyaA 2oyoB 2oyyD 2oz8A 2ozeA 2ozgA 2ozhA 2ozjA
2ozlC 2oznA 2oznB 2oztA 2ozvB 2ozzA 2p09A 2p0aB 2p0bA 2p0lA
2p0mB 2p0nA 2p0oA 2p0sB 2p0tA 2p0wA 2p10B 2p11B 2p12B 2p14A
2p1mB 2p25A 2p2sB 2p35A 2p38A 2p3pB 2p4fA 2p4hX 2p4oA 2p4zA
2p51A 2p57A 2p58A 2p58B 2p58C 2p5kA 2p5vC 2p62A 2p64A 2p65A
2p6wA 2p75B 2p7iB 2p8iB 2p8jA 2p8qB 2p8rA 2p8tA 2p9wA 2p9xA
2pa7B 2pagA 2pblD 2pc1A 2pcsA 2pd1D 2pd2A 2pefA 2petA 2pezB
2pfiA 2pftA 2pfwA 2pfzA 2pg3A 2pgeA 2pgnB 2phnB 2phpE 2piaA
2pjdA 2pjpA 2pjuD 2pk8A 2pkdF 2pkeB 2pkfA 2pliD 2plxB 2pm7C
2pmaA 2pmyA 2pn1A 2pn2A 2pndA 2pneA 2pnlJ 2pnqA 2pnwA 2pocB
2pofA 2pomA 2porA 2ppqA 2ppvA 2pq7A 2pq8A 2pqvB 2pqxA 2pr7B
2prsB 2prvB 2prxB 2ps5B 2psbA 2pspA 2pstX 2ptrB 2pttB 2pu3A
2puzB 2pv2D 2pv4A 2pv7B 2pw9D 2pwwA 2pxxA 2py6A 2pyqD 2pywA
2pyxB 2q00B 2q01C 2q02D 2q03A 2q04F 2q07A 2q0tA 2q0xB 2q0yA
2q12A 2q18X 2q1wB 2q1zD 2q22C 2q24B 2q28B 2q2fA 2q35A 2q3qA
2q3sF 2q3tA 2q40A 2q43A 2q48A 2q4aB 2q4kC 2q4mA 2q4oB 2q4uA
2q4wA 2q4xB 2q4zB 2q58B 2q5cA 2q66A 2q6kA 2q6qB 2q78D 2q7aB
2q7fA 2q7sB 2q82A 2q83A 2q88A 2q8kA 2q8pA 2q8vB 2q9kA 2q9rA
2q9uB 2qb7B 2qcpX 2qcuB 2qcvA 2qdfA 2qdjA 2qdlB 2qdqA 2qe8B
2qe9B 2qeaA 2qecA 2qedA 2qeuC 2qevA 2qf4A 2qfaB 2qfaC 2qfeA
2qg7D 2qgoA 2qgyB 2qh9A 2qhfA 2qhoH 2qhpA 2qhqB 2qibA 2qihB
2qipA 2qiyA 2qiyC 2qjfB 2qjlA 2qjtB 2qjwD 2qjzB 2qk1A 2qkdA
2qkoB 2qkpC 2ql3B 2ql8A 2qlcE 2qltA 2qlzB 2qm4C 2qmlA 2qmqA
2qn6B 2qngA 2qniA 2qnlA 2qntA 2qolA 2qp2A 2qpvB 2qq4B 2qq8A
2qqyA 2qr4B 2qr6A 2qrdA 2qrlA 2qruA 2qryC 2qs9A 2qsbA 2qsdH
2qsfA 2qsiB 2qskA 2qsvA 2qswA 2qsxB 2qt1A 2qtdA 2qtqC 2qtsD
2qtvD 2qu7A 2qu8A 2qudB 2qupA 2qv0A 2qv3A 2qv5B 2qv6B 2qvgA
2qvpC 2qw5B 2qwuA 2qwwB 2qwzB 2qx5B 2qxfA 2qy6A 2qyaC 2qybA
2qycB 2qyfD 2qyvA 2qzbB 2qzcA 2qziD 2qzqA 2qzuA 2r01A 2r0lB
2r0xA 2r16A 2r19A 2r1iB 2r1jR 2r25A 2r2aB 2r2cA 2r2dF 2r2zA
2r39A 2r3sA 2r44A 2r4fD 2r4gA 2r4iB 2r51A 2r5oA 2r5sA 2r5uA
2r60A 2r6jB 2r6vA 2r6zA 2r78A 2r7gC 2r85B 2r8wB 2r91C 2r9fA
2r9qB 2ra1A 2ra9A 2raaA 2raeA 2rafB 2rasB 2rauA 2rb7A 2rbcA
2rbdB 2rbgB 2rbkA 2rc3D 2rccA 2rciA 2rd7A 2rd9B 2rdeA 2rdgA
2rdqA 2re2A 2reeA 2remC 2retE 2reuA 2reyA 2rffA 2rfrA 2rg4A
2rg8A 2rgqA 2rgyA 2rh0A 2rh2A 2rh3A 2rhfA 2rhmA 2rhwA 2rieC
2rihA 2rinB 2riqA 2rj2A 2rjiB 2rjoA 2rk9B 2rkhA 2rklC 2rknA
2rkqA 2rkvA 2rl8B 2rldC 2sakA 2scpB 2spcB 2sqcB 2trcP 2uu8A
2uurA 2uuzB 2uv4A 2uvfB 2uvkA 2uvpA 2uw1A 2uwjE 2ux1I 2uy1A
2uyoA 2uytA 2uz0A 2uz1A 2v03A 2v0hA 2v0oB 2v0pB 2v0xB 2v1qB
2v1yA 2v25A 2v33B 2v3aA 2v3gA 2v3kA 2v4xA 2v52M 2v54B 2v57D
2v66D 2v6kB 2v6vB 2v76C 2v79A 2v7fA 2v7kA 2v7sA 2v84A 2v8iA
2v8qA 2v8qB 2v8tB 2v94A 2v9kA 2v9lA 2v9vA 2vakA 2vb1A 2vbkA

2vbuA 2vc8A 2vchA 2vclA 2vdfA 2vdjA 2vduD 2ve3A 2ve8F 2veqA
2vfoA 2vfrA 2vfxH 2vhaA 2vjwA 2vk2A 2vk8C 2vkjB 2vlaA 2vlgA
2vliB 2vlqA 2vm9A 2vngA 2vokA 2vpaA 2vpbB 2vq2A 2vqcA 2vqgH
2vqpA 2vrsB 2vs0B 2vs7A 2vswB 2vsyA 2vtwE 2vunC 2vv6C 2vveA
2vvfC 2vvmB 2vvwA 2vw8A 2vwaB 2vwsA 2vx8B 2vxgB 2vxnA 2vxpA
2vxtI 2vy1A 2vy8A 2vyoA 2vzcA 2vzpA 2vzyD 2w07B 2w0gA 2w0iA
2w18A 2w1jB 2w1rA 2w1vB 2w2gB 2w2nE 2w2rA 2w2uB 2w31A 2w39A
2w3gB 2w3pB 2w3qA 2w3xF 2w3yA 2w40D 2w42B 2w4sD 2w50A 2w56A
2w5eD 2w5qA 2w6aA 2w7aB 2w7vA 2w7yA 2w8tA 2w8xB 2w91A 2wadB
2wagA 2waoA 2waxD 2wb0X 2wb6A 2wbfX 2wbmB 2wbnA 2wcrB 2wcwB
2wdcA 2wdqH 2wdsA 2we5B 2we8B 2wefA 2wf7A 2wfoA 2wfpA 2wfwA
2wg7A 2wh6A 2wi8A 2wiuB 2wj5A 2wj9B 2wk1A 2wkbD 2wkdA 2wl8B
2wlvA 2wm3A 2wmyD 2wnfA 2wnkA 2wnpF 2wolA 2woyA 2wp7A 2wpvF
2wpxB 2wq4A 2wqfA 2wqkB 2wshA 2wswA 2wteB 2wtgA 2wtmD 2wtpA
2wujB 2wuqB 2wvbB 2wviA 2wvqA 2ww5A 2ww6C 2wweA 2wwxB 2wx3A
2wy3B 2wy4A 2wy8Q 2wyaC 2wzbA 2wzkA 2wzoA 2x0dA 2x0qA 2x27X
2x29A 2x2sA 2x2uA 2x32A 2x3jA 2x3lA 2x3mA 2x46A 2x49A 2x4dA
2x4jA 2x4lA 2x5cB 2x5fA 2x5hB 2x5nA 2x5pA 2x5qB 2x5rA 2x5xA
2x5yA 2x65B 2x6rB 2x78A 2x7qA 2x8tB 2x8xX 2x9kA 2x9oA 2x9qB
2x9zA 2xbgA 2xc8B 2xcbB 2xcjB 2xdjC 2xdoD 2xdpA 2xedC 2xepB
2xesA 2xetB 2xeuA 2xevB 2xf7B 2xfgB 2xfnB 2xfrA 2xfvA 2xg5B
2xgrA 2xhaB 2xhfB 2xhgA 2xhiA 2xi7C 2xioA 2xj4A 2xjpA 2xkiA
2xlgA 2xlkB 2xm5A 2xmjB 2xmoA 2xn6A 2xn6B 2xnqA 2xocB 2xodA
2xolB 2xomA 2xovA 2xp1A 2xppA 2xppB 2xpwA 2xqxA 2xrwA 2xryA
2xsaA 2xsdC 2xseA 2xt2A 2xtcA 2xtpA 2xtsA 2xtsD 2xttA 2xtyA
2xu0A 2xu3A 2xu8B 2xusA 2xvcA 2xvmA 2xvoD 2xvsA 2xvyA 2xw6C
2xwsA 2xwtC 2xwvA 2xxpA 2xyiA 2xz2A 2xz9A 2xzeB 2xziB 2y0oA
2y1bA 2y2mA 2y3mA 2y4yC 2y53A 2y5pD 2y6xA 2y71A 2y78A 2y7bA
2y7eB 2y7pA 2y8dA 2y8gB 2y8nB 2y8uB 2y9uA 2yalB 2yb1A 2ybfB
2ybyA 2yc3A 2ycdA 2ychA 2yeqB 2yevF 2yf2B 2yf4C 2yfaA 2yfuA
2yfvB 2yfvC 2yg2A 2yg9B 2yggA 2ygkA 2yh5A 2yh6C 2yh9C 2yhaA
2yhcA 2yhgA 2yhoG 2yijB 2yilE 2yimB 2yizC 2yj3A 2yjgB 2yjlB
2yjpB 2yk4A 2ykfA 2yleA 2yleB 2ylnA 2ymkA 2ymoA 2ymvA 2yn0A
2yn5A 2ynaA 2ynqC 2yopC 2yqyB 2yqzB 2yskA 2yv4A 2yv5A 2yveB
2yviA 2yvqA 2yvsB 2yvtA 2ywvB 2yx0A 2yxnA 2yxoA 2yxyA 2yy3B
2yy8B 2yykA 2yysB 2yyyB 2yziB 2yzjC 2yzsB 2yztA 2yzyA 2z08A
2z0bE 2z0dA 2z0jF 2z0rL 2z0xA 2z14A 2z2nA 2z30B 2z3jB 2z3qC
2z3xC 2z51A 2z5bA 2z5bB 2z5eA 2z5wA 2z69B 2z6oA 2z6rA 2z72A
2z7bA 2z84A 2z86A 2z8fB 2z98A 2z9wB 2za4D 2zayA 2zb4A 2zblA
2zcaA 2zcmA 2zcxA 2zd7A 2zdpB 2zdsE 2ze3A 2ze7A 2zejB 2zexB
2zfdA 2zfdB 2zfuB 2zfyA 2zg6B 2zgyB 2zhjA 2zk9X 2zktA 2znlB
2znrA 2zogA 2zosB 2zouA 2zpaB 2zptX 2zq5A 2zqmA 2zqoA 2zrrA
2zs0C 2zsiB 2zsjC 2zt5A 2zu9A 2zuxB 2zvyB 2zw2A 2zw5B 2zwaB
2zwsA 2zxeB 2zxeG 2zxrA 2zy4D 2zyrB 2zzjA 3a02A 3a07A 3a09A

3a0sA 3a16C 3a1fA 3a1gA 3a1gB 3a1jA 3a1jB 3a1jC 3a1yF 3a2eA
3a2zA 3a35A 3a4cA 3a4mA 3a4rB 3a57A 3a5fA 3a5pA 3a6rB 3a72A
3a8gA 3a8rB 3a98B 3a99A 3a9iA 3a9jC 3a9lB 3a9sA 3aa0A 3aa0B
3aafB 3abdB 3abdY 3abhA 3abiB 3abqA 3abqD 3achA 3acxA 3adrB
3adyA 3aehB 3aeiA 3afoA 3ag7A 3agcA 3agnA 3agyB 3ah9E 3ahnB
3aiaA 3aihB 3aj1A 3aj4A 3aj7A 3ajfD 3ajvA 3ajwA 3ak8E 3akbA
3akeA 3akjB 3al2A 3aljA 3alrC 3amiA 3amrA 3anoB 3anpB 3aofB
3aonA 3aotA 3aowC 3apaA 3apsA 3aq2B 3aqbA 3aqeD 3aqtA 3atsA
3au4B 3awuA 3awuB 3axbA 3axgF 3axjA 3axjB 3ay5A 3ayhA 3ayhB
3ayvD 3azdA 3azoB 3b0dB 3b0dW 3b0fB 3b0gA 3b0tA 3b0xA 3b21A
3b33A 3b40A 3b42B 3b4qB 3b4uB 3b5eA 3b5mC 3b5oA 3b64A 3b6eA
3b6hA 3b77C 3b79A 3b7cA 3b7fA 3b7hA 3b81A 3b85B 3b8fB 3b8lC
3b8xA 3b9oB 3b9tB 3b9wA 3ba3A 3bamA 3basA 3bb9F 3bbjA 3bbyA
3bbzB 3bc1F 3bc9A 3bcyA 3bczC 3bd1B 3bdiA 3bdvB 3be3A 3be6B
3bemB 3besR 3bexA 3beyE 3bf5B 3bf7A 3bfmA 3bfoB 3bg2A 3bgeA
3bguA 3bgyB 3bh0A 3bh7B 3bhdB 3bhnA 3bhoA 3bhpC 3bhqB 3bhwA
3biyA 3bj6B 3bjaA 3bjdC 3bjeB 3bjnA 3bjoA 3bjqH 3bkhA 3bkwB
3bkxB 3bl4A 3bl9B 3blnA 3blzI 3bm1B 3bm3B 3bm7A 3bmaC 3bmvA
3bmzB 3bn0A 3bn7A 3bo6A 3bodA 3boeA 3bonA 3boqB 3bosB 3bowC
3bpjB 3bpkB 3bpqA 3bpqB 3bptA 3bpvA 3bq3A 3bq9A 3bqkA 3bqoA
3bqpB 3bqwA 3bqxA 3bqzB 3brcB 3brdD 3brkX 3brsB 3bruB 3brvC
3bs4A 3bs6B 3bs7B 3bsoA 3bt3B 3bt4A 3bt5A 3btpA 3btpB 3butA
3buuA 3bv8A 3bvfF 3bw6A 3bwhA 3bwsB 3bwuD 3bwvB 3bwwA 3bwxA
3bwzA 3by8A 3by9B 3bypA 3byqB 3bywB 3c0tA 3c0wA 3c18C 3c19A
3c1aB 3c1qB 3c1yB 3c24A 3c26A 3c2bB 3c2eA 3c2gA 3c2qB 3c2uC
3c37A 3c3dC 3c3pA 3c3vA 3c3yB 3c4aA 3c4nB 3c4rD 3c57A 3c5eA
3c5nB 3c5vA 3c5xC 3c6aA 3c6fC 3c6kB 3c70A 3c7aA 3c7mA 3c7xA
3c85C 3c8dB 3c8iA 3c8lB 3c8vC 3c8yA 3c8zB 3c96A 3c9aA 3c9fA
3c9hA 3c9pA 3c9uA 3ca8A 3caiA 3canA 3cawA 3cbnA 3cbwA 3cc1A
3cc8A 3ccdB 3ccfB 3ccyA 3cddE 3cdlA 3ce8A 3ce9D 3cecA 3ceuB
3cewD 3cexA 3cfuB 3cg6A 3cggB 3cgxA 3ch4B 3chhA 3chjA 3chmA
3ci3A 3ci6B 3ci9A 3cijB 3cimA 3cinA 3citA 3cjdB 3cjeA 3cjiD
3cjjA 3cjlB 3cjmA 3cjpB 3cjsA 3cjsB 3cjyA 3ck1B 3ck6E 3ckjA
3ckkA 3ckmA 3cl5A 3claA 3clmA 3cloA 3clqB 3clwB 3cm3A 3cmbD
3cmnA 3cneD 3cniA 3cnrA 3cnuA 3cnyB 3co5B 3colB 3coqA 3covA
3cp0A 3cp7B 3cpxC 3cq1A 3cqbA 3cqjA 3cqnB 3cr3A 3craA 3crrA
3crvA 3cryA 3cs5A 3csvA 3csxB 3ct5A 3ct6B 3ct8A 3ct9A 3ctpA
3ctvA 3cu2B 3cu3A 3cu9A 3cuoA 3cuzA 3cvjC 3cvoB 3cvzC 3cwcB
3cwfB 3cwrB 3cx2A 3cxgA 3cymA 3cypD 3cz1B 3czqB 3czxB 3d00A
3d01G 3d03C 3d06A 3d0fB 3d0jA 3d0kA 3d19A 3d1bC 3d1cA 3d1lB
3d1pA 3d1rA 3d2lD 3d2qD 3d2uA 3d30A 3d32B 3d33A 3d34B 3d37A
3d3bA 3d3bJ 3d3kC 3d3mB 3d3oA 3d3rA 3d3sC 3d3yA 3d40A 3d4eA
3d4uB 3d59A 3d6wB 3d79A 3d7aA 3d7iB 3d7jB 3d7lE 3d7rB 3d8pA
3d9rC 3d9xB 3da5A 3da8B 3dadB 3dalB 3daoA 3db2B 3db7A 3dboA

3dbyQ 3dc7C 3dcfB 3dclE 3dcmX 3dczA 3dd7A 3ddcB 3dddA 3ddeB
3ddtC 3deeA 3defA 3deoA 3dewA 3df6B 3df7A 3df8A 3dffA 3dfgA
3dfuB 3dfzA 3dgpA 3dgpB 3dhaA 3dhuD 3dhxB 3di5A 3dj1A 3djeB
3djlA 3dk9A 3dkaA 3dkmA 3dkqC 3dkrA 3dkzA 3dlcA 3dliC 3dlqI
3dlqR 3dluD 3dm8A 3dmcA 3dmeA 3dmgA 3dmlA 3dmnA 3dmyA 3dn7A
3dnhB 3dnjA 3dnpA 3dnsA 3dnxA 3do8A 3douA 3dp7B 3dpjA 3dqpA
3dr2B 3dr5A 3draB 3drfA 3ds4B 3ds8A 3dsbB 3dsdA 3dskA 3dsoA
3dssA 3dt5A 3dtdI 3dtnA 3dtyA 3dupA 3duwA 3dwoX 3dx5A 3dxeA
3dxiB 3dxlA 3dxpA 3dxrA 3dxrB 3dxyA 3dydB 3dyjA 3dytA 3dz1A
3dzaD 3e03A 3e05B 3e0eA 3e0hA 3e0rC 3e0sB 3e0xB 3e0zD 3e11A
3e15B 3e19C 3e1eC 3e1rB 3e35A 3e38B 3e3vA 3e4bD 3e4gA 3e4vA
3e4wA 3e56A 3e57A 3e58B 3e59A 3e5xA 3e6mH 3e7kD 3e7qB 3e8lC
3e8mD 3e8oB 3e8sA 3e8tA 3e96B 3e97A 3e98A 3e99A 3e9fA 3e9uA
3e9vA 3ea0A 3ea6A 3eafA 3eb8A 3ebbD 3ebeC 3ebtA 3ebyA 3ec3A
3ec6A 3ecfA 3edfB 3edvB 3edyA 3eeaA 3eehA 3eeqA 3eerA 3ef8A
3efgA 3egaA 3egeA 3ehcA 3ehdB 3ehgA 3ei3B 3einA 3eipB 3ej9B
3ej9C 3ejfA 3ejkA 3ejvA 3ek3A 3ekiA 3eleA 3elfA 3elsA 3emfB
3emiA 3emxB 3en0A 3en8A 3enuA 3eo4B 3eo6B 3eo7A 3eofA 3eoiB
3eojA 3ep1A 3eqxB 3er6B 3er9B 3ermE 3ervA 3erwD 3es1A 3eshC
3esiA 3eskA 3eslB 3essA 3etoB 3etvA 3etzA 3eulB 3eunA 3eurA
3eusA 3evnA 3evyA 3evzA 3ewgA 3exmA 3exnA 3exzE 3ey5A 3eyeA
3eyiB 3eytB 3eyyB 3ez0D 3ez2A 3eziB 3f08A 3f0dF 3f13B 3f14A
3f1iH 3f1lB 3f1pB 3f1zJ 3f2eA 3f2iB 3f2zA 3f3bA 3f40A 3f42A
3f43A 3f4aA 3f4mA 3f4sA 3f5bA 3f5hB 3f5oF 3f5rA 3f62A 3f67A
3f6cA 3f6gB 3f6vA 3f6yA 3f75P 3f7eB 3f7sA 3f7wA 3f8bA 3f8dA
3f8tA 3f8xD 3f95A 3f9uA 3facD 3fakA 3fanA 3fauB 3favB 3fayA
3fb9B 3fbgB 3fblA 3fbuB 3fcmB 3fd4B 3fdiB 3fdjA 3fdqA 3fdsC
3fegA 3fetD 3feuA 3ff2A 3ff4A 3fflD 3ffvB 3fg8B 3fg9C 3fghA
3fgrA 3fgvA 3fh3A 3fhdA 3fhkD 3fhlB 3fi9B 3fidA 3fj1A 3fjuB
3fjvA 3fk4B 3fk8A 3fkcA 3fkeB 3fkqA 3fl2A 3flaA 3fldB 3flpA
3fm0A 3fm2A 3fmcD 3fnbA 3fndA 3fniA 3fnrA 3fo8D 3fokJ 3fotA
3fouA 3fpcD 3fpfB 3fpnA 3fprA 3fqmA 3frhA 3frmD 3frnA 3frqB
3fryA 3fs3A 3fs8B 3fsaA 3fsoA 3fssA 3ft1A 3ft7A 3ftdA 3fuyA
3fvvA 3fvzA 3fwbB 3fwkA 3fwyA 3fwzB 3fx7B 3fxdB 3fymA 3fz5C
3fzeA 3fzqB 3fzxA 3fzyB 3g02B 3g0kA 3g0mA 3g13B 3g16A 3g1jA
3g21A 3g27A 3g2bA 3g2eA 3g36C 3g3sB 3g3tA 3g3zB 3g40A 3g46A
3g48A 3g4nB 3g5bA 3g5jB 3g5sA 3g67A 3g74D 3g7qA 3g7rB 3g85A
3g8yA 3g8zA 3g91A 3g98A 3g9mB 3g9yA 3ga3A 3ga4A 3ga7A 3gaeA
3gb5A 3gbvA 3gbyA 3gd0A 3gd6A 3gdhC 3ge3A 3ge3B 3ge3C 3ge3E
3gfaA 3gffB 3gfsL 3gg7A 3ggdA 3ggnB 3ggyA 3ghaA 3ghfA 3giaA
3gj0A 3gjoH 3gjyA 3gk6A 3gk7A 3gkeC 3gkjA 3glaA 3glcC 3gldB
3glvA 3gm5A 3gmeD 3gmfA 3gmiA 3gmxB 3gn3B 3gnaA 3gneB 3gnfB
3gnjD 3gnzP 3go2A 3gocB 3goeA 3gonA 3gozA 3gp6A 3gpgB 3gpiA
3gpvB 3gq8A 3gqhB 3gqqE 3gqvA 3gqxA 3gr3A 3gr5A 3graA 3grcD

3grdB 3greA 3grlA 3grzB 3gs2B 3gs2C 3gseA 3gtzC 3gudB 3guyC
3gv0A 3gwbB 3gwcF 3gwiA 3gwjC 3gwlB 3gwnA 3gwrA 3gwyA 3gx8A
3gxhA 3gxqB 3gxvB 3gybB 3gygD 3gykD 3gyyD 3gzrB 3h05A 3h0dB
3h0nA 3h0oA 3h0uB 3h16D 3h1dA 3h1nA 3h1tA 3h20A 3h2gA 3h2sB
3h2yA 3h49A 3h4cA 3h4lA 3h4tA 3h51B 3h5jA 3h5lA 3h6eB 3h6jA
3h6pA 3h6qA 3h74A 3h75A 3h79A 3h7aC 3h7hA 3h7hB 3h7iA 3h7jB
3h86G 3h8dC 3h8gC 3h8kB 3h8tB 3h8uA 3h8vB 3h9cA 3h9wA 3ha2A
3ha9A 3hbcA 3hbmA 3hc1A 3hc7A 3hdjB 3hdxA 3he4H 3he5B 3hf5B
3hfeB 3hfiA 3hfwA 3hg9A 3hgtA 3hguA 3hhfB 3hi0B 3hi2D 3hieB
3himB 3hj7A 3hjhA 3hkaB 3hklB 3hkmB 3hkvB 3hkwC 3hl1A 3hl6B
3hlsD 3hlxD 3hmsA 3hmzA 3hn3B 3hn5B 3hnoC 3hnyM 3ho6A 3ho7B
3hoiA 3holA 3hpcX 3hpyB 3hq1B 3hqxA 3hr0B 3hr6A 3hrdG 3hrgA
3hroA 3hrpA 3hrqB 3hrzA 3hrzB 3hs3B 3hshD 3hsrB 3hsyA 3htkA
3htkB 3htkC 3htrB 3htuC 3htuD 3htyK 3hugF 3huhC 3hutA 3huuB
3hv2B 3hvaB 3hwuA 3hx3A 3hx8B 3hyiA 3hynA 3hz6A 3hz8A 3hzpA
3i00A 3i09B 3i0wA 3i0yC 3i1aB 3i2kA 3i2nA 3i2vA 3i33A 3i38A
3i3vB 3i45A 3i4gA 3i4zB 3i57B 3i5qB 3i6eA 3i83B 3i8bA 3i94A
3i96C 3i9fB 3i9yA 3ia1B 3iagC 3iarA 3ib7A 3ibhA 3ibwB 3ic9B
3icuA 3id1A 3idfA 3iduB 3idwA 3ie4B 3ie7A 3iekA 3ieyB 3iezA
3if4B 3ifeA 3ifnP 3ifrB 3ifuA 3ig9D 3ighX 3igmB 3ih6E 3ihmB
3ii2A 3iiiA 3iisM 3ij6D 3ijdB 3ijlA 3ikbB 3ikwA 3ilsA 3ilvA
3ilwB 3im1A 3im3A 3im6A 3imkA 3imoD 3in6B 3io1A 3io3A 3io5B
3ip0A 3ip4B 3ip4C 3ipfB 3ipjA 3iprE 3iraA 3irbA 3irsA 3isaA
3it3A 3it4A 3it5B 3iteA 3itwB 3iu0A 3iukB 3iupB 3iuuA 3iuwB
3iv1D 3ivpD 3ivvA 3ix0D 3ix3A 3ixsL 3jq1A 3jqyA 3jrnA 3jrtA
3js6A 3js8A 3jsjC 3jsyB 3jszA 3jtnA 3jtwA 3jtzA 3ju4A 3judA
3jumA 3jv1A 3jv9B 3jwgA 3jx9B 3jxoB 3jy6D 3jybB 3jygD 3jyzA
3jz0B 3jzyA 3k01A 3k05B 3k0yA 3k0zA 3k1hA 3k1rB 3k1tA 3k2nB
3k2oA 3k32E 3k3dA 3k3fA 3k3vA 3k4iA 3k50A 3k5jA 3k63A 3k67B
3k6gC 3k6gD 3k6mC 3k6rA 3k6tC 3k6yA 3k7cB 3k7iB 3k7pA 3k8gB
3k8uA 3k93A 3k94A 3ka5B 3ka7A 3kaeC 3kanB 3kb4B 3kb9A 3kbqA
3kbrA 3kc2A 3kcuE 3kd6A 3kdwA 3ke2A 3ke7B 3keaB 3keoB 3kewA
3keyA 3kf6A 3kf6B 3kf8C 3kf8D 3kffA 3kg0A 3kgkA 3kgwA 3kgyB
3kgzB 3kh0B 3kh1B 3kh8B 3kheA 3khqA 3kizA 3kjhA 3kk4C 3kk7B
3kkcD 3kkgA 3kkiB 3kklA 3kkzA 3kljA 3klqB 3kluA 3km3B 3km5A
3kmaA 3kmuA 3kndB 3knzD 3kojB 3kp1F 3kp7A 3kp8A 3kpeA 3kq5A
3kqnA 3kraC 3ks6A 3ksnA 3ksuA 3ksxA 3ktaB 3ktaC 3ktcB 3ku3B
3kuvB 3kv1A 3kvhA 3kvnA 3kvpA 3kw2B 3kweA 3kwrA 3kxrA 3kxtA
3kxwA 3kyfA 3kyjA 3kyzA 3kz3B 3kz5E 3kzhB 3kzpA 3kzqE 3kzxA
3l0qA 3l1nA 3l1wD 3l22A 3l2bB 3l2hB 3l32A 3l39A 3l46A 3l49D
3l4cA 3l4eA 3l4nA 3l4qD 3l51A 3l51B 3l60A 3l6bA 3l6tB 3l7hD
3l81A 3l82B 3l8dA 3l8eA 3l9bA 3la7A 3laeA 3lagA 3laxA 3lazB
3lb2B 3lc0A 3lccA 3ld1A 3ld7A 3ldtA 3lduA 3ldvB 3le4A 3letB
3lfjB 3lfkB 3lfuA 3lgbB 3lgdA 3lgeF 3lgiC 3lhcA 3lheA 3lhlA

3lhnA 3lhqB 3lhsA 3lidB 3ljbB 3ljxA 3lkbB 3lkdB 3lkeC 3lkmA
3ll3B 3ll5D 3llcA 3llkA 3lloA 3llpA 3lluA 3llvA 3lm5A 3lmzA
3lo8A 3lodA 3logC 3lopA 3loqA 3louB 3lp5A 3lpzA 3lqbA 3lqcA
3lqnA 3lqvB 3lrtB 3ls9A 3lsjA 3lt7F 3ltiA 3luaA 3luiA 3lumA
3lvyD 3lw3B 3lw7B 3lwaA 3lwtX 3lx3A 3lxrF 3ly7A 3lyeA 3lywA
3lyxB 3lzkD 3m03A 3m0fB 3m0mD 3m1rB 3m1tA 3m2tA 3m31A 3m33A
3m3hA 3m3pA 3m4iA 3m4wD 3m5bA 3m5qA 3m66A 3m6nC 3m6zB 3m73A
3m7kA 3m7nA 3m7oA 3m89A 3m8jA 3m91A 3m91B 3m92B 3m9lA 3m9qA
3mabB 3madA 3malB 3maoA 3mayH 3mb2D 3mb5A 3mbkA 3mbrX 3mcbB
3mceA 3mcwB 3mcxA 3md1A 3md7A 3mdmA 3mduA 3me0B 3me7A 3meaA
3memA 3merB 3mfbB 3mfxC 3mgdB 3mggB 3mhxB 3mi0I 3milA 3mj6A
3mjhB 3mjoB 3mk4A 3mkcA 3mkhD 3mkoA 3mkqF 3mlnA 3mmgA 3mmhB
3mmpG 3mmyG 3mn2A 3mn5S 3moiA 3mpcB 3mpkA 3mq0A 3mqdA 3mqqB
3mqzA 3mr0A 3mseB 3mstA 3mswA 3msxB 3mszA 3mt0A 3mtrB 3mu7A
3mujB 3mvcA 3mvsA 3mvuA 3mw6F 3mw8A 3mwxA 3mwzA 3mx7A 3mxmB
3mxnA 3mxnB 3mxoB 3mxzA 3my2A 3mydA 3n01A 3n08B 3n0aA 3n0kA
3n0rA 3n0uC 3n10A 3n17A 3n1eA 3n2wC 3n3mA 3n3uA 3n3yB 3n40P
3n4jA 3n54B 3n5bB 3n6tA 3n6xA 3n72B 3n77A 3n79A 3n7xA 3n9yA
3na2D 3na7A 3napA 3nb2B 3nbcA 3nceA 3nctC 3nd1B 3ndhB 3ndqA
3ndzF 3necB 3nehB 3nfiE 3nftA 3ng7X 3ngwA 3ni0B 3ni8A 3nj2B
3njdA 3njeA 3njnC 3nk4A 3nk6B 3nkeC 3nkgB 3nklA 3nksA 3nkuB
3nkzA 3nl9A 3nlcA 3nnfA 3nngB 3no0A 3no2A 3no3A 3no4A 3no6A
3no7B 3nohA 3nojA 3noqA 3npdA 3nphB 3nqbB 3nqnB 3nqoB 3nqzA
3nr5A 3nraB 3nreD 3nrhB 3nrlB 3nrsA 3nrwA 3nrxA 3ns6B 3nswF
3nsxB 3ntkA 3ntvA 3nufA 3nuqA 3nutA 3nvoB 3nvsA 3nvwB 3nvxA
3nw4A 3nycA 3nzlA 3nznA 3nztA 3o0gE 3o0lA 3o0qB 3o0yB 3o10C
3o12A 3o14B 3o22A 3o2iB 3o2rA 3o2tA 3o3mA 3o3mB 3o48A 3o4hD
3o4pA 3o53A 3o59X 3o6qA 3o6qD 3o6zB 3o71B 3o7bA 3o83B 3o8mA
3o94D 3o9zA 3oa5B 3oajA 3oamD 3obeA 3obfB 3obhB 3oblA 3obqA
3oc8A 3ocjA 3ocuA 3od1A 3od8H 3odtB 3odvA 3oe3A 3oepA 3of4B
3ofkB 3og6A 3og9B 3oghA 3ognA 3oh3A 3oheB 3ohgA 3ohsX 3oiqA
3oiqB 3oisD 3oiyA 3oizA 3oj0A 3okgA 3okpA 3okqA 3okxB 3oloB
3om0A 3omyA 3on9A 3ondB 3onhA 3onjA 3onmB 3onqB 3oo8A 3ooqJ
3ooxA 3ooyB 3op6B 3op9A 3opcB 3oq4A 3oqqA 3or1D 3or1E 3orhD
3orjA 3orkA 3oruA 3oryA 3oseA 3ostA 3osvA 3ot2A 3otdA 3otiB
3otnA 3ou2A 3oueA 3oulA 3ov5A 3ov8A 3ov9A 3ovkC 3ow8A 3owcB
3owrC 3oxhA 3oy2B 3oyoB 3oyvA 3oyzA 3oz2A 3ozpA 3p02A 3p06A
3p0bA 3p0fA 3p0kA 3p0yA 3p1vA 3p1xA 3p24A 3p2eB 3p3cA 3p4hA
3p51A 3p6lA 3p6zI 3p7iA 3p8aA 3p8cF 3p9aA 3p9vA 3p9zA 3pasB
3pb6X 3pbtA 3pcvA 3pd7A 3pdgA 3pdyB 3pe5A 3pe6A 3pe7A 3pesB
3petA 3pf0A 3pf2A 3pf7A 3pfeA 3pfgA 3pfoB 3pfyA 3pg0A 3pg7B
3pguA 3phhA 3phsA 3pijB 3piuA 3pivA 3piwA 3pjpA 3pjvD 3pkoA
3pkzD 3pl0B 3pl8A 3plnA 3pluA 3plwA 3pm2A 3pmoA 3pmsA 3pnnA
3pnxF 3pohA 3pojB 3popC 3powA 3pp2B 3pp5A 3ppbB 3pplA 3ppmB

3proD 3ps0D 3pshA 3psmA 3pt1A 3pt5A 3ptyA 3pu5A 3pu9B 3pucA
3pveB 3pvhA 3pviB 3pvtA 3pvvA 3pw3D 3py9A 3pycA 3pywA 3q18B
3q1cA 3q1nA 3q1tB 3q1xA 3q20B 3q2iA 3q3eA 3q3jB 3q46A 3q4oA
3q63D 3q64A 3q6aD 3q6bA 3q6zA 3q7mA 3q7rB 3q87A 3q87B 3q8dB
3q9dB 3q9lB 3qaoA 3qb8B 3qbmB 3qc0A 3qc1A 3qc7A 3qd7X 3qdlD
3qf2A 3qf7D 3qfeA 3qflA 3qfmA 3qftA 3qguB 3qh6A 3qh9A 3qhbB
3qheB 3qhoC 3qhpB 3qhqA 3qhyB 3qi7B 3qitA 3qj4B 3qksC 3qnmA
3qnsA 3qooA 3qp4A 3qqaA 3qqmH 3qr7B 3qraA 3qrlA 3qs2A 3qsjA
3qslA 3qszA 3qthB 3qu3C 3qu5A 3qufA 3qv0A 3qvpA 3qvsA 3qw9B
3qweA 3qwgB 3qwlA 3qwnC 3qwuB 3qx1A 3qxcA 3qxyB 3qy7A 3qy9A
3qyeB 3qzbA 3qzmA 3qzrA 3qzxA 3r0aA 3r0nA 3r15A 3r1kA 3r1xB
3r24A 3r2pA 3r2uD 3r3cA 3r3pB 3r41A 3r44A 3r4iF 3r4zA 3r5tA
3r62B 3r6aA 3r72A 3r75A 3r7aB 3r87A 3r8jB 3r90F 3r9fB 3r9mA
3rbnA 3rbsA 3rbyA 3rc1A 3rd5A 3renB 3retA 3rf0B 3rf3B 3rfeB
3rfiA 3rftC 3rfyA 3rgaA 3rgcB 3rgoA 3rh0B 3rhtD 3rhzA 3ripA
3rj2X 3rjoA 3rjuA 3rkcB 3rkgA 3rl5A 3rlfF 3rlgA 3rlkA 3rloA
3rlsA 3rm3A 3rmhA 3rmqA 3rmuD 3rnlA 3rnqA 3rnrB 3rnvA 3ro2B
3ro3A 3robB 3rofA 3rotB 3rpcB 3rpeA 3rpfB 3rppC 3rpzA 3rq1C
3rq4A 3rq9B 3rqaB 3rriB 3rs1B 3rstG 3rt2A 3rt3C 3rufB 3rvcA
3rwnB 3rx6A 3rx9A 3ry0A 3ry4A 3rzaA 3rznA 3s0aA 3s2jA 3s2rB
3s32A 3s35X 3s44A 3s4eA 3s4lA 3s4yA 3s5bB 3s5jA 3s5wA 3s64A
3s6eB 3s6fA 3s6pB 3s6pE 3s84B 3s8gA 3s8gB 3s8gC 3s8iA 3s8mA
3s8sA 3s98A 3s99A 3s9dC 3s9dD 3s9xA 3saoA 3sb4A 3sbmA 3sbtB
3sc7X 3sd6A 3sd7A 3sdbA 3sfvB 3sg0A 3sggA 3sgwA 3shgA 3shgB
3shpB 3shqA 3shsA 3sibA 3sigA 3sj5B 3sjhB 3sk2B 3sk9A 3skqA
3skvA 3sl9F 3slrA 3sm4B 3smvA 3snyA 3so6A 3sojA 3sokA 3sp7A
3sq7D 3sqlA 3sqzA 3sreA 3sriB 3ss7X 3ssbD 3ssoC 3stoA 3su6A
3suiB 3sukA 3sumD 3suuA 3swyA 3sx6A 3sxmB 3sxuA 3sxuB 3sy1A
3sz3A 3szaB 3szvA 3szyA 3t0yD 3t1oA 3t2cA 3t33A 3t3lA 3t47B
3t4lB 3t4rA 3t5nA 3t5xA 3t6aA 3t7aA 3t7dB 3t7hB 3t7lA 3t7zA
3t8kB 3t92A 3t9oB 3t9wA 3t9yA 3tacB 3tahB 3tbdA 3tbnA 3tboA
3tc3B 3tcvA 3tdgA 3tdnB 3tdsE 3te8B 3teeA 3tekB 3teqC 3teuA
3tfgB 3tfjB 3tg0D 3tg2A 3tg9B 3thgA 3thiA 3tj5B 3tj8B 3tjmA
3tjrA 3tjyA 3tl4X 3tlqB 3tm4A 3tm8A 3tmgB 3to3A 3towA 3tpdA
3tq2A 3ts3B 3ts9A 3tsaA 3tt9A 3ttcA 3ttgA 3tufA 3tufB 3tuoA
3tutA 3tv0B 3tvaB 3tvjA 3tvqA 3tvtB 3tw0D 3tw8C 3tx3A 3txsD
3txyA 3ty1A 3tysA 3tzyA 3u07A 3u0cB 3u0rA 3u0vA 3u12A 3u1cB
3u1uB 3u26A 3u28B 3u28C 3u2aA 3u2gA 3u2rA 3u2uA 3u4gA 3u4tA
3u4vB 3u4zB 3u50C 3u52B 3u5vA 3u64A 3u65B 3u7qB 3u7qC 3u7zA
3u8vB 3u99A 3u9gA 3u9jA 3u9rB 3u9wA 3uafA 3uanA 3ub1B 3ub2A
3ub6A 3uc9A 3ucsB 3ud1C 3uebD 3uenA 3uf6B 3ufbA 3ufcX 3ufeA
3uffB 3ug9A 3ugfB 3uguA 3uh8A 3ui4A 3uidA 3uitA 3ujcA 3ukwC
3ukyC 3ulbA 3uljB 3ultA 3umhA 3umoA 3un6A 3uo2B 3uoiU 3uplB
3upsA 3uqcD 3ur8B 3urgA 3urrA 3us3A 3ut4B 3uueA 3uulA 3uuwD

3uv0A 3uv1A 3uv5A 3uw3B 3uwpA 3ux2A 3uxfA 3uxjB 3v0dB 3v0rA
3v1aA 3v1vA 3v32A 3v3lA 3v3tA 3v42B 3v46A 3v4cB 3v4gA 3v4kA
3v4yD 3v5cC 3v5rB 3v67B 3v68A 3v69A 3v6gA 3v6iB 3v6oA 3v7bB
3v7nA 3v93D 3v96A 3v9oA 3v9rD 3va9A 3vb0C 3vbcA 3vbjA 3vc1L
3vc8B 3vcaA 3vcxA 3vdjA 3vejB 3venA 3vepG 3vfzA 3vg8B 3vgiA
3vglA 3vgpA 3vgzC 3vhjA 3vhlA 3vi6A 3viiA 3viqA 3vivB 3vj9A
3vk5A 3vk6A 3vkwA 3vl1B 3vldA 3vmkA 3vmnA 3vn0C 3vn5A 3volA
3vopB 3voqB 3vorA 3votB 3vp5A 3vp9A 3vpbA 3vpbF 3vpzA 3vqjA
3vqtC 3vr0A 3vrhA 3vs8G 3vsjB 3vsnA 3vsvB 3vtxA 3vu9A 3vu9B
3vubA 3vupB 3vuqC 3vvvA 3vvyD 3vwaA 3vwnX 3vx0A 3vxcB 3vy8X
3vypB 3vywD 3vyxA 3vz9B 3vz9D 3vzaF 3vzhA 3vzxA 3w06A 3w07A
3w0eA 3w0fA 3w0kB 3w0oA 3w0tD 3w15A 3w15B 3w1eA 3w1oB 3w20B
3w36B 3w3wB 3w42A 3w4sA 3w4tA 3w54C 3w56A 3w5sB 3w6bB 3w6sA
3w6wB 3w7yB 3w8iB 3wa1A 3warA 3wasB 3wcqA 3wdnA 3we0A 3we9A
3wecA 3wfdB 3wfdC 3wg3B 3wgxA 3wh1A 3wh2A 3whjA 3wi3B 3wisA
3wiwA 3wjtA 3wkgA 3wkxA 3wkyB 3wliA 3wmiA 3wmiB 3wmtB 3wmvB
3wn7M 3wndA 3wnzA 3wo6A 3woeC 3woeD 3wozB 3wqbB 3wqcA 3wqmA
3wrbA 3wryB 3ws7A 3wsgE 3wt0C 3wtdA 3wttG 3wucA 3wupA 3wurA
3wuzA 3wv4B 3wv7C 3wvaB 3wvqA 3wvtA 3wvzB 3ww3A 3wwcA 3wwnA
3wwqL 3wwtB 3wx4A 3wx7A 3wxfA 3wydB 3wz3A 3wzsA 3x01B 3x0iA
3x0tA 3x0uB 3x0vA 3x1lB 3x1lC 3x27D 3x2mA 3x30A 3x34A 3x38B
3zbdB 3zboB 3zdbA 3zdoG 3zdsF 3zeuA 3zf8A 3zfiA 3zfpA 3zg9A
3zghA 3zgjB 3zh5A 3zh9B 3zhiA 3zi1A 3zidB 3zieD 3zigA 3zihB
3zilA 3ziuB 3ziwF 3zj0A 3zjaA 3zjxD 3zk4C 3zl8A 3zmdB 3zn3A
3zn4A 3zn6A 3znuA 3znvA 3zojA 3zoqB 3zpjA 3zpxB 3zqoQ 3zqsA
3zr8X 3zrgA 3zriA 3zrxB 3zsjA 3zssC 3zsuA 3zt9A 3zthB 3ztvA
3zucA 3zuiA 3zvsC 3zwfA 3zx3B 3zxcB 3zxkA 3zxnA 3zy7B 3zypA
3zzhC 3zzoA 3zzpA 3zzsG 3zzyA 4a02A 4a0eB 4a0pA 4a15A 4a1rA
4a20A 4a29A 4a2bA 4a2vA 4a37A 4a3pA 4a3zA 4a4jA 4a56A 4a57C
4a5nC 4a5uB 4a5zB 4a6dA 4a6hD 4a6qA 4a7uA 4a7wB 4a8jA 4a8jE
4a8jF 4a8xB 4a9vA 4abxC 4ac1X 4acfC 4aciB 4acjA 4aclA 4acvA
4admD 4adnA 4adzA 4ae2A 4ae5D 4ae7A 4aeeB 4aeqA 4aezG 4af1A
4af8A 4affA 4afkA 4aflE 4afmA 4ag6B 4agrB 4agsB 4aikA 4aivA
4ak2A 4akkA 4alzA 4amqA 4annA 4anoA 4ao6A 4apoB 4aq4A 4aqlA
4aqnB 4aqoA 4ar9A 4artA 4aruA 4as2D 4asmB 4at0A 4at7B 4ateA
4atgA 4atmA 4au1A 4avrA 4avsE 4aw7A 4awxB 4axdA 4axoB 4axvA
4ay0A 4ayaB 4ayoA 4az6A 4azsA 4b0mA 4b0zA 4b1mB 4b1yB 4b1yM
4b2fA 4b2nA 4b2oA 4b46A 4b4cA 4b4dA 4b4sA 4b4uB 4b4yA 4b5oA
4b60A 4b6gA 4b6xB 4b89A 4b8eB 4b9gB 4bb9A 4bbyB 4bc3A 4be3B
4beuA 4bfcA 4bfgA 4bfhA 4bfoA 4bg7A 4bgbA 4bgpA 4bhqB 4bhrA
4bhuD 4bi3B 4bj0A 4bjaA 4bjiA 4bjjA 4bjjB 4bjsB 4bjtB 4bjzA
4bk0B 4bk7A 4blpC 4bluA 4bmdA 4bn4A 4bndA 4boeA 4bolA 4boqA
4bouA 4bpfA 4bpsA 4bpzA 4bqhA 4bqnA 4bqqA 4bqyA 4brcB 4bspA
4bsxB 4bt7A 4btbA 4bvxA 4bvxB 4bwqB 4bwrA 4bwvA 4bx8B 4bxfA

4bxiA 4bxmA 4byzA 4bzaA 4c08A 4c0fD 4c1aD 4c1wA 4c24A 4c2fA
4c2lA 4c3sA 4c47A 4c4aA 4c5cB 4c5eH 4c5kB 4c5wA 4c6aA 4c6sA
4c7nA 4c89C 4c92B 4c92E 4c97A 4c9bB 4c9sC 4c9yA 4ca1B 4cadC
4cahB 4cayA 4cayC 4cbeA 4cbuG 4cc9B 4ccsA 4ccvA 4ccwA 4cd5A
4cd8A 4cdjB 4cdpA 4ce8C 4cekA 4cfiA 4cfsD 4cgoA 4cgrA 4cguB
4chdA 4cheA 4chiA 4chmD 4ci7B 4ci9A 4cicB 4cihC 4ciiA 4cijB
4cj0A 4cj0B 4cjdA 4ckmB 4clqA 4clqB 4cmrB 4cn0B 4cnnA 4co6C
4co8A 4cogC 4cp6A 4cpcG 4crhA 4cruA 4cruB 4cs4A 4csrA 4csrB
4ct0B 4ct3B 4cu5D 4cuaB 4cv7A 4cvbA 4cvdA 4cvoA 4cvqA 4cvrA
4cw4A 4cxfB 4cz5D 4czgA 4czxA 4czxB 4d05B 4d07B 4d0pA 4d0qA
4d2hC 4d2kB 4d4zA 4d53A 4d5bA 4d5tF 4d6gA 4d6vA 4d6zA 4d79B
4d7cB 4d7eD 4d7jA 4d8bA 4d8mA 4d9bB 4da2A 4dacB 4damD 4db5A
4dcbA 4dckA 4dd5A 4ddpA 4devC 4dgfA 4dhxA 4di9A 4didB 4dixA
4djaA 4djcB 4dk2A 4dkcB 4dkkA 4dkwC 4dlhA 4dloB 4dm5D 4dmgB
4dmiD 4dmoA 4dmvA 4dndA 4dnhA 4dnyA 4do4A 4do7A 4doiB 4dolA
4dooB 4doqD 4dovA 4dpbX 4dq9B 4dqjB 4dqzA 4driB 4dt4A 4dt5A
4dthA 4dv8A 4dvcA 4dveC 4dvkA 4dwdA 4dwrA 4dxrB 4dylA 4dynA
4dyqB 4dz1A 4dz4C 4dziA 4dzzB 4e0aB 4e0qB 4e17B 4e1pA 4e2gH
4e2uA 4e2xA 4e3eA 4e3xA 4e40A 4e45D 4e6nC 4e6sA 4e6uA 4e6wA
4e6zA 4e74A 4e9sA 4e9xC 4ea9A 4eadA 4eaeB 4ebbA 4ebjB 4ebyA
4edpA 4ee6A 4efiA 4efoA 4egcB 4egdA 4eguA 4egwB 4eh1A 4ehcA
4ehuB 4ehxA 4ei0B 4ei7B 4eicA 4eijA 4ejrB 4ekxA 4el6A 4elnA
4emnB 4emoA 4emtB 4eneA 4eo0A 4eogA 4ep4B 4eq3A 4eqbB 4eqpA
4eqqA 4eqsB 4ercA 4ernA 4errA 4eryA 4es1A 4es8B 4eskB 4esmA
4esqA 4eswA 4etnA 4etrA 4eu0A 4eu9A 4eunA 4euoA 4ev1A 4evfA
4evuB 4evwB 4evyB 4ew6A 4exjA 4exkA 4exoA 4extA 4eycA 4eysA
4eytF 4ezgA 4eziA 4f01A 4f02F 4f03A 4f06A 4f0qB 4f0wA 4f0zC
4f1jB 4f1vA 4f2fA 4f3nA 4f3vA 4f43A 4f55A 4f7gB 4f7hA 4f7uP
4f87C 4f98A 4f9cB 4fa8A 4fbcB 4fbdB 4fbjA 4fbsA 4fbwB 4fbwC
4fc3E 4fcaA 4fcgA 4fchB 4fd7D 4fdbA 4fddB 4fekB 4ff5A 4fflA
4ffuC 4fgcE 4fgmA 4fgqB 4fgwA 4fhrA 4fhrB 4fibB 4finB 4fixA
4fk9A 4flaB 4flbA 4fleA 4fp4A 4fp5G 4fppA 4fprC 4fpwB 4fqeA
4fqgB 4fqnD 4fsdA 4ftfA 4fuvA 4fvdA 4fvgA 4fwvA 4fwwA 4fxqA
4fybB 4fypA 4fyyB 4fz2B 4fz4A 4fzlB 4fzpA 4fzqE 4fzvB 4g08A
4g0aD 4g0iB 4g0xA 4g1iA 4g1oA 4g1qB 4g26A 4g29A 4g2eA 4g2sC
4g38A 4g3aA 4g3fA 4g3hB 4g3nA 4g3oA 4g3vA 4g4gA 4g4kB 4g54A
4g55A 4g6dB 4g6iB 4g6qB 4g6tA 4g6tB 4g78A 4g79A 4g7nA 4g7vS
4g7xA 4g7xB 4g8aB 4g8tA 4g92A 4g9eB 4g9qA 4g9sA 4g9sB 4ga2A
4gakA 4gb5A 4gbjB 4gbmA 4gc3A 4gdiE 4gdoE 4gehA 4geiA 4gf3A
4gf3B 4ggtA 4gi3C 4gimB 4gioB 4gipB 4giwA 4gj4D 4gjrA 4gjzA
4gkhH 4glkA 4gm6D 4gmoA 4gmqA 4gmuA 4gn5A 4gneA 4gofA 4gosA
4gouA 4gpsA 4gq1A 4gqbA 4gqmA 4gs3A 4gs5A 4gt8A 4gt9A 4gu5A
4gvbB 4gvqA 4gwbA 4gwgA 4gx8D 4gxbA 4gxbB 4gxtA 4gxwA 4gyoA
4gytA 4gzcA 4gzkA 4h0cA 4h0fB 4h14A 4h18C 4h2dA 4h3tA 4h3vA

4h3wB 4h41A 4h4dB 4h4nA 4h59A 4h5iA 4h5sB 4h6cI 4h6qC 4h6xA
4h79A 4h7lB 4h7wA 4h7yC 4h86A 4h87A 4h8eA 4h9nC 4ha7B 4hatB
4hbeB 4hbzA 4hc9A 4hceA 4hcsA 4hddA 4hdeA 4hdoA 4hdqC 4he6A
4heiA 4hfkD 4hfqB 4hfsB 4hfvA 4hguA 4hh3C 4hh5A 4hhjA 4hhrA
4hhvB 4hhxA 4hi0B 4hi6D 4hi8A 4hi8B 4hiaB 4hj1C 4hjiB 4hjzA
4hkgA 4hkhG 4hkuA 4hl4A 4hlsB 4hlyA 4hmsB 4hnlA 4hnoA 4hpmA
4hpmD 4hr1B 4hr3A 4hr6C 4hroB 4hrsA 4hrvB 4hs1A 4hs2A 4hstA
4hstB 4hsuB 4htfA 4htgA 4htmA 4htpB 4htuB 4huaA 4hvkA 4hvyA
4hwcA 4hwnA 4hwvB 4hxfB 4hy4B 4hylB 4hynB 4hz4A 4hz9A 4hz9B
4hziB 4hzoA 4i0oA 4i0wA 4i0xC 4i0xJ 4i16A 4i1fA 4i1kB 4i1lA
4i1sB 4i2oB 4i3mA 4i3yB 4i4cB 4i4oB 4i5jA 4i5tB 4i66A 4i6kA
4i6rA 4i6vC 4i6xA 4i6yA 4i71A 4i84B 4i86B 4i8iA 4i90A 4i93A
4i9xA 4iauA 4ic1K 4ic4A 4ic9A 4icvA 4id9B 4ie5A 4igbC 4igiA
4iglD 4iguB 4ihqC 4ihuB 4ihzA 4iikA 4iilA 4ijaA 4ijnA 4ikdA
4iknA 4ikvA 4il7A 4illA 4iloB 4in0A 4inaB 4indS 4inoB 4inwA
4inzB 4io2B 4ipiA 4ipsA 4ipuA 4iqbA 4irfA 4irvD 4iujA 4iulB
4iupA 4iusA 4iuwA 4ivnA 4iwbB 4ix3B 4iyaA 4iymH 4izhA 4j0dA
4j0wA 4j1oA 4j1vA 4j27B 4j2gB 4j32A 4j32B 4j37A 4j39A 4j42B
4j4aH 4j4hC 4j5rB 4j6oA 4j73A 4j7bF 4j7hB 4j7nA 4j7oA 4j7qA
4j8cA 4j8sA 4j9yB 4jaqC 4jbdA 4jbeA 4jbuA 4jccA 4jcqA 4jd0A
4jdeB 4jdnB 4jdxE 4je3A 4je3B 4jejA 4jemB 4jf8A 4jg2A 4jg9B
4jgiA 4jglA 4jgpB 4jgwB 4jhcA 4jhnD 4jhtA 4jifA 4jifB 4jimA
4jiuA 4jj0A 4jj9B 4jk8B 4jleA 4jmpA 4jn7A 4jndA 4jnhA 4jnuD
4jnzA 4jo0A 4jo7A 4jobA 4joiA 4joxA 4jp0A 4jp6A 4jpnB 4jqfA
4jqpA 4jquB 4jrlA 4js0B 4js1A 4jtmA 4jvcA 4jw2A 4jwjB 4jxrA
4jykB 4jz5A 4jzpB 4k02A 4k0dB 4k0nA 4k12A 4k12B 4k1cB 4k1pF
4k22B 4k2jB 4k36B 4k3zA 4k6nA 4k70B 4k7bA 4k7jB 4k82A 4k84B
4k8wA 4k90A 4k90B 4k94C 4k9zA 4kalA 4kcfA 4kddA 4kdrA 4kdwA
4ke2A 4kefA 4kemA 4kf9A 4kfuC 4kg0A 4kgdB 4kgiC 4khbC 4kiaA
4kjmB 4kk7A 4kkrB 4kkuC 4kkzC 4kl0A 4km6A 4kmcA 4kn8A 4knaB
4knkB 4kopC 4kp1A 4kp3D 4kppA 4kq7B 4kq9A 4kqdB 4kqiA 4kqpA
4krdB 4krgB 4krrA 4kruA 4ks9B 4ksnD 4kt3A 4kt3B 4kt6B 4kt6C
4ktbB 4ktwB 4ku0B 4ku0D 4kv2B 4kv7A 4kvgD 4kvxA 4kwdC 4kxqB
4kyqA 4l0cE 4l0kC 4l0nB 4l0rB 4l2hA 4l2iA 4l2iB 4l2wA 4l4eA
4l4qB 4l57A 4l5eA 4l5gB 4l6uB 4l77B 4l8aA 4l8hB 4l8iA 4l9aA
4l9bA 4l9eA 4l9hA 4l9oB 4l9pA 4l9pB 4l9uB 4la2B 4lanA 4lb8A
4ld1A 4ld6A 4ldvA 4le3C 4le7A 4lebA 4lf0A 4lfhG 4lflC 4lfuA
4lg8A 4lgjA 4lh6A 4lh9A 4lhfA 4litA 4lj9A 4ljiB 4ljrA 4ljsA
4lksA 4lldB 4lleB 4lloA 4lmgB 4lmoC 4lmyB 4lo6B 4loaA 4lonB
4looB 4lowB 4loxA 4lp8A 4lpqA 4lpsA 4lrlC 4lrtC 4lrtD 4lrvH
4lsdD 4lttA 4ltyA 4luaA 4lukA 4lunU 4lupC 4lv5A 4lv5B 4lvfB
4lviA 4lvnP 4lvpA 4lwsA 4lwsB 4lwuA 4lx2A 4lx2B 4lxqB 4lxrJ
4ly1C 4ly4A 4lyaA 4lypA 4lyyB 4lzkC 4lzxB 4m0nA 4m0qA 4m0wA
4m1aB 4m1bA 4m1gA 4m1uA 4m1xD 4m23A 4m2mA 4m37A 4m3lD 4m3oB

4m3pD 4m44E 4m51A 4m5bA 4m5dB 4m5rA 4m68A 4m6bF 4m6tA 4m70J
4m73A 4m7oA 4m7tA 4m82A 4m83A 4m85B 4m8aA 4m8dL 4m91A 4m9kA
4maaA 4maiA 4makB 4mamB 4maqB 4maxC 4mb0A 4mb7A 4mbyF 4mc3A
4mcjH 4mcoB 4mcwA 4mdaA 4me2A 4meaA 4mejC 4mesA 4mf5A 4mfiA
4mfkA 4mgqA 4mhlA 4mi7A 4mi8B 4mijA 4miyA 4mj2B 4mjdB 4mjkB
4mk6A 4mkoB 4mkxA 4ml1C 4ml9B 4mloA 4mm2B 4mmsC 4mn4C 4mn5B
4mncA 4mnnA 4mnoA 4mo0A 4mo1B 4mp8A 4mptA 4mr0A 4ms4B 4mswC
4msxA 4mt4A 4mt8A 4mtmA 4mtuA 4muoB 4muqA 4muvA 4mv4A 4mveA
4mvtC 4mydC 4mykA 4mypB 4myzB 4mz7A 4mzaB 4mzcA 4mzjA 4mzvA
4mzyA 4n01A 4n02A 4n06A 4n0hA 4n0hF 4n0lA 4n0nA 4n0pF 4n0tA
4n13A 4n1iA 4n2kA 4n2pD 4n2xG 4n30A 4n3aB 4n3xC 4n3yC 4n49A
4n4jA 4n4uA 4n5hX 4n5uA 4n6kA 4n6qA 4n77A 4n7cA 4n7fB 4n7wA
4n8nD 4n9wA 4na1B 4narA 4nawD 4nawK 4nb5C 4nbpA 4nbxA 4nc7B
4ncbB 4ncdA 4ndhA 4ndoA 4ndsB 4ne2A 4ne3A 4nesA 4nf0B 4nf1G
4nfaA 4nfuA 4nfuB 4ng0B 4ngdA 4nhbB 4nheA 4ni6A 4nj6B 4njcC
4njhB 4nkbA 4nkpD 4nl9A 4nmwA 4nmxA 4nmyB 4nn2B 4nn5A 4nn5B
4nnoA 4nnqB 4noaA 4nobA 4nocB 4noeC 4nogA 4nooB 4npdA 4nplA
4nq0A 4nqiC 4nqjA 4nqwA 4nqwB 4nrhD 4ns5A 4nsvA 4nsxA 4ntdA
4ntqA 4ntqB 4nutB 4nuuA 4nuxA 4nv0B 4nv4A 4nwbA 4nwyA 4nxyA
4nyhC 4nyqA 4nzgC 4nzrM 4nzvA 4o00A 4o06A 4o0aA 4o0cA 4o1eA
4o1rA 4o2hA 4o32A 4o3vB 4o4fA 4o5fA 4o5jA 4o5vA 4o65A 4o66D
4o6kC 4o6uA 4o6yB 4o7hA 4o7jB 4o7kA 4o87B 4o8sA 4o8vB 4o8wD
4o9dA 4o9lA 4oa3A 4obxB 4ociA 4ocvA 4od6A 4od8A 4od8C 4oe9A
4of6A 4ofaA 4ofkB 4oh7A 4oh8B 4ohjA 4ohxA 4oi3A 4oi4A 4oieA
4oihB 4oj8C 4ojxA 4okeB 4okoC 4okzB 4olsB 4oltB 4om8B 4ombA
4omfB 4omfG 4omgB 4onrA 4oo4B 4oobA 4oogB 4opcA 4oq9N 4oqpA
4oqvA 4oseA 4osnA 4otmB 4otnA 4ou0A 4ou6B 4ou9C 4ouhC 4oujB
4ounA 4ousA 4ov4A 4ov8A 4ovkA 4ow5A 4owfA 4owtA 4owtB 4owtC
4ox0A 4oxiA 4oxxA 4oydB 4oyhD 4ozkA 4ozuA 4ozwA 4p04B 4p09A
4p0gA 4p0tB 4p17B 4p1mB 4p1nA 4p1zD 4p27A 4p29A 4p3aB 4p3fA
4p3hA 4p3vA 4p5eB 4p5iA 4p5nB 4p5xA 4p6bB 4p6qA 4p79A 4p7cA
4p7oA 4p82A 4p8iB 4p8nA 4p9iA 4pasA 4pauA 4pawA 4pbzB 4pd6A
4pdnA 4pdxA 4pedA 4peoA 4peuA 4pf3A 4pgrA 4ph2B 4ph8A 4phjB
4phqC 4phrA 4pi8A 4pibJ 4pivB 4pj2B 4pj2D 4pk9A 4pkeA 4pkfB
4pkmA 4plzA 4pm4B 4pmkB 4pmoA 4pmxA 4pneB 4pnoA 4po6A 4po6B
4ponB 4pp4A 4pq9A 4pqdA 4pqgB 4pqhA 4pqqA 4pqzA 4psfB 4psrB
4psuA 4pt1B 4pt7D 4ptzC 4puiA 4pvkA 4pw2A 4pwsA 4pwwA 4pwyA
4pxeA 4pxwB 4py3C 4py9A 4pz0A 4pz1A 4pz3A 4q1tB 4q25B 4q29A
4q2lA 4q2sA 4q2uC 4q3kA 4q3oC 4q4gX 4q4w1 4q4w2 4q4w4 4q5eA
4q5wB 4q62A 4q63A 4q6vA 4q7aD 4q7eA 4q7oA 4q7qA 4q86D 4q8gB
4qa8A 4qamB 4qasB 4qboA 4qbuA 4qc6A 4qdcA 4qdgB 4qdjA 4qdnA
4qekA 4qflA 4qfuI 4qgoA 4qgsA 4qhqA 4qicD 4qitA 4qjvA 4qjvD
4qkwC 4ql0A 4qlaB 4qlpA 4qlpB 4qm6B 4qm9B 4qmaA 4qmhA 4qn8C
4qndA 4qo5A 4qosA 4qp5B 4qpnA 4qptA 4qpwA 4qq0A 4qq6A 4qrhD

4qtcA 4qtjA 4qtpC 4qtqA 4qusA 4qxaB 4qxlA 4qxzB 4r0sB 4r12A
4r17F 4r1dA 4r1dB 4r1qE 4r29C 4r2fA 4r2xB 4r2yC 4r30B 4r3nB
4r3qA 4r42C 4r4mB 4r4xA 4r52A 4r5rB 4r6hA 4r6yA 4r75A 4r78A
4r7qA 4r7rA 4r80A 4r81A 4r8hA 4r9fA 4r9iA 4r9pA 4rajA 4raxA
4rayA 4rbrB 4rbxA 4rd4A 4rd8B 4recA 4reiA 4rekA 4reoA 4repA
4rfqA 4rfuC 4rg1A 4rgdA 4rgiA 4rgpB 4rh0A 4rhjF 4rhpB 4rhsA
4rhzA 4rhzB 4ri2A 4ri5A 4ri6B 4rjwA 4rk2B 4rk6B 4rkhC 4rksB
4rl3B 4rl8D 4rlcA 4rleA 4rlzB 4rmlA 4rmmA 4rmxC 4rocA 4rp3A
4rp9A 4rpmA 4rptB 4rriA 4rs7D 4rscB 4rt1C 4rthA 4ru0B 4ru1F
4ru3A 4ru4F 4ruqB 4ruwA 4rvcA 4rvqA 4rw0B 4rwuA 4rxiA 4rxjA
4rxtA 4rxvA 4ry1B 4ry8A 4ry9B 4ryoA 4rz4H 4rz9A 4rzlB 4s1aA
4s1bA 4s1hA 4s28A 4s2rP 4s36A 4s3iB 4s3oC 4sgbI 4tjvA 4tkbA
4tkcA 4tkxL 4tl6A 4tm7A 4tmeB 4tmxA 4tpnA 4tpsB 4tpsC 4tq1A
4tq1B 4tq2A 4tq3A 4tqgA 4tqxA 4tr3A 4tr6B 4trhA 4trkA 4trtA
4ts4A 4tsdB 4ttwA 4tvsB 4tvvA 4tw5A 4tx3B 4tx5B 4txdA 4txrA
4txrC 4txwA 4tyzB 4tz1A 4tzmA 4u04A 4u0oB 4u12A 4u19B 4u1eB
4u1eG 4u1eI 4u1fA 4u3vA 4u4cB 4u4eA 4u4hA 4u4vA 4u5hE 4u5rA
4u5wC 4u63A 4u77A 4u7iA 4u89A 4u8fB 4u8pC 4u98A 4u9bA 4u9hL
4u9hS 4u9nA 4u9oA 4u9pC 4u9uA 4u9vB 4ua6A 4uafE 4uasA 4uc1C
4uc8A 4uciA 4ud4A 4udeA 4udkE 4udsA 4udxX 4ue0B 4ue8A 4ue8B
4ue9B 4uecB 4uejB 4ueyB 4uf0A 4uf1A 4uf7A 4ufqB 4uhcA 4uhoA
4uhqA 4uhtB 4uhvB 4uiqA 4uj7A 4um2A 4un1C 4un2B 4unuA 4uobA
4uonA 4uosA 4upuB 4uqwB 4uqxA 4uqzB 4us5A 4uskA 4uslD 4usoD
4usrA 4ut1A 4utuB 4uu3B 4uulB 4uuuA 4uuyA 4uvjA 4uvqA 4uw9A
4uwmB 4uwwA 4uwxA 4uybA 4uyiA 4uyrA 4uz1A 4uz8A 4uzsC 4uzyA
4uzzA 4uzzB 4v0hC 4v0pA 4v0wD 4v12A 4v17B 4v1gA 4v1jA 4v1sB
4v24B 4v2bB 4v2oA 4v3iA 4w4kB 4w4kC 4w4lC 4w5xA 4w64A 4w66A
4w6yA 4w78C 4w78H 4w7gA 4w7lB 4w7wA 4w8bA 4w8hA 4w8kB 4w8pA
4w8qA 4w9uB 4w9wA 4w9zA 4wa0A 4waiD 4watA 4wbdA 4wbeB 4wbjB
4wbtC 4wbyA 4wcjA 4wcxA 4wd1A 4wd8C 4wdcA 4we2A 4webE 4weeA
4wfcD 4wfcE 4wfqA 4wftC 4wfwA 4wh5A 4wh9A 4wheA 4whiA 4whnC
4wi1A 4wiaA 4wiqA 4wj1A 4wjiA 4wjoB 4wjtA 4wk0A 4wkaA 4wksA
4wksC 4wkzA 4wkzB 4wmaA 4wmlA 4wmyB 4wn5A 4wndB 4wnnD 4wolC
4wp6A 4wp9A 4wpgA 4wpkA 4wpmA 4wpyA 4wqdA 4wqkA 4wsfA 4wt3A
4wtpA 4wu0A 4wuiA 4wv4A 4wvaB 4ww7A 4ww7B 4wwfA 4wx0A 4wxaC
4wxmE 4wy4A 4wy4B 4wy4C 4wy4D 4wy9A 4wyhA 4wz4A 4wzrB 4wzxA
4x00B 4x1tA 4x1zB 4x28A 4x2cB 4x2hA 4x2hB 4x2pA 4x2rA 4x3iA
4x3nB 4x3xA 4x4wA 4x5mC 4x5pA 4x7gA 4x7rA 4x84B 4x86A 4x86B
4x8kB 4x8qA 4x8yB 4x90D 4x9fA 4x9kA 4x9tA 4x9xA 4x9zA 4xa9C
4xabA 4xalA 4xaxA 4xb6D 4xbaB 4xcbC 4xcpA 4xdnA 4xdnB 4xduA
4xdxA 4xe7A 4xeaA 4xekA 4xemA 4xezA 4xfjA 4xfkA 4xfmA 4xglA
4xgoB 4xguD 4xgvD 4xhtB 4xinA 4xj5A 4xjyA 4xkzA 4xlgA 4xltA
4xlyA 4xlzB 4xmrB 4xnhC 4xo1A 4xoiB 4xomB 4xotA 4xplA 4xpmA
4xpmB 4xpxA 4xpzA 4xqaB 4xqcA 4xr9A 4xrfA 4xrmB 4xrwA 4xslB

4xt6A 4xtbA 4xtrG 4xu6A 4xukA 4xuoB 4xurC 4xuvB 4xuwB 4xvxA
4xwjA 4xwwA 4xxfA 4xxhB 4xxiB 4xxuB 4xxwA 4xxxA 4xy5A 4xypA
4xywA 4xz6A 4xz7A 4xzaA 4xzfA 4xzrA 4xzzB 4y0bB 4y0hA 4y0lA
4y1bA 4y1wA 4y2fA 4y5jA 4y60C 4y68A 4y6wA 4y6xB 4y7dB 4y7lA
4y7sA 4y7uA 4y88A 4y99B 4y9iA 4y9tA 4y9vA 4yaaA 4yarA 4yb8B
4ybaB 4ybgA 4ybrA 4yc7B 4ycaB 4ycbB 4ycsE 4yd8A 4yddB 4ydrB
4ye7A 4yepA 4yf1A 4yfbE 4yg0A 4ygbD 4yh8A 4yh8B 4yhbA 4yhcB
4yhsA 4yhvA 4yi0A 4yiiA 4yj1A 4yj6A 4yk2A 4ykiB 4yl8A 4yl8B
4ylmX 4ymrB 4yn3B 4ynhB 4ynxA 4yomA 4yorA 4yp6C 4ypcA 4ypnA
4ypoB 4yrdB 4ys4A 4ysiA 4yslA 4yt2A 4ytbA 4ytdA 4ytkA 4ytlA
4ytoB 4ytwC 4ytwD 4yu8A 4yucA 4yudA 4yv4H 4yvoA 4yvqA 4ywaD
4ywfA 4ywkA 4ywtB 4ywzA 4yx6A 4yx7A 4yxpA 4yy2A 4yycA 4yz0A
4yz6A 4yzeC 4yzgA 4yzkA 4yzoC 4yzzA 4z04A 4z0gA 4z0vA 4z0yB
4z24B 4z29B 4z39B 4z3gA 4z3tB 4z4aA 4z54A 4z67A 4z6mB 4z7eB
4z7xA 4z80B 4z80C 4z8eC 4z8tA 4z8tB 4z8wA 4z9dB 4z9hA 4z9pA
4za1A 4za6B 4za9A 4zasE 4zavA 4zb3A 4zbgA 4zbhA 4zboD 4zbwA
4zc3A 4zcdA 4zceB 4zcnA 4zcrA 4zd6F 4zdjA 4zdsA 4zdtA 4zdtD
4ze8D 4zevB 4zf7A 4zflJ 4zfoF 4zfvB 4zgpA 4zh0A 4zh5B 4zhbA
4zhwA 4zi5B 4zieA 4zilB 4zj7B 4zj9A 4zjhA 4zjuA 4zk1F 4zkdA
4zkjB 4zkqA 4zkyB 4zl4A 4zldA 4zlhA 4zmiA 4zmkA 4znkA 4zo2B
4zoqG 4zosD 4zotA 4zoxB 4zoyA 4zp0A 4zpcA 4zpxB 4zqaA 4zquA
4zqxA 4zr7C 4zr8B 4zrlB 4zs9B 4zsfA 4zsiB 4zsxA 4zuaA 4zurB
4zv0A 4zv0B 4zv5B 4zvcA 4zvfA 4zw9A 4zwnD 4zwoA 4zwqG 4zx2A
4zy8B 4zy9B 4zylB 4zzfA 4zzzB 5a07B 5a0jA 5a0lB 5a0nA 5a0rA
5a0yF 5a10A 5a12D 5a1iA 5a1nA 5a1nB 5a1qB 5a1sC 5a2vA 5a35A
5a3aA 5a4aA 5a61A 5a62A 5a67A 5a6sB 5a6wB 5a6wC 5a71A 5a7gA
5a7vA 5a7zA 5a8cA 5a8jA 5a95B 5a96A 5a9cA 5a9dA 5a9hA 5a9tA
5ab4B 5absA 5abvB 5abxB 5ae0A 5aegB 5af0D 5af3B 5afdA 5afoA
5afyL 5ag8B 5agdB 5agiA 5agrA 5ah1A 5ahkA 5aigB 5aimA 5aizA
5ajgA 5ajjA 5ajoA 5ajsC 5al6A 5am2A 5ambA 5amhA 5amtA 5an5J
5an6C 5anpB 5anrC 5anzA 5ao9A 5aogA 5aooA 5aotA 5aozA 5apgB
5aq0A 5aq5E 5aqcA 5aqmB 5aunA 5awsA 5ax6A 5ax7A 5ay6B 5ayqB
5aysC 5ayvB 5azbA 5azpA 5azwA 5azxB 5b08A 5b0hB 5b0nB 5b0uB
5b19B 5b1qB 5b1rA 5b3dB 5b3fA 5b3pA 5b3tA 5b46B 5b4bB 5b4sB
5b4zA 5b52B 5b5iA 5b5lA 5b5qB 5b5zA 5b66Y 5b6cA 5b6dA 5b6qA
5b7eA 5b7fA 5b7hA 5b7wA 5b7yB 5b82A 5b8dA 5bjxA 5bk6B 5bmoC
5bmqA 5bnxC 5bnzA 5bo7B 5bobC 5boiA 5bopD 5bowA 5bp3A 5bp8A
5bp9A 5bpdA 5bpkA 5bpkC 5bpxA 5bpzA 5bq5B 5br4B 5brhB 5brlA
5bs1A 5bseA 5bt9C 5btoA 5btyA 5bu3B 5bupA 5buwA 5bvaA 5bvlA
5bw0B 5bwdC 5bwjC 5bx1A 5bxaA 5bxrB 5by4A 5by5A 5by7D 5by8A
5by8B 5bz0A 5bz3A 5bzaC 5c05B 5c0pA 5c0yB 5c12A 5c17A 5c1fB
5c22C 5c2iB 5c2mA 5c2uA 5c30A 5c40B 5c4yB 5c50A 5c50B 5c55A
5c5cA 5c5dD 5c5gA 5c5hA 5c5zB 5c6dA 5c6hP 5c6kB 5c6sA 5c79B
5c8aC 5c8gB 5c8hA 5c8zB 5c90E 5c98A 5c9fD 5c9lA 5c9oA 5ca8A

5cajA 5ccbB 5ccfA 5cd2A 5cdkA 5cdvA 5ce6A 5ce7A 5cecA 5cecB
5cegB 5cegC 5cemA 5cesA 5cfjA 5cgqA 5cgqB 5cgzB 5chhA 5chlA
5chpA 5chsA 5cjzA 5cklA 5ckwA 5cl2B 5cm2Z 5cmlB 5cnwA 5co8A
5cofA 5cosC 5cotA 5cowA 5cpcB 5cphB 5cqgB 5cr4B 5cr6D 5cr9A
5crbB 5cs0A 5csrB 5ctaA 5ctmA 5cttB 5cu7A 5cukA 5cuoB 5cv0A
5cvaE 5cvdB 5cvwA 5cwgA 5cwhA 5cwiA 5cwqA 5cwtB 5cwwB 5cwwC
5cx2A 5cx7I 5cx8B 5cxxB 5cyaA 5cybA 5cyvA 5cywB 5czcA 5czgA
5czlA 5czyA 5d01B 5d16B 5d18A 5d1mB 5d1rA 5d22A 5d23A 5d2eA
5d2kA 5d3aA 5d3kA 5d3qB 5d3xB 5d4fA 5d4vB 5d50B 5d5kB 5d5nA
5d5yB 5d6eA 5d6hA 5d6hB 5d7uB 5d8mA 5d8vA 5d9rA 5dblA 5dclB
5dcuB 5ddwD 5de3A 5decD 5deqA 5dfkA 5dfyA 5dggA 5dgjA 5dgqB
5dh0B 5dhdA 5di0B 5dicA 5dilB 5djeB 5djhA 5djoB 5djtA 5dlbA
5dleD 5dlkD 5dm2A 5dmaA 5dmdB 5dmmA 5dn2G 5dn7A 5dnkB 5docB
5dofB 5dokB 5dp2A 5dpoA 5dqlD 5dtcA 5du9A 5dutA 5dviA 5dvwB
5dwaB 5dwdA 5dwzB 5dxlA 5dyqB 5dz2B 5dz8A 5dzeA 5dzoA 5e04B
5e0yA 5e10A 5e1lA 5e1vA 5e1wA 5e1yA 5e3bA 5e3eB 5e3eE 5e3iB
5e3qA 5e4bB 5e4gA 5e56A 5e57A 5e6gA 5e6qA 5e6xA 5e71A 5e75A
5e7hA 5e8zA 5e9nA 5e9pA 5eb8A 5ebaA 5ec0A 5ecdA 5eckD 5ed9B
5edfA 5edlA 5efmA 5efrA 5eftH 5efvB 5efzF 5egfA 5eh1A 5ehaA
5eikA 5ej0A 5ej8H 5ejdF 5ejlA 5ejrA 5ejyA 5ek5A 5el3A 5el9A
5elbC 5elkA 5emiA 5emxB 5en8B 5enqB 5ep6A 5ep9B 5epeA 5epwA
5eqjA 5eqnB 5eqzA 5er9B 5ereA 5ermB 5erqA 5et1B 5et3A 5eu0B
5eurC 5evhA 5ew0A 5ewoA 5ewpB 5ewuA 5ewyA 5ex2A 5ex8A 5exeA
5exeC 5eynA 5ez7A 5ezqA 5ezuB 5f18A 5f1bB 5f22A 5f22B 5f29B
5f2kA 5f2tA 5f30B 5f47A 5f4cB 5f4wA 5f5oD 5f5sB 5f6lJ 5f6rB
5f74B 5f7rE 5f7vA 5f9pB 5f9qA 5fa1B 5fa8A 5fafA 5fb4A 5fbfA
5fbyB 5fcgC 5fcmB 5fcnB 5fd9A 5fewA 5ffdA 5ffgB 5ffpA 5ffqA
5ffxC 5fg0A 5fglA 5fgpA 5fh7A 5fhkD 5fi3A 5fiaB 5fidA 5fiiB
5fjdC 5fjzC 5flgB 5flhB 5flwA 5flyB 5fmdA 5fmoL 5fmoS 5fmrA
5fmuC 5fn7B 5fn8B 5fnoA 5fnpB 5foeA 5fpwA 5fq4B 5fqeB 5fr7B
5freA 5fryB 5fs4B 5fs8A 5fshB 5fsvA 5ft0B 5ftbA 5fu5A 5fukA
5fv8A 5fvdD 5fvjB 5fvkA 5fwhA 5fwzA 5fxqA 5fxsA 5fyaB 5fydB
5fypB 5fzpB 5fzsA 5g1lA 5g1sN 5g1wE 5g23B 5g25A 5g28A 5g2cA
5g2dA 5g2pD 5g2uA 5g38A 5g3qB 5g3tB 5g3xA 5g4iA 5g4kA 5g4zA
5g51A 5g5cA 5g5oF 5ggbA 5ggnA 5gheA 5gi7A 5gizA 5gj7A 5gjhA
5gjkB 5gk1D 5gk9B 5gkeA 5gkvA 5gkxB 5gl6B 5gmcD 5gmdA 5gmtB
5gn1D 5gn2D 5gnaB 5gnfB 5gngB 5gooA 5goxB 5gpoA 5gpyA 5gpyB
5gq1E 5gqfB 5grqA 5gs7A 5gt5A 5gtqA 5gtuA 5gtuB 5gudF 5gufA
5guqC 5gv0A 5gv3B 5gv8A 5gvaA 5gvcB 5gvdA 5gvvF 5gw9C 5gwnA
5gxeA 5gxwB 5gyqA 5gyyA 5gz3B 5gzaA 5gzkA 5gztA 5gzyA 5h02A
5h06B 5h0jA 5h0mA 5h0qA 5h1xB 5h28A 5h29A 5h2fO 5h3jA 5h3jB
5h3kA 5h3vA 5h3xA 5h4sA 5h5aB 5h5fA 5h5pA 5h62B 5h66C 5h6iB
5h6tA 5h6xB 5h6zA 5h72F 5h7eA 5h7kA 5h9iA 5h9nA 5halA 5hamB
5hawB 5haxB 5hazA 5hb6A 5hb7A 5hbpA 5hcnA 5hd9A 5hdaC 5hdkD

5hdmB 5hdwA 5he9A 5he9E 5heaB 5heeA 5hfiA 5hgjB 5hgzA 5hheD
5hhjA 5hi8B 5hiuD 5hj9A 5hjfB 5hjmA 5hjqA 5hk3B 5hkqA 5hkqI
5hl3A 5hl8A 5hljA 5hloB 5hmlA 5hmqE 5hnoC 5hnvA 5hoeD 5hokC
5honB 5hopB 5hpfC 5hqhA 5hraB 5hrbA 5hsfA 5hsiB 5hstA 5hsxA
5ht7B 5htlA 5htxA 5hu7A 5hubA 5husA 5hvxA 5hwaA 5hweA 5hwhA
5hwkA 5hwnB 5hx0B 5hxkB 5hxyF 5hyzA 5hzdA 5i0cA 5i0qA 5i0yA
5i1uB 5i20D 5i2cC 5i2hB 5i34B 5i39A 5i3eB 5i41B 5i45A 5i47B
5i4aC 5i5dB 5i5mB 5i5nB 5i62A 5i6kA 5i6rA 5i7iA 5i8jB 5i8tA
5i90A 5i95A 5i9jA 5iaaC 5ib0E 5ibvA 5ibwC 5ibzD 5ic7A 5icuA
5idbA 5idhA 5idvA 5iffB 5ig0A 5ig6A 5ig8B 5igiA 5iheA 5ihfB
5ihwA 5ii6A 5ii8A 5iibB 5ijaA 5ijjA 5ik4A 5ikjA 5ikjB 5il6B
5ilbA 5imaA 5imuA 5in3B 5in4B 5inbB 5inrC 5io9B 5iojB 5ipxA
5ipyB 5iqjC 5ir2A 5ir4A 5isuA 5isvA 5it0A 5it3B 5itmD 5itqA
5iu1A 5iucB 5iufB 5iw9B 5iwzA 5ix8A 5ixgB 5ixhB 5ixpA 5iyzA
5izaA 5izeA 5izsD 5iztA 5izwB 5j03A 5j09H 5j1bB 5j1gB 5j1jA
5j1kA 5j1lD 5j1nA 5j1sA 5j1sB 5j2lA 5j2yB 5j3tA 5j3tB 5j3tC
5j3uA 5j41B 5j49B 5j4aB 5j4aC 5j4fA 5j4iA 5j4lA 5j4mA 5j4oA
5j4uA 5j53A 5j5lA 5j6qA 5j6yA 5j71A 5j7eD 5j7mB 5j7rB 5j80A
5j90A 5j93A 5j9iG 5ja9C 5jawH 5jazB 5jblA 5jbnB 5jbrB 5jbxA
5jcaS 5jcyB 5jdaA 5jdjL 5jdkA 5je2B 5jedA 5jeeA 5jelA 5jenC
5jeqA 5jffD 5jg9B 5jgeE 5jgfA 5jgkB 5jiaE 5jicA 5jieD 5jigA
5jipB 5jj2A 5jjaB 5jjoA 5jjxA 5jk0D 5jk2D 5jkiA 5jkqB 5jlbA
5jm8F 5jmbB 5jnmA 5jnoA 5jnoB 5jnpB 5jo8A 5jp2E 5jphC 5jpoD
5jqyA 5jrbF 5jrcC 5jrhA 5jrjA 5jrtA 5jryA 5jufA 5jugA 5juhA
5jviE 5jvkA 5jw9A 5jwoB 5jxcC 5jxmA 5jxzA 5jzeD 5jzuB 5k08A
5k0pJ 5k1pA 5k21C 5k2lA 5k2xA 5k34A 5k3qA 5k3xA 5k4bA 5k4xA
5k5dC 5k5zD 5k62A 5k7fA 5k87A 5k8cA 5k8pD 5k8sA 5k91A 5kagK
5karA 5kayB 5kc1B 5kc8A 5kciA 5kd5A 5kddA 5kdoB 5kdoG 5kdsA
5kdzA 5ke1B 5kecA 5kefB 5kewC 5kewD 5kf9A 5kfzA 5kiaA 5kkoF
5kkpA 5kkuC 5klaA 5kleA 5klhB 5knhI 5knkB 5ko5A 5ko9A 5koeB
5koxA 5kprA 5ktcA 5ktkA 5ktnA 5ku5A 5kutA 5kuxA 5kvbA 5kvmA
5kvrA 5kvsB 5kwbA 5kweD 5kwnA 5kx4B 5kxhA 5ky0B 5ky4B 5kzaA
5kzzA 5l04B 5l09B 5l0lB 5l0rB 5l0vA 5l16A 5l1aA 5l20A 5l2lE
5l37C 5l44B 5l4lA 5l71A 5l73A 5l74A 5l77A 5l7jA 5l7zA 5l87A
5l8sC 5l8xB 5l9zA 5l9zB 5lacA 5lalB 5lb3E 5lb4A 5lb7B 5lbdB
5lc9D 5lcyC 5ld9B 5lddB 5lddD 5ldvA 5lf2B 5lf9A 5lfnA 5lgdA
5lgdB 5lggA 5lhwA 5lhxB 5lj8B 5ljmA 5ljwB 5ljxA 5lkqA 5lljB
5lltB 5lmgB 5lndB 5lniA 5lnnA 5lnrD 5lolA 5loqC 5lp9A 5lpaA
5lpiB 5lq6A 5lrrD 5lrtA 5ls7B 5lsiD 5lsiE 5lslG 5lt5B 5lteA
5ltgA 5lu5D 5lunD 5lusF 5lw0B 5lw3A 5lw5A 5lx8A 5lx9A 5lxeB
5lxfB 5lxnE 5lxuA 5lxvA 5lxvB 5lxxB 5lxzB 5ly0B 5ly3A 5ly8A
5ly9A 5lypA 5lzkB 5lznA 5m0iB 5m0nA 5m0yB 5m10A 5m12A 5m17A
5m1mA 5m1pB 5m1xA 5m26A 5m26H 5m2oB 5m2pA 5m2yB 5m33B 5m3iB
5m45C 5m4yE 5m5eB 5m5eC 5m72A 5m72B 5m7dA 5m7yA 5m86D 5m89B

5m8cB 5m8hD 5m97A 5mabA 5malA 5maoA 5mawE 5mb2B 5mbxA 5mc1A
5mc7B 5mc9A 5mdiA 5mdtA 5mduA 5me4A 5me5B 5mebA 5mecA 5mf2B
5mfoD 5mgwA 5mgzA 5mhjA 5mixA 5mjhA 5mjrA 5mk9A 5mkcC 5ml3B
5mldE 5mltA 5mlzA 5mphA 5mptA 5mpwC 5mq8A 5mqhA 5mqiA 5mqpC
5mr1A 5mriA 5msmE 5msmF 5msnA 5mu4D 5mu5A 5mu9A 5mujA 5mulA
5munA 5muxE 5muzA 5mv0B 5mvwB 5mvwD 5mwbA 5mwnB 5mx2F 5mx2K
5mx2V 5mx9A 5my5A 5my7A 5n0lC 5n22B 5n2cB 5n2uC 5n3uA 5n3uB
5n40A 5n41A 5n5eT 5n5fF 5n5pC 5n6fA 5n76C 5n7eB 5n7sB 5n7zA
5n81B 5n8aX 5n9mA 5na1A 5naaA 5nakB 5ncgB 5ncjB 5nf2A 5nf4B
5ng9A 5ngdD 5nggA 5ngjA 5nglC 5ngnB 5ngoE 5ngwA 5nhmD 5nioA
5nirB 5nj9C 5nj9D 5njlA 5njoA 5nkmB 5nkmC 5nkqB 5nl9B 5nmnA
5nmoB 5nnyA 5no8B 5noaA 5nodA 5nohA 5nonC 5nopB 5npkB 5nptA
5npyA 5nqoA 5nqvB 5nrmA 5nsaA 5nsjB 5nt7A 5nt7D 5nthA 5nulA
5nusA 5nusB 5nuvA 5nv9A 5nvmD 5nwpB 5nxfA 5nxkC 5nykA 5nypA
5nzgA 5nzoB 5nzxA 5o01A 5o0tA 5o0uA 5o15A 5o1lB 5o29A 5o2dA
5o30C 5o33B 5o37A 5o3mD 5o58A 5o5sA 5o63B 5o65A 5o75A 5o8mD
5o99B 5o9eA 5o9eB 5o9mA 5oaqA 5oaqL 5oarC 5obtF 5od4A 5od6B
5odqC 5odqF 5odqH 5odqJ 5odtB 5oe3C 5oe8A 5oeiA 5oemA 5oh5A
5ohoB 5ohqA 5oi7B 5oixC 5oj7A 5ok8C 5ol4C 5ol9A 5olrC 5oluA
5omkA 5omtA 5oo7B 5oooB 5opbA 5opfA 5opgA 5opqC 5opzA 5oq3A
5oswA 5ouoA 5oupA 5ovoA 5ovuB 5oyjC 5p9vA 5pmxA 5suiA 5suzA
5sv2A 5sv6A 5svdB 5swcE 5swuB 5sy4B 5sy8O 5sybB 5syrB 5sytA
5szcA 5szhA 5t09A 5t0fB 5t11F 5t12A 5t2pB 5t2xB 5t39A 5t3bA
5t44B 5t51A 5t51B 5t59C 5t5iJ 5t6jA 5t6jB 5t76A 5t77A 5t7aA
5t7zA 5t86A 5t86I 5t87E 5t8cA 5t95A 5t9yC 5tabA 5tcbA 5tckB
5td6A 5tdaA 5tdeA 5tdyB 5tedB 5tf3A 5tfpA 5tfqA 5tgfB 5tgnA
5tgqA 5thxA 5tifA 5tihA 5tipC 5tjaA 5tjjB 5tjzA 5tk8A 5tkwA
5tl4D 5tleA 5tnvA 5tohB 5tpiA 5tqbA 5tqbB 5tqiB 5trqA 5ts9D
5tsaA 5tscB 5tseA 5tsqA 5tt6A 5ttaA 5ttyA 5tucB 5tuhB 5tuiB
5tuuA 5tuuB 5tuxA 5tvoA 5tvoB 5tw9F 5twaA 5twbA 5txuA 5tz5A
5tzdA 5tzjA 5tzpA 5u0iA 5u19A 5u22A 5u2lA 5u2pA 5u32A 5u35B
5u3qB 5u4hB 5u4sB 5u4uA 5u55D 5u5gB 5u5nB 5u69A 5u75A 5u7aB
5u7fA 5u7wA 5u81A 5u96A 5u9jB 5u9nB 5ua5A 5uamB 5uaoC 5uayA
5uazA 5ub3A 5ubaA 5ubeA 5ubpA 5ubpB 5ubpC 5ubwA 5ucsB 5ucvB
5ud5B 5udnB 5udqF 5ue1B 5uebB 5uejA 5ufkA 5ufnA 5uftA 5ugrA
5ugwA 5uh7A 5ui9B 5ujcA 5ujdB 5ujuB 5ukhA 5ukiA 5ulmA 5um2A
5um7A 5umfB 5umhB 5umsA 5umvA 5un7A 5un7B 5uniA 5uniB 5uofA
5upbC 5uq6A 5uqdA 5uqjA 5uqzA 5usbA 5uswB 5ut3A 5utkA 5uuiA
5uvdA 5uvgA 5uw8B 5uwaB 5uwcG 5uwcI 5uwrD 5uwzA 5ux1C 5uxmA
5uz8A 5uzgB 5uzxB 5v01A 5v07Z 5v0zC 5v13B 5v1tA 5v1yB 5v2oF
5v2qA 5v37A 5v3nA 5v3nB 5v3sB 5v3wA 5v44A 5v50F 5v6fA 5v6gC
5v77B 5v7pA 5v87B 5v8kA 5v8sA 5v8wE 5v8wF 5vacA 5vb9A 5vbdA
5vbnA 5vcmA 5vegC 5vfaB 5vflL 5vfyB 5vfzA 5vg2U 5vg3B 5vgbA
5vgbB 5vglA 5vgtA 5vhgA 5viaA 5vipA 5vipB 5vitI 5vitR 5vj2B

5vjiA 5vjwA 5vkdA 5vkyA 5vliC 5vlqA 5vmdC 5vmnA 5vnyA 5vogA
5volE 5vpeD 5vpsA 5vqeA 5vr2B 5vrhA 5vrvE 5vsmB 5vsxA 5vt9D
5vtgA 5vugA 5vx1B 5vxeA 5vykC 5vz4B 5w0gA 5w15D 5w1eA 5w1uB
5w2fA 5w2iA 5w2lB 5w3xC 5w53B 5w56B 5w5cE 5w6yB 5w7bB 5w7bC
5w83A 5w83B 5w8eA 5w8oB 5w8qB 5w8xA 5w93B 5w95A 5w98A 5wanA
5wasA 5wasB 5wasC 5wbfA 5wc6M 5wcjA 5wd9A 5wddB 5we0F 5we0G
5we0K 5we8B 5weeB 5wf2A 5wfbA 5wggA 5wgiA 5whmA 5wi2A 5wjeA
5wk0A 5wk4B 5wl1A 5wmfC 5wn9A 5woqA 5wpsA 5wqjA 5wqvA 5wqwA
5wriA 5ws5T 5ws7A 5wsfA 5wstB 5wsvD 5wsyB 5wtqD 5wucA 5wujA
5ww9D 5wwdA 5wwoA 5wwwA 5wx9A 5wxmA 5wxmU 5wy4A 5wzjA 5wzqA
5x03B 5x14A 5x1eC 5x1eD 5x1eE 5x2eA 5x2nB 5x3dA 5x40A 5x42B
5x42C 5x4bA 5x4rA 5x4sA 5x4tA 5x57A 5x5mA 5x5vA 5x62A 5x6sB
5x6vF 5x6vG 5x7nA 5x86A 5x89A 5x9aA 5x9cB 5x9iB 5x9jB 5x9kB
5x9lA 5xa5A 5xa5B 5xaqA 5xauA 5xauB 5xauF 5xavB 5xb6H 5xbcA
5xbfA 5xbiB 5xbjA 5xbuA 5xccB 5xcyA 5xdcD 5xdhC 5xdtA 5xdzB
5xe7A 5xeaA 5xecA 5xevA 5xfmC 5xg2A 5xgaA 5xgcA 5xgtA 5xguB
5xhwA 5xisA 5xiuA 5xj1A 5xjgB 5xjgC 5xjnA 5xk6B 5xkrC 5xktA
5xkxA 5xljA 5xlsA 5xlyA 5xlyB 5xm5B 5xmzA 5xn9B 5xnhA 5xodB
5xojE 5xomB 5xp6A 5xpcB 5xqhA 5xsjL 5xspB 5xssB 5xtaC 5xunB
5xveA 5xvrA 5xvsA 5xvtA 5xw4B 5xxsB 5xzgA 5y11C 5y1fA 5y2sA
5y2wA 5y37A 5y3cA 5y4zA 5y59B 5y5aA 5y5aB 5y5sQ 5y78A 5y7yA
5y8eA 5y90A 5y96A 5y9eE 5y9gB 5y9qA 5y9wC 5yagB 5yalA 5yayB
5yckA 5ydeA 5yegA 5ygbA 5ygeB 5yghB 5yhyA 5yi7D 5yixB 5yjwA
5yk0A 5yowA 5yq5B 5yqjA 5yqwA 5ys3A 5yufA 5ywzA 5yxmA 5z0uA
5z11B 5z1gB 5z1gC 5z38D 5z42A 5z67A 5z8oA 5z9yA 5zb8C 5zbyA
5ze8A 5zhoA 5zhzA 6amgB 6ansD 6anwB 6anzA 6ao1C 6ao3A 6ao7A
6ao8A 6ao9A 6aonB 6aozC 6appB 6ar7A 6as4A 6atwA 6au8A 6au8C
6avxA 6ax7B 6az5A 6b0gE 6b29A 6b2vA 6b2yA 6b3aA 6b3pA 6b3xA
6b3yA 6b4aB 6b5kA 6b6uA 6b7pB 6b8wB 6b91A 6b9fA 6b9hA 6b9hB
6b9mA 6b9mD 6b9rC 6b9xA 6b9xB 6b9xC 6b9xD 6b9xE 6bc7A 6bcbA
6bevA 6bfnA 6bhcA 6bk0A 6bl5A 6blkD 6bniA 6bo0A 6bs9C 6bsyA
6bxoB 6bypA 6bzgA 6bzgB 6bzhC 6c2jA 6c2jB 6c34A 6c3cA 6c4mC
6c4vA 6c5bA 6c62C 6c6zB 6c8rB 6cb2A 6cbnA 6cd7A 6cd9A 6ck0B
6ckgA 6ckoA 6cktA 6cojA 6cojB 6cpyB 6cr0A 6cumA 6cuqB 6cwqA
6cy6A 6eh4D 6ehbC 6ehdA 6ehiB 6ehnA 6ei1A 6ei3A 6eioA 6ej2A
6ekbA 6ektA 6el8D 6elmA 6eluJ 6emgA 6emyA 6enoA 6ensA 6enxA
6eomA 6eozA 6epzD 6eqeA 6eroB 6es9A 6etlA 6eu6A 6euwA 6exaA
6expF 6ey0A 6ey4A 6ey6D 6eyuB 6f0pA 6f6pC 6f72A 6f8aB 6f8pA
6f9gA 6fdgB 6feaF 6fhoA 6fljA 6flkB 6fnuA 6ft2A 6fthA 6gajC
6rlxD 7a3hA 7odcA 8abpA

#### 5.2 The 495-protein test set for the contact map predictor

1bxoA 1d1gA 1eq2A 1f0cA 1f35A 1fkmA 1h32A 1h4xA 1j98A 1khxA
1l8wA 1lqvA 1mbaA 1nmoA 1pu6A 1qhdA 1qwoA 1ro7A 1u58A 1vclB
1vk1A 1vqoH 1vqoW 1ws6A 1xv5A 1yg9A 1yrkA 1z6mA 2aotA 2apjA
2cduA 2ci1A 2dg6A 2ghtA 2gigC 2hsjA 2hy5A 2ih2E 2iwrA 2iy9A
2j58A 2nqtA 2oizC 2oplA 2ot4A 2p39A 2ph0A 2pn6A 2qgqA 2r8rA
2v2fF 2v8fA 2vgaA 2vovA 2w15A 2w6kA 2w7zA 2wfiA 2xgvA 2ynmC
2ynmD 3a98C 3a9fA 3al9A 3bijA 3cetA 3cp3A 3cx5B 3cx5C 3cx5D
3d0wA 3dwcA 3e21A 3e2oA 3edoB 3eglA 3egnA 3eqzA 3f0hA 3f8kA
3f9sB 3fdeA 3ff1A 3fh1A 3fjsB 3fkaC 3floB 3fmyA 3fncA 3fsgA
3fz4A 3g14B 3g1pA 3g23A 3gemA 3giwA 3go9A 3gu3B 3gycA 3h6rA
3hduC 3hi2A 3i0zB 3i3wA 3ib5A 3igfA 3ip3A 3iruA 3ismC 3itqA
3iu6A 3iwfA 3jq0A 3jtxB 3ju7B 3juiA 3k1uA 3k6qC 3ke3A 3kkfA
3kq0A 3kxpA 3l0aA 3l9aX 3ledA 3li9A 3licA 3lm3A 3lmaD 3lvuA
3lxqA 3lxzA 3m4rA 3m84A 3mazA 3mc3A 3mczA 3mdpA 3mf7A 3mfiA
3mgbA 3mjfA 3mnmA 3mtqB 3mtwA 3n1fC 3n91A 3nedA 3njcA 3o5yA
3oopA 3oosA 3oouA 3oovA 3oqpA 3os4A 3p1gA 3pj0A 3pt8B 3pwxA
3qmlC 3qtaA 3qwmA 3rjvA 3rwxA 3sd2A 3sonA 3t6oA 3t6sA 3tc8A
3u0hA 3u3lC 3u52C 3u5sA 3ubyA 3uoaB 3urzA 3vygA 3wa2A 3zuzA
4akpA 4aukA 4ay9B 4bq6D 4bwcA 4c12A 4chgA 4citA 4ck4A 4d9iA
4dlqA 4dweA 4e29A 4e5vA 4eojB 4epsA 4exrA 4fxiA 4g4sP 4gemA
4ghnA 4gqbB 4gudA 4h0aA 4hd1A 4hesA 4hlbA 4hyzA 4i79A 4iumA
4iyjA 4jg5A 4k7cA 4kbxA 4kwaA 4l0jA 4l8nA 4lq8A 4lqxA 4lrzE
4ng2E 4nk6A 4nn5C 4nzkA 4o2tA 4oi4B 4opmA 4opwA 4p98A 4pbdA
4pdyA 4pioA 4pw0A 4pxyA 4q53A 4q7fA 4qblA 4qn1A 4qrkA 4qvsA
4r03A 4r7fA 4rv0B 4rv5A 4s0uC 4tqrA 4u3eB 4uabA 4v00A 4v4mE
4wlhA 4wriA 4wuvA 4wv4B 4x28C 4x9rA 4xp7A 4zi3C 5a0yB 5a3dA
5aooB 5b66U 5c33B 5cozA 5d78A 5e65B 5eqvA 5eqwE 5h9fA 5ht2A
5klpA 5kvgE 5l2xA 5l8eA 5le5I 5le5J 5le5L 5lsfC 5n0oA 5n2pA
5nb4A 5o8wB 5oc7D 5t1pA 5t5iC 5toqA 5u3aA 5udiA 5umpA 5uouA
5uukA 5uxxA 5v5hA 5v8dA 5vxvA 5w7pA 5wxxA 5xefA 5xjl2 5xyfA
5yf4A 5ylbA 5ylwA 5yrhA 5yxgA 5z08C 5z3eA 5z4aA 5zbtA 5zfkA
5zgnA 5ziqA 5zqhA 5zt3A 5zt8A 5zzuA 6a0nA 6a52A 6a5hA 6ad3A
6ae1A 6aikA 6akmA 6aqkA 6b7lA 6bdeA 6beaA 6bk6A 6br8A 6brpB
6bsuA 6byfA 6c0bA 6c6nA 6cj7A 6cokA 6d0hA 6d2sA 6d2yA 6d71B
6d9fA 6dchA 6dg6F 6dgmB 6dhpU 6dj8A 6dxoA 6e0oA 6e5fA 6e7rD
6ejiA 6eskA 6f45D 6f5zA 6f7dA 6fdfA 6fdlA 6ff1A 6fkgA 6ftqA
6fv5A 6fzzA 6g7wA 6g94A 6g9xA 6gbiA 6geuA 6gf6A 6ghoB 6gjeB
6greA 6grhC 6gs2B 6gv5A 6gy8A 6gz0A 6h48A 6hatA 6hd8B 6heiA
6hfzA 6hhmA 6hjxC 6hqzA 6i0iA 6i50A 6i5rA 6i9kA 6iahA 6idpA
6iffB 6iieA 6ij2D 6iluA 6imlA 6imvA 6iqtC 6istB 6iujA 6iy9A
6iz2A 6jc4D 6jcnA 6jkrF 6jl7A 6jliA 6ki2A 6km7B 6kmeA 6kplA
6ktyA 6l1oA 6l4pB 6l5tA 6lu2A 6m7zE 6mgcA 6micA 6mpzA 6mq7A
6mt7A 6mw7A 6mzpA 6n2cA 6n7mA 6n9hA 6nepA 6nibA 6njwA 6nkhD

6nl1A 6nlpA 6o1bA 6o35C 6o6jA 6o80A 6oghA 6ohiA 6oj1A 6ompA
6on1A 6ouxA 6oz1A 6ozuA 6p0fA 6pb3A 6pfnD 6prhA 6pt4B 6pv4B
6pwgA 6q67A 6q9cA 6qq4A 6qspB 6qwoA 6qycA 6qziA 6r5wA 6r7oA
6rimH 6rjxA 6rn5A 6rnkA 6rnqB 6s07A 6s0eA 6sg8B 6sizB 6sjqA
6sooA 6t3zA 6tjhA 6tpvB 6tv1A 6tv2D 6tx2A 6tzkA 6u8zA 6uloA
6uncD 6uofA 6uxuB 6vkzA 6vtvB 6vycB 6w08B 6w1tU 6w3dA 6wajA
6xvmB 6xyaB 6y3pA 6y7fA 7br2D
